# A laboratory module that explores RNA interference and codon optimization through fluorescence microscopy using *Caenorhabditis elegans*

**DOI:** 10.1101/2020.10.17.344069

**Authors:** Nicholas J. Palmisano, Maryam A. Azmi, Taylor N. Medwig-Kinney, Frances E. Q. Moore, Rumana Rahman, Wan Zhang, Rebecca C. Adikes, David Q. Matus

## Abstract

Scientific research experiences are beneficial to students allowing them to gain laboratory and problem-solving skills, as well as foundational research skills in a team-based setting. We designed a laboratory module to provide a guided research experience to stimulate curiosity, introduce students to experimental techniques, and provide students with foundational skills needed for higher levels of guided inquiry. In this laboratory module, students learn about RNA interference (RNAi) and codon optimization using the research organism *Caenorhabditis elegans* (*C. elegans*). Students are given the opportunity to perform a commonly used method of gene downregulation in *C. elegans* where they visualize gene depletion using fluorescence microscopy and quantify the efficacy of depletion using quantitative image analysis. The module presented here educates students on how to report their results and findings by generating publication quality figures and figure legends. The activities outlined exemplify ways by which students can improve their critical thinking, data interpretation, and technical skills, all of which are beneficial for future laboratory classes, independent inquiry-based research projects, and careers in the life sciences and beyond.

## SCIENTIFIC TEACHING CONTENT

### Learning Goals

#### General knowledge

- Gain experience working with *C. elegans*
- Understand the process of RNA interference and importance of codon optimization

#### Technical skills

- Develop mastery in fluorescence microscopy techniques and image analysis

#### Communication skills

- Enhance their writing skills

### Learning Objectives

Students will be able to:

#### General knowledge

- Demonstrate the ability to synchronize *C. elegans* nematodes and perform an RNAi experiment
- Describe what RNAi is and how it affects gene expression/activity
- Explain what codon usage means

#### Technical skills

- Acquire images using an epifluorescence microscope
- Calculate mean fluorescence intensity from acquired fluorescence micrographs
- Perform statistical tests to determine the significance of results

#### Communication skills

- Generate publication quality figures and figure legends
- Effectively formulate conclusions from data and logically present results

## INTRODUCTION

Inquiry-based learning is a form of active learning where students can gain problem solving skills that can help better facilitate inquisitive thinking while simultaneously allowing them to make unique discoveries about the natural world(1–3). In contrast to teacher-centered instruction, where facts are disseminated to students, inquiry-based learning encourages students to foster their own independent learning with the assistance of the instructor(1–3). In addition, inquiry-based learning puts emphasis on students developing scientific skills, such as making observations, developing hypotheses, analyzing data, and formulating conclusions(1–3).

Course-based Undergraduate Research Experiences (CUREs) are a form of inquiry-based learning that provide students with a genuine research experience. Students enrolled in CUREs develop or are given a research question with an unknown outcome, apply the scientific method to address the question, collect and analyze data, and communicate their results(3–5). Students that participate in a CURE learn the necessary skills and techniques they need to carry out the tasks required(6–8), and at the same time gain confidence in their ability to engage in the scientific process(9–11). Assessment of student learning gains reveal that CUREs improve students’ abilities to think critically, interpret data, communicate results, and collaborate as a team, when compared to traditional lab courses(12–17). A critical aspect of CUREs, as well as independent research, is obtaining the foundational skills and introductory training needed for understanding a specific system and/or research topic of interest. Several inquiry-based learning models have been developed to provide students with these foundational skills prior to their independent research projects(18–20).

Here we describe a level 1 guided-inquiry laboratory module(21) that prepares students for higher levels of guided inquiry and CUREs. This module is employed in the first half of our upper division undergraduate CURE on developmental genetics, which is used to prepare students for independent inquiry-based group research projects that occur in the second half of the course. In this module students are introduced to the research organism, *Caenorhabditis elegans* (*C. elegans*), to explore the concepts of RNA interference (RNAi) and codon optimization. *C. elegans* offers many advantages that make it an ideal research organism, such as a fast life cycle, large brood sizes, and easy access to genetic manipulation by forward and/or reverse genetic approaches(22–24). Additionally, they are transparent, which allows for visualization of all tissue types, and the real-time visualization of fluorescently-tagged reporter proteins expressed in various tissues of interest(25, 26). Using the protocols outlined in this paper, students will conduct an RNAi experiment using *C. elegans* where they will visualize first-hand how RNAi depletes a GFP-tagged transgene and how codon optimization significantly impacts gene expression.

Similar to other laboratory modules involving *C. elegans* and RNAi (27, 28), this module allows students to make connections between the concepts they learn about in molecular and developmental genetics with the observations they make while conducting the RNAi experiment in the laboratory. At the same time, students gain experience working with an organism commonly used in the research setting. Our goal is that the experiences gained from this module will prepare students for higher levels of guided inquiry by enhancing their scientific and communication skills. This module can also be used as a “stepping-stone” or “bootcamp” exercise to provide students with a set of skills and tools for the inquiry-based module of a CURE using *C. elegans* as a model organism. Finally, modules like the one presented here have a positive impact on student development and at the same time provide the prerequisites needed for success in CUREs.

### Intended Audience

This laboratory module was employed in the first half of upper-level undergraduate developmental genetics laboratory course (BIO327) at Stony Brook University. Most students enrolled in the course were Juniors or Seniors; however, the module can be implemented as a “bootcamp” exercise for first-year graduate students to gain hands-on bench experience working with *C. elegans*.

### Required Learning Time

The module requires a minimum of four lab sessions of approximately 3 hours each. We found this was ample time for students to become accustomed to working with *C. elegans* and proficient in the necessary skills needed to complete the module. Instructors can adjust the timing of the module to any desired length of time they feel is appropriate.

### Prerequisite Student Knowledge

To complete this module, students should have taken introductory biology and introductory biology laboratory that exposes students to core biological principles, such as gene expression, and basic organismal biology. It is highly encouraged that students have familiarity with basic laboratory procedures, such as micropipetting and sterile techniques. Prior to the module, all necessary materials and information needed to complete the assignments are provided, and students receive an introduction to RNAi, codon optimization, and basic microscopy. We highly recommend this module be implemented after students have gained a basic understanding of how to work with and manipulate *C. elegans* (29).

### Prerequisite Teacher Knowledge

Instructors implementing this course should have experience working with *C. elegans*. Ideally, there should be access to temperature controlled incubators and other equipment needed for *C. elegans* maintenance(29). Importantly, a good understanding of concepts involving RNA interference(30) and codon optimization(31, 32) is essential for this module. We have provided a PowerPoint presentation with an accompanying script for instructors to use when teaching students about RNAi and codon optimization (Supporting file S1. A Laboratory Module-GFP RNAi *C. elegans* Lecture). In addition, we have provided instructors with a list of common misconceptions and questions from students when conducting the module (Supporting file S14. A Laboratory Module-Common Student Misconceptions and Questions). Lastly, instructors should know how to operate stereomicroscopes, compound light microscopes, epifluorescence light microscopes, and image processing software, such as Fiji/ImageJ(33).

## SCIENTIFIC TEACHING THEMES

### Active Learning

Several active learning strategies that are implemented throughout this module include a modified think-pair-share exercise, clicker polling questions, and a peer review activity. Students are asked a series of clicker polling questions during the RNAi lecture that focus on students’ conceptual understanding of RNAi(Supporting file S1: A Laboratory Module-GFP RNAi *C. elegans* Lecture). For the modified think-pair-share exercise, prior to the GFP RNAi experiment, students are assigned a GFP RNAi worksheet to work on independently at home (Think component) (Supporting file S5. A Laboratory Module-Student GFP RNAi Worksheet). In brief, the worksheet contains a series of questions, which promotes independent thinking about the RNAi experiment, and guides the students in formulating their hypothesis (see below). After completing the worksheet at home, students form into groups during their next lab session (Pair component), and while preparing for the GFP RNAi experiment, they are encouraged to discuss amongst themselves their findings and share their hypotheses. When conducting the experiment in class, instructors and teaching assistants approach each group and ask them to share their findings from the worksheet (Modified share component). This is followed by a series of additional questions asked by the instructor(s) to further test their understanding of RNAi(Supporting file S3-A Laboratory Module-GFP RNAi Module Worksheet Discussion Questions & Answers). This modified share component of the think-pair-share activity provides an equitable opportunity for all groups to validate their understanding rather than a select few groups sharing in front of the entire class(34).

For the peer review activity, after completing their lab report assignment (Supporting file S2. A Laboratory Module-Grading Rubric and Example Lab Report), students are randomly assigned to review and constructively critique another fellow student’s laboratory report. Students are first instructed to upload their lab reports into their designated Google Drive folder as a Google document (.docx file), which allows their peer reviewer to easily comment on the reports in real-time and create editable suggestions. Each peer reviewer is instructed to provide feedback and suggestions on the required components of their lab report (i.e. Nucleotide alignment figure, data table of quantification, etc.; See Supporting file S2. A Laboratory Module-Grading Rubric and Example Lab Report). Specifically, each student must review each other’s work with specific criteria in mind, such as the clarity of writing (Is a hypothesis clearly stated and is there enough detail to understand the results?), statistical tests performed (Are appropriate statistical tests performed on the data?), and organization of data (Is the data organized in such a way that results can be clearly interpreted?). We emphasized to the students that all critiques should be professional and constructive and should avoid any condescending language. The purpose of this assignment is to get students to become familiar with the scientific process of peer review, appreciate the importance of quality work in delivering a clear message, and encourage the exchange of ideas. Most importantly, peer review as an active learning strategy stimulates students to reflect on their own written work, and results in improvements on their own writing(35, 36).

### Assessment

Student assessments are conducted at multiple levels throughout the module. During the short introductory lectures given, students are asked a series of clicker polling questions incorporated into the lecture (Supporting file S1: A Laboratory Module-GFP RNAi *C. elegans* Lecture) and are informally assessed based on whether their answers are correct or incorrect. We also informally assessed students on their ability to provide constructive feedback during the peer-mediated review activity (see above), which counted as part of their participation grade, as well as their ability to answer questions asked by instructors during the modified think-pair-share activity (Supporting file S3-A Laboratory Module-GFP RNAi Module Worksheet Discussion Questions & Answers). Although we did not require students to submit a lab notebook for the course, we did create a Google Drive folder organized by class section, where students were encouraged to upload their quantified data and any observations made into their individualized sub-folders. They were also asked to submit their completed lab report as a Google doc for grading by instructors and teaching assistants into their individualized sub-folder. Along these lines, students are formally graded based on the quality of their lab report assignment, which includes a graph and table of their results, a “publication quality” figure using acquired fluorescence micrographs along with an accompanying figure legend, and a results text write-up (Supporting file S2. A Laboratory Module-Grading Rubric and Example Lab Report).

### Inclusive Teaching

We have designed this module to be all-inclusive by differentiating content and lesson material to reach all types of learners. The hands-on activities of this module capture the attention and engagement of kinesthetic and tactile learners. Our short lectures that contain images, provide written instruction, and facilitate discussion amongst the class are accommodating to both visual and auditory learners. Given that Stony Brook University consists of a highly diverse population of students, during group activities, we can easily divide our class into diversified groups at random using a freely available random name picking software called wheeldecide.com. We highly recommend that instructors utilize this tool given that it avoids any self-selection or instructor selection biases.

To ensure that students feel welcomed, we establish classroom “etiquette”, similar to that suggested by Tanner 2013, where we emphasize that all students are expected to support one another and share their ideas in a judgement free manner (37). On the very first day of class, we implemented an ice-breaker activity, called “catch the ball”, where all students and faculty “threw” around an imaginary ball to one another, and those who “caught” the ball on a turn introduced themselves, shared their interests, hobbies, and goals. We suggest a similar activity be implemented during the start of the course so that instructors can familiarize themselves with their students. To further create an inclusive learning environment, we ensure that all students have the means to be successful in the module. We ensure class material for the lesson is posted on Blackboard and/or in Google Drive in a timely fashion so that students can access it prior to the start of class and after. For students that may not have equal access to technology, hard copies, as well as digital copies, of assignments and lab protocols were provided to students. We also hold office hours on request and have discussion boards available so everybody can benefit from each other’s questions and/or discussions. Moreover, based on information obtained from class assessments (see above) and observations, students who have difficulties with any of the class content receive extra support and guidance as needed. Thus, the module ensures equity and inclusivity by reaching all types of learners and ensuring students receive the support they need to succeed in the module.

## LESSON PLAN

### Overview of the module

In this module, students will use *C. elegans* as a model organism to understand how codon optimization significantly impacts gene expression and how RNAi interference can precisely downregulate gene activity.

Specifically, students will work with two GFP-expressing *C. elegans* strains, where one strain expresses a non-codon optimized (NCO) GFP fusion protein (GFP_NCO_), while the other strain expresses a codon optimized (CO) GFP fusion protein (GFP_CO_). The GFP_NCO_ and GFP_CO_ tags are each fused to the histone protein, *his-58* (H2B), and are each expressed under the control of a ubiquitous promoter, *eft-3,* which promotes expression in all cells. Students will treat each strain with an empty vector (control) RNAi bacterial clone or an RNAi bacterial clone that produces double stranded RNA (dsRNA) specific to only the non-codon optimized GFP variant (GFP_NCO_) (Review Timmons and Fire, 1998 for a detailed description on how RNAi works in *C. elegans*). Through fluorescence microscopy, students will observe differences in GFP expression in each strain due to codon optimization, and they will observe that significant depletion occurs only in the strain expressing *eft-3>H2B::GFP_NCO_*. From their understanding of RNAi and codon optimization, we anticipate that students will be able to accurately predict these results and explain why depletion occurs only in the strain expressing *eft-3>H2B::GFP_NCO_*.

Prior to the module, we present students with a lecture on gene regulation (Supporting file S1: A Laboratory Module-GFP RNAi *C. elegans* Lecture). We have provided instructors with a script that accompanies the lecture (Supporting file S1: A Laboratory Module-GFP RNAi *C. elegans* Lecture). We recommend that instructors review Corsi et al., 2015 for a comprehensive overview of *C. elegans* as a research organism. The lecture discusses the topic of RNAi, which is a biological process where in the presence of exogenous dsRNA results in post-transcriptional gene silencing(23, 38–41). One method used to administer *C. elegans* with dsRNA is to feed them with *E. coli* expressing a vector capable of producing dsRNA, that is complementary to a target gene of interest(42–44). *C. elegans* are unique in that they have a systemic RNAi response, meaning that dsRNA spreads throughout all tissues, with the exception of most neurons(45, 46). Thus, loss-of-function phenotypes for genes of interest can be assessed in almost any tissue of interest using RNAi.

We also provide our students with a brief overview of codon optimization when discussing the GFP RNAi worksheet (Supporting file S5. A Laboratory Module-Student GFP RNAi Worksheet). For a detailed overview of codon optimization, we highly recommend instructors review Hanson and Coller, 2018. Codon optimization is the modification of a DNA sequence such that the frequency of codons used by a particular organism, for a specific amino acid, is taken into consideration when designing gene fusions or introducing exogenous DNA(47–49). Codon optimization significantly enhances the expression level of a particular protein due to the correlation between codon usage and tRNA abundance, and mRNA stability(50–53). Thus, the expression levels of codon optimized genes will be more robust than those of non-codon optimized genes.

Overall, we anticipate this module will fulfill several goals, which include increasing student proficiency in using the scientific method and development of critical thinking skills. After completing this module, students will be able to conduct controlled experiments using a model organism. In addition, they will be able to explain what RNAi is and how it can be used to assess loss-of-function phenotypes for any gene of interest. Lastly, students will be able to state the importance of codon optimization as it pertains to gene expression.

### GFP RNAi Module

#### Student and instructor preparation

To carry out the GFP RNAi module, both instructors and students should have a general understanding of *C. elegans* development(26). To prepare the students for the experiment, we presented a short lecture on RNAi and codon bias (Supporting file S1: A Laboratory Module-GFP RNAi *C. elegans* Lecture) and devised a “GFP RNAi worksheet” (Supporting file S5. A Laboratory Module-Student GFP RNAi Worksheet). The goal of this worksheet is to drive students to formulate hypotheses as to whether the GFP_NCO_ RNAi clone will efficiently knock down GFP intensity levels in the strain expressing H2B::GFP_CO_ or H2B::GFP_NCO_. In this worksheet, the students are provided with the nucleotide and amino acid sequences for the codon and non-codon optimized H2B::GFP fusion proteins, as well as the dsRNA nucleotide targeting sequence (in DNA form) for the GFP_NCO_ RNAi clone (Supporting file S5. A Laboratory Module-Student GFP RNAi Worksheet). Using the sequences provided, students will make a pairwise sequence alignment using EMBOSS Needle (https://www.ebi.ac.uk/Tools/psa/emboss_needle/). They will then compare the percent similarities between the different sequences and determine whether the dsRNA targeting sequence for GFP_NCO_ RNAi is most similar to H2B::GFP_CO_ or H2B::GFP_NCO_. Through this process, students will see that the dsRNA targeting sequence encoded by the GFP_NCO_ RNAi clone is 100% identical to the GFP_NCO_ sequence and not the GFP_CO_ sequence, and therefore should hypothesize that the GFP_NCO_ RNAi clone will significantly deplete the H2B::GFP_NCO_ strain. Students will also appreciate that the control RNAi clone is called “empty vector” because it does not produce a dsRNA product.

To conduct the RNAi experiment, the students should grow up both the *eft-3>H2B::GFP_CO_* and *eft-3>H2B::GFP_NCO_* strains (DQM583 and DQM594, respectively) initially on NGM plates containing an *E. coli* diet (*E. coli* variant OP50)(29) (Supporting file S4. A Laboratory Module-Detailed Protocols, Section II). Please note that worms are initially grown on OP50-seeded NGM plates prior to treatment with a different variant of *E. coli* (variant HT115(DE3)) that expresses dsRNA-producing vectors. Along these lines, RNAi plates utilize the HT115 variants of *E. coli* that can produce dsRNA rather than OP50 (44). Prior to the experiment, instructors should have RNAi plates made that contain *E. coli* specific to empty vector control (*T444T*) and GFP_NCO_ (Supporting file S4. A Laboratory Module-Detailed Protocols, Section IV). Moreover, we recommend that instructors have additional RNAi plates as students do tend to make occasional errors, such as accidentally contaminating plates. To acquire a sufficient number of L1 larvae for the experiment, we recommend that instructors ensure that students have at least six NGM plates containing ~250 gravid adults for bleach synchronization(29, 54).

When NGM plates are full of gravid adults (~250 adults on each plate), students should treat each strain with alkaline hypochlorite solution(55) (Figure 1, Step 1) (Supporting file S4. A Laboratory Module-Detailed Protocols, Section V) to create synchronized L1s. Approximately 50-100 L1 animals should be pipetted onto control and GFP_NCO_-specific RNAi plates (Figure 1, Step 2). Individual RNAi plates should have no more than ~50-100 worms to prevent overcrowding and depletion of the *E. coli* food source (Figure 1, Step 2) (Please note that instructors may need to do the bleaching and plating steps for students to allow for efficient completion of the RNAi experiment). The L1s are then cultured on the RNAi plates at the desired temperature until the L3 or L4 stage is reached (Figure 1, Step 3). Once the desired stage is reached, students can mount the animals on microscope slides for imaging. To immobilize the worms for image analysis, worms can be added to a droplet of M9 buffer (5μL) in the center of the slide, surrounded by Nemagel solution (InVivo Biosystems) or ~1 μl of M9 containing 5mM levamisole (Figure 1, Step 3). We recommend that students pick ~10 animals for imaging at a time. (Supporting file S4. A Laboratory Module-Detailed Protocols, Section VI).

**Figure 1:**
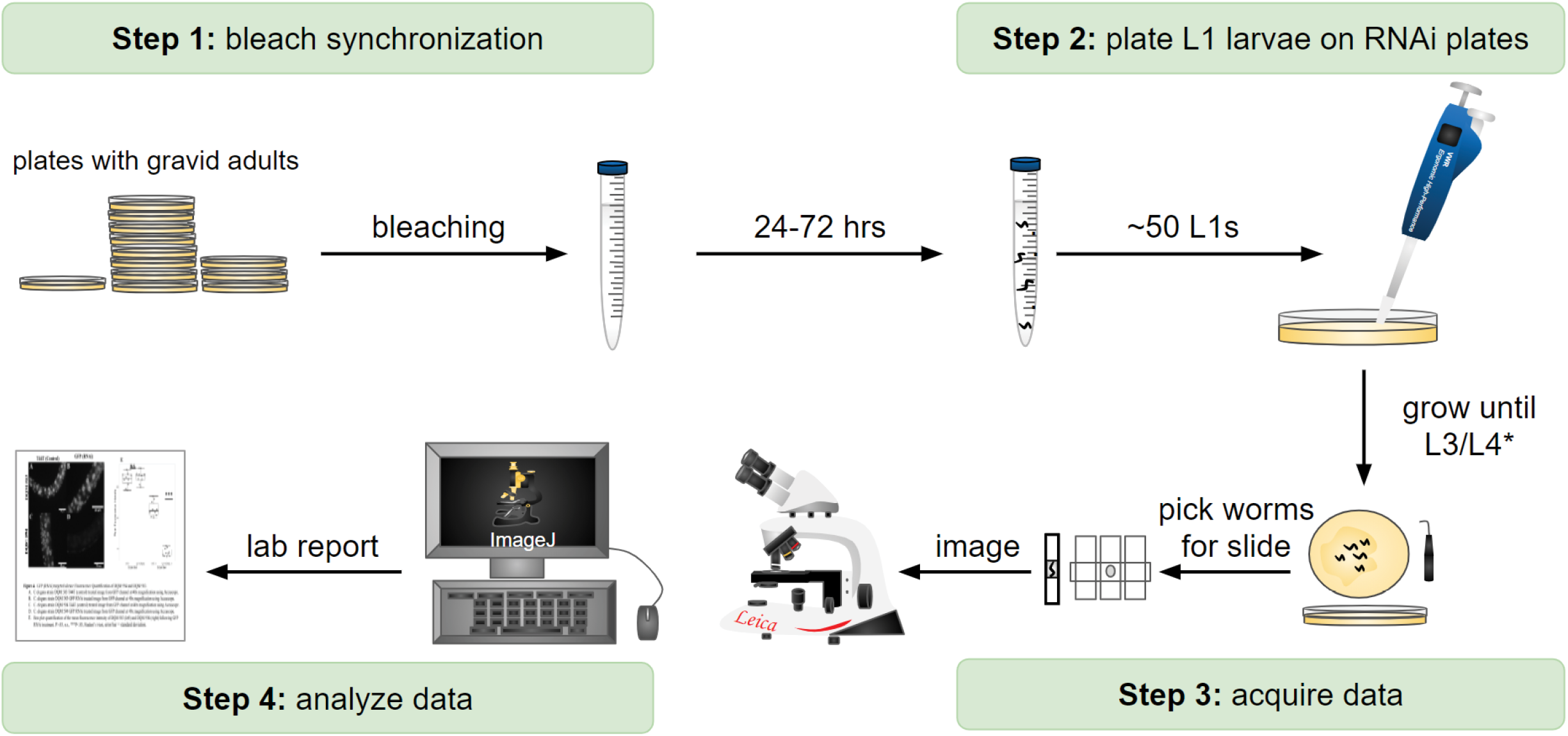
Workflow diagram of the GFP RNAi module. **(Step 1)** For both codon optimized (*eft-3>H2B::GFP*_*CO*_) and non-codon optimized (*eft-3>H2B::GFP*_*NCO*_) strains, 10 OP50-seeded NGM plates each containing ~50-100 *C. elegans* gravid adults were treated with alkaline hypochlorite solution to obtain synchronized larvae. **(Step 2)** After 24 hours in M9 solution (and up to 72 hours), hatched L1 larvae are ready for plating onto RNAi plates (control or *T444T* RNAi and GFP_NCO_ RNAi). For optimal RNAi efficiency and to avoid overcrowding/starvation, ~50 worms per plate will suffice. **(Step 3)** L1 larvae are grown until the L3/L4 larval stage and then mounted on 5% agarose pad slides (containing levamisole (anesthetic) and a drop of M9 buffer) for image acquisition. *Growth times will vary based on temperature (see text for more details). **(Step 4)** Images are acquired and then analyzed using Fiji/ImageJ to determine the mean fluorescence intensity. Results are briefly explained in the lab report and submitted along with a publication quality figure with figure legend.

#### Student Experimental Results

Students quantified H2B::GFP fluorescence depletion using two wide-field epifluorescence microscopes, the Accu-Scope or Leica DMLB (Figure 1, Step 4, Figure 2 A and B). For imaging consistency, instructors should predetermine the imaging settings (exposure time, magnification, camera gain and binning) using the *eft-3>H2B::GFP*_*CO*_ strain (DQM583) as a baseline due to it having the highest expression level. Both *eft-3>H2B::GFP*_*CO*_ and *eft-3>H2B::GFP*_*NCO*_ strains were imaged for each RNAi treatment (control and GFP_NCO_). From the data acquired by the students, several qualitative observations were made (Figure 2 A and B). First, the overall fluorescence intensity of the GFP_CO_ strain was visually much brighter than the GFP_NCO_ strain. Second, treating the GFP_NCO_ strain with GFP_NCO_ RNAi strongly reduced the fluorescence intensity of GFP, whereas treating the GFP_CO_ strain with GFP_NCO_ RNAi did not (Figure 2 A and B, *eft-3>H2B::GFP* column). Third, in the GFP_NCO_ strain treated with GFP_NCO_ RNAi, although the fluorescence intensity of GFP was strongly reduced, some nuclei still showed high levels of GFP, which correspond to the cells that are insensitive to RNAi, most notably neurons (Figure 2A, *eft-3>H2B::GFP*_*NCO*_; GFP_NCO_ RNAi).

**Figure 2.**
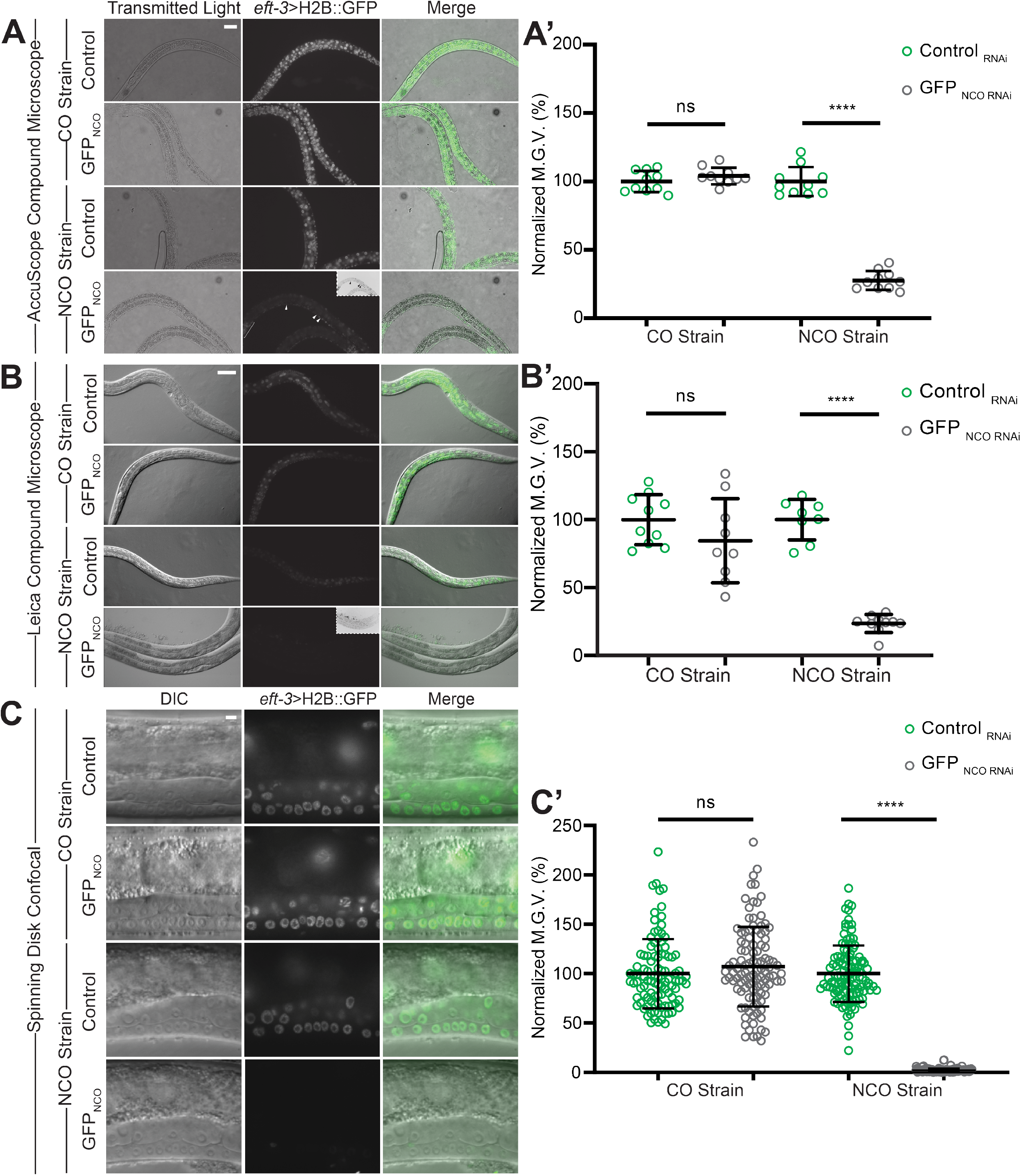
RNAi specificity between codon optimized genes and non-codon optimized H2B::GFP strains. **(A-C)** Representative DIC and fluorescence micrographs of *eft-3*>H2B::GFP_CO_ (CO strain) and *eft-3*>H2B::GFP_NCO_ (NCO strain) strains treated with RNAi against empty vector (control) or GFP_NCO_. Micrographs were collected by students using AccuScope (A) or Leica (B) epifluorescence compound microscopes or were collected by instructors using a custom modified upright spinning disk confocal microscope (C). Images represent either the whole-body (A and B) or the midsection (C) of the animal. **(A’-C’)** Quantification of the normalized mean gray value (Normalized M.G.V) of H2B::GFP expression, shown as a percentage, in CO and NCO strains. Mean fluorescence intensity for either whole-body (A and B) or midsection nuclei (C) are shown. N≥8 animals per treatment (A’ and B’) or N≥6 animals per treatment and n>100 midsection nuclei (C’). Error bars denote mean with SD. Scale bars: 50 μm (A), 25 μm (A), 5 μm (C). Arrow heads (A) denote neurons in the *eft-3*>H2B::GFP_NCO_ not affected by GFP_NCO_ RNAi; inset shows the same image inverted. Statistical analysis was performed using an unpaired, two-tailed, Student’s *t-test* with Welch’s correction or Mann-Whitney U test. n.s.: not significant. p-value ****≤0.0001.

To analyze the data quantitatively, we instructed students to quantify whole-body GFP fluorescence intensity for 10 animals from each strain grown on control and GFP_NCO_ RNAi, using Fiji/ImageJ2(33). Briefly, the entire body of each worm was outlined and the mean fluorescence intensity (MFI) was then measured for both GFP and an area of background. The background MFI measurement was then subtracted from the GFP MFI measurement to reduce background noise and obtain a mean gray value (MGV). Mean gray values were normalized by dividing the MFI in RNAi-treated animals by the average MFI in control-treated animals (Supporting file S4. A Laboratory Module-Detailed Protocols, Section VII; Supporting file S6. A Laboratory Module-Student Instructions for GFP RNAi Module; Supporting file S7. A Laboratory Module-Student Transcripts for Tutorial Videos 1-5; Supporting file S8. A Laboratory Module-Opening Images in Fiji/Image J Tutorial Video; Supporting file S9. A Laboratory Module-Measuring Mean Fluorescence Intensity for Single Z data Tutorial Video; Supporting file S10. A Laboratory Module-Measuring Mean Fluorescence Intensity for Confocal Z-stack data Tutorial Video; and Supporting file S11. A Laboratory Module-Compiling Data Tutorial Video). The mean gray values obtained from each imaging system (microscope) are plotted next to their respective micrographs (Figure 2 A’ and B’).

By plotting the normalized MGV, students were able to clearly see that treating the GFP_NCO_ strain with GFP_NCO_ RNAi significantly reduced the expression of GFP compared to control-treated animals (Figure 2A, 2A’, and 2B, 2B’, NCO strain; control RNAi vs. GFP_NCO_ RNAi). Moreover, the students noted that treatment with GFP_NCO_ RNAi had no effect on GFP expression levels in the codon-optimized strain (Figure 2A, 2A’, and 2B, 2B’, CO strain; control RNAi vs. GFP_NCO_ RNAi). To determine the statistical significance of their results, students performed a Student’s t-test comparing control MGV’s to the MGV’s for the GFP_NCO_ and GFP_CO_ strains. To assess whether the students successfully carried out the experiment, we instructed them to document their results as part of their lab report assignment by creating a publication quality figure. Their figures included representative images of their fluorescent micrographs, along with a dot plot of their quantified data, table of their raw data values, and written description of their results (Supporting file S2. A Laboratory Module-Grading Rubric and Example Lab Report and Supporting file S6. A Laboratory Module-Student Instructions for GFP RNAi Module). From these results, and the results obtained from the GFP RNAi worksheet, it should become evident to the students that RNAi specificity is largely dependent on the sequence homology/similarity between the target gene sequence and the sequence of the dsRNA produced by the RNAi clone itself.

#### Extended Results (Optional)

Compared to wide-field epifluorescence microscopy, confocal microscopy improves resolution such that unwanted out-of-focus light is significantly reduced, and the detail of cellular objects is greatly enhanced(56). Thus, to show students high quality images of nuclear DNA labeled with H2B::GFP, we acquired spinning-disk confocal images for both the *eft-3>H2B::GFP*_*CO*_ and *eft-3>H2B::GFP*_*NCO*_ strains (Figure 2C, 2C’, 3, and 4). Importantly, these spinning-disk confocal images served to better illustrate some of the key concepts discussed in the lab module, such as codon optimization and lineage specific differences in RNAi susceptibility.

**Figure 3.**
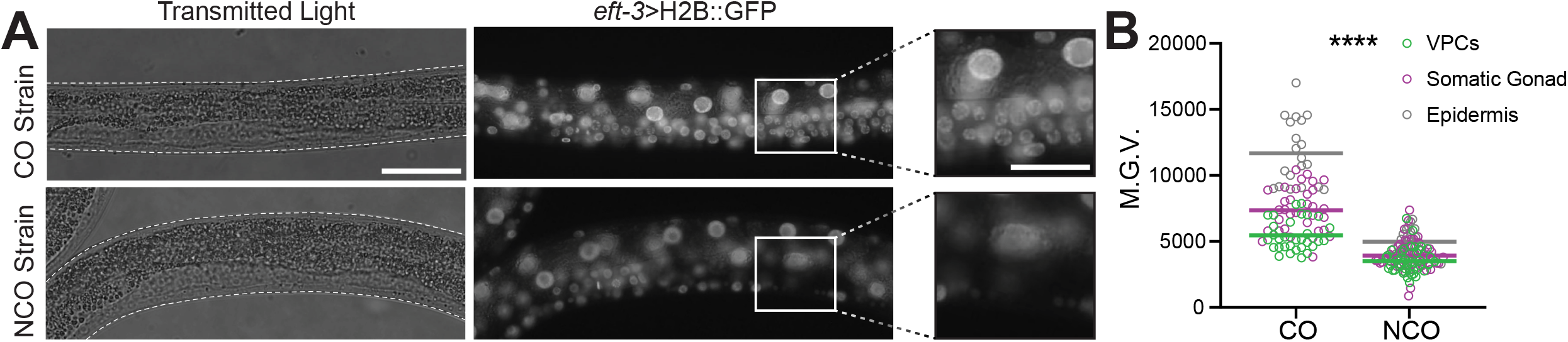
Codon optimization results in improved gene expression in the germline. **(A)** Representative DIC and fluorescence micrographs of the *C. elegans* germline for *eft-3*>H2B::GFP_CO_ (CO strain) and *eft-3*>H2B::GFP_NCO_ (NCO strain) strains. Insets represent increased magnification of the germline to emphasize expression differences between the CO and NCO strains. **(B)** Quantification of the Normalized M.G.V of H2B::GFP expression in individual nuclei of the midsection, shown as a percentage, in CO and NCO strains. Colored lines represent the mean M.G.V for individual lineages. N≥6 animals per strain and n>100 midsection nuclei. Scale bars: 50 μm (insets: 25 μm). Error bars denote mean with SD. Statistical analysis was performed using an unpaired, two-tailed, Mann-Whitney U test. p-value ****≤0.0001

**Figure 4.**
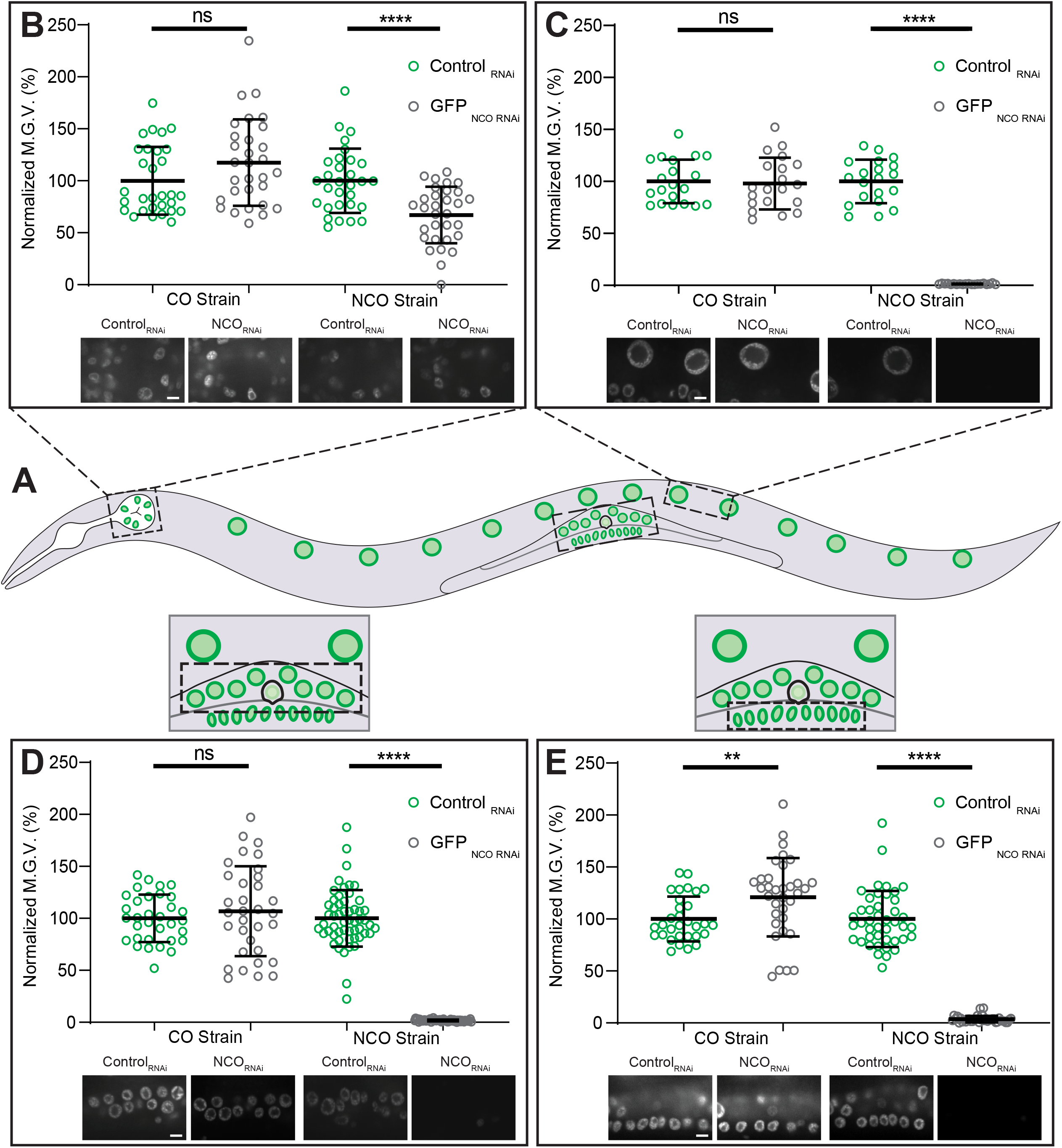
Lineage specific differences in RNAi susceptibility. **(A)** Cartoon schematic of a single *C. elegans* nematode with different cell lineages outlined. The cell lineages shown and quantified include pharyngeal cells (B), intestinal cells (C), somatic gonadal cells (D), and vulval precursor cells (VPCs) (E). **(B-E)** For each lineage, the Normalized M.G.V for individual nuclei was quantified in CO and NCO strains treated with control or GFP_NCO_ dsRNA. For each lineage, N≥6 animals/RNAi clone and n≥30 nuclei. Scale bars: 5 μm. Statistical analysis was performed using an unpaired, two-tailed, Student’s *t-test* with Welch’s correction or Mann-Whitney U test. n.s.: not significant. p-value **≤0.01, ****≤0.0001.

From the confocal fluorescence micrographs, it becomes more apparent that treatment with GFPNCO RNAi significantly reduces GFP fluorescence intensity in the GFP_NCO_ strain, but not in the GFP_CO_ strain (Figure 2C, 2C’; CO strain vs. NCO strain; GFP_NCO_ RNAi vs. control RNAi). To highlight the differences in expression levels between codon optimized and non-codon optimized H2B::GFP fusion proteins, we took spinning disk confocal images of the *C. elegans* germline. In general, codon optimized transgenes are more robustly expressed in the germline than non-codon optimized transgenes(57, 58). In line with this, H2B::GFP fluorescence expression was more robust in germ cells when GFP is codon-optimized as opposed to when it is non-codon optimized the transgene is silenced, likely due to piRNA sequences present in the GFP_NCO_ nucleotide sequence ((59)) (Figure 3; CO strain vs. NCO strain).

In *C. elegans*, certain cell lineages show different sensitivities to exogenous dsRNA. For example, neurons and pharyngeal cells are less sensitive to RNAi compared to other somatic tissues(45, 60–62). To emphasize to students that certain lineages are more resistant to RNAi, we acquired spinning-disk confocal images of nuclei from various cell lineages commonly studied in *C. elegans* (Figure 4A), such as pharyngeal cells (Figure 4B), intestinal cells (Figure 4C), somatic gonadal cells (Figure 4D), and vulval precursor cells (figure 4E). For each of the cell lineages examined, once again treatment with GFP_NCO_ RNAi significantly reduced GFP fluorescence intensity levels in the GFP_NCO_ strain, but not in the GFP_CO_ strain (Figure 4B-E). However, with respect to the GFP_NCO_ strain treated with GFP_NCO_ RNAi, the percent decrease in GFP intensity levels in the pharyngeal cells was much less than the decrease found in the other cell types examined (Figure 4B compared to Figures 4C-E). Thus, these observations can be used in the classroom to clearly illustrate to students that certain cell types show different sensitivities to exogenous dsRNA.

## TEACHING DISCUSSION

The laboratory module presented here teaches a variety of common techniques employed by *C. elegans* researchers and exposes students to various concepts in molecular genetics and microscopy. During this module, students will become proficient at working with a widely used research organism, be able to conduct controlled experiments, analyze data, produce publication quality images, and have a basic understanding of microscopy. In addition, students will have a solid foundation as to how RNAi works, how it can be used to study gene function, and the importance of codon optimization on proper gene expression

This module clearly illustrates that certain cell types are less or more prone to the effects of dsRNA treatment, and that codon optimization results in improved gene expression in tissues (i.e. the germline). The advantage of using a strain that drives ubiquitous expression of H2B::GFP is that it is extremely bright and nuclear localized, and therefore easily visible on widefield epifluorescence microscopes, which are commonly available in most laboratory classrooms. For classrooms that have access to high resolution microscopes, such as a spinning-disk confocal, this module can be easily adapted for use on those types of microscopes as shown in Figures 2C, 3, and 4. The additional benefit of the strains used in this module is that students can immediately see differences in depletion between H2B::GFP_CO_ and H2B::GFP_NCO_ upon GFP_NCO_ RNAi treatment.

Upon completing this module, students will acquire the basic foundational skills needed for independent inquiry-based research projects involving *C. elegans*. Some examples of inquiry-based research projects that can follow this module, as part of a laboratory course such as our developmental genetics course, include a reverse genetics screen to identify genes important for specific processes of interest, such as longevity. In this example, with the assistance of their instructor, students can design a simple research question, such as “Do fat metabolism genes play a role in regulating lifespan?”. Students can search the literature for fat metabolism genes of interest, use either the Ahringer or Vidal RNAi libraries (Source Bioscience) to isolate clones specific for those genes, and determine if their depletion reduces or enhances longevity. The search for genes can be conducted individually or as a group. If the instructor decides to have students work together in a group, each student can select a gene they are interested in and then collectively decide on one gene to limit their focus on. The instructor could then have groups present in front of the class, where each student in a group explains why they chose their gene of interest, and then further explain why as a group they decided to follow up on their agreed upon gene. Working in groups is highly encouraged given that it promotes inclusivity, encourages the sharing of opinions, gives each student a sense of responsibility, and enhances student learning as a whole(63, 64).

To experimentally determine if depletion of their gene of interest affects longevity, a lifespan analysis can be conducted(65). Here, students can take ~100 synchronized adult worms and feed them with an RNAi bacterial clone that produces dsRNA specific to their gene of interest or empty vector (as a control). The students can then monitor the worm’s survival over time until their death (defined as the inability to respond to prodding)(65). Students can plot their data in the form of a Kaplan-Meier survival curve, and their results can then be documented and written up in research paper format or as a lab report. Additionally, students can practice their communication and presentation skills by presenting their findings to the class. The independent inquiry-based research projects that follow this module are limitless and can focus on a wide range of cellular processes, such as cell cycle regulation, cellular invasion, stress-resistance pathways, vesicle trafficking, and much more.

Although most lecture and laboratory-based classrooms use expository styles of instruction, classrooms that utilize active learning styles of instruction significantly enhance student performance and learning outcomes(15, 66, 67). Examples of active learning strategies that have been implemented throughout this module include a variation of Think-Pair-Share(68) and a Peer Review activity (see section on “Active Learning”). Our modified Think-Pair-Share activity gives students an opportunity to independently test their understanding of a concept(s), facilitates dialogue and the exchange of ideas between individuals, and allows students to verify their understanding with an instructor by sharing their findings and results. In contrast to the traditional share component, discussing their findings privately with instructors is a modification of the think-pair-share activity that gives all groups an opportunity to share their understanding of class content, as opposed to only a few representative groups sharing their knowledge to the entire class(34). One large advantage of the peer review activity implemented in this module is that it allows students to become familiar with the scientific process of peer review. Additionally, it prepares students to become accustomed to giving and receiving feedback in the workforce(69), and stimulates students to reflect on their own written work, which results in improvements on their own writing(35, 36, 70).

One additional active learning strategy that can be utilized in this module is the Jigsaw method(71). The jigsaw method is a two-phase activity where students are responsible for learning course content and teaching it to their peers(72). Although this active learning strategy was not implemented in this specific module of our course, we have designed a jigsaw activity that can be administered while introducing students to codon bias and optimization (Supporting file S13. A Laboratory Module-Jigsaw Active Learning Activity & Post-Module Assessment (Optional)). In the first phase of the activity, students are divided into several groups or teams, where each team focuses on three activities: 1. Transcription and translation, 2. Codon bias, and 3. Sequence alignment. Although group sizes will vary depending on class size, we recommend that groups consist of three students. For each activity, a learning goal and learning objective is provided so that students have a broad understanding of the purpose of the activity and know what they should be able to complete at the end of the activity (Supporting file S13. A Laboratory Module-Jigsaw Active Learning Activity & Post-Module Assessment (Optional)). Additionally, instructions for each activity are provided for the students to follow to become “experts” in each activity (Supporting file S13. A Laboratory Module-Jigsaw Active Learning Activity & Post-Module Assessment (Optional)). To assess their understanding and mastery of the activities, we have developed a series of questions that are associated with the learning levels of Blooms Taxonomy (Supporting file S13. A Laboratory Module-Jigsaw Active Learning Activity & Post-Module Assessment (Optional)). We estimate that 30 minutes to 1 hour is sufficient to complete all three activities simultaneously; however, instructors may have to adjust their time needs accordingly. Once each group has “mastered” their activity, the second phase begins where new groups are created that consist of one student from each original group. In these new groups, each student or “expert” teaches the other about their expertise or the subject matter from the first phase of the activity. To determine if students are adequately taught the subject matter by their peers, we have developed a post-assessment activity that consists of 15 questions along with an answer key for instructors. The advantage of this activity is that it promotes cooperation between peers in a team-based setting and greatly improves student learning and retention(73). In all, there are various active learning strategies that can be implemented in this module, which foster peer-to-peer communication, promote student engagement, and stimulate higher-order thinking.

An additional advantage of this module is that it can be adapted for remote teaching and online learning. The RNAi lecture and imaging tutorials on various aspects of the module (i.e. measuring mean fluorescence intensity) can be held synchronously during the scheduled time of class by utilizing the share screen option in video conferencing apps, such as Zoom or Google Meet, or asynchronously by uploading the image analysis video tutorials supplied onto Blackboard, Google Drive (Supporting file S8. A Laboratory Module-Opening Images in Fiji/Image J Tutorial Video; Supporting file S9. A Laboratory Module-Measuring Mean Fluorescence Intensity for Single Z data Tutorial Video; Supporting file S10. A Laboratory Module-Measuring Mean Fluorescence Intensity for Confocal Z-stack Data Tutorial Video; and Supporting file S11. A Laboratory Module-Compiling Data Tutorial Video; and Supporting file S12. A Laboratory Module-Formatting Images for Figure Generation Tutorial Video). Depending on the instructor and/or institution, the module can be implemented in a fully remote, or in a hybrid fashion, with in-person and online components. If fully remote, instructors can teach image analysis alongside with their lectures over Zoom and provide students with access to previously acquired raw data sets from epifluorescence and/or confocal microscopes through Blackboard or Google Drive. The students can then take those images and quantify the data in front of their instructor over Zoom or some other platform (or at home if more time is needed). For the GFP RNAi worksheet, after working on it independently at home, the instructor could create groups using breakout rooms, allowing each group to discuss their findings in a team-based setting. After an allotted amount of time (i.e. 20-30 minutes), the instructor can then join each breakout room to hear their discussion. Alternatively, a hybrid setting approach could be implemented where students could come into class on specific days to acquire their data and then perform the quantifications and other components of the module (lab report generation, peer-review activities, etc.) online or at home on other days. We adapted this distance learning technique for the second half of our course during the SARS-CoV2 pandemic in the Spring of 2020 and 2021 and received positive feedback from our students about the adaptability of the course.

Whether fully in class, or online/hybrid, based on the knowledge gained from the tutorials, compiled raw data, and the GFP RNAi worksheet, students will be able to formulate their hypothesis, test it by analyzing the supplied data, and present their findings by generating a publication quality figure. One additional advantage of this module is that at the graduate level, it can be particularly useful for graduate student rotations and can serve as an introductory “bootcamp” or “stepping-stone” to introduce the experimental techniques used in *C. elegans* research. Here, entry-level graduate students who have not previously worked with *C. elegans* will have the opportunity to do so and can immediately start acquiring data by conducting a reverse genetics screen devised by the principal investigator and/or themselves. Over time, these students can become confident enough to develop and plan their own projects.

In summary, this module is an excellent resource for instructors interested in conveying a real-life science experience to their students and serves as an excellent opportunity for students to gain the hands-on experience they need in order to pursue a career in biology.

## Supporting information

Tutorial Video 1

Tutorial Video 2

Tutorial Video 3

Tutorial Video 4

Tutorial Video 5

## SUPPORTING MATERIALS

Supporting file S1: A Laboratory Module-GFP RNAi *C. elegans* Lecture

Supporting file S2. A Laboratory Module-Grading Rubric and Example Lab Report

Supporting file S3-A Laboratory Module-GFP RNAi Module Worksheet Discussion Questions & Answers

Supporting file S4. A Laboratory Module-Detailed Protocols

Supporting file S5. A Laboratory Module-Student GFP RNAi Worksheet

Supporting file S6. A Laboratory Module-Student Instructions for GFP RNAi Module

Supporting file S7. A Laboratory Module-Student Transcripts for Tutorial Videos 1-5

Supporting file S8. A Laboratory Module-Opening Images in Fiji/Image J Tutorial Video

Supporting file S9. A Laboratory Module-Measuring Mean Fluorescence Intensity for Single Z data Tutorial Video

Supporting file S10. A Laboratory Module-Measuring Mean Fluorescence Intensity for Confocal Z-stack Data Tutorial Video

Supporting file S11. A Laboratory Module-Compiling Data Tutorial Video

Supporting file S12. A Laboratory Module-Formatting Images for Figure Generation Tutorial Video

Supporting file S13. A Laboratory Module-Jigsaw Active Learning Activity & Post-Module Assessment (Optional)

Supporting file S14. A Laboratory Module-Common Student Misconceptions and Questions

## ACKNOWLEDGEMENTS

Funding for this work was provided by startup funds to D.Q.M from Stony Brook University. We are grateful to Michael A. Q. Martinez and Katie L. Palmisano for critical reading of the manuscript and helpful suggestions. We would like to thank all developmental genetics students who have taken this course for their enthusiasm and for making the course enjoyable. We give special thanks to the teaching assistants Scott Yang, Andrew Hillowe, Roy Raheb, Narges Zali, Sam Chiappone, Stephen Ruis, Nazia Jamil, Chris Zhao, Kateryna Davydovych, and Asim Nadeen for their hard work and dedication to teaching developmental genetics and course faculty, Gerald Thomsen and J. Peter Gergen. We would like to thank the lab coordinators Mary A. Bernero and Albert Wilkinson for their assistance with laboratory set up and preparation. We are grateful to Thomas Geer of Nobska Imaging, and thankful for MicroOptics, for their support in imaging and microscopy.

