## Supplemental File 1 for "A laboratory module that explores RNA interference and codon optimization through fluorescence microscopy using *Caenorhabditis elegans*"

### Slide 1
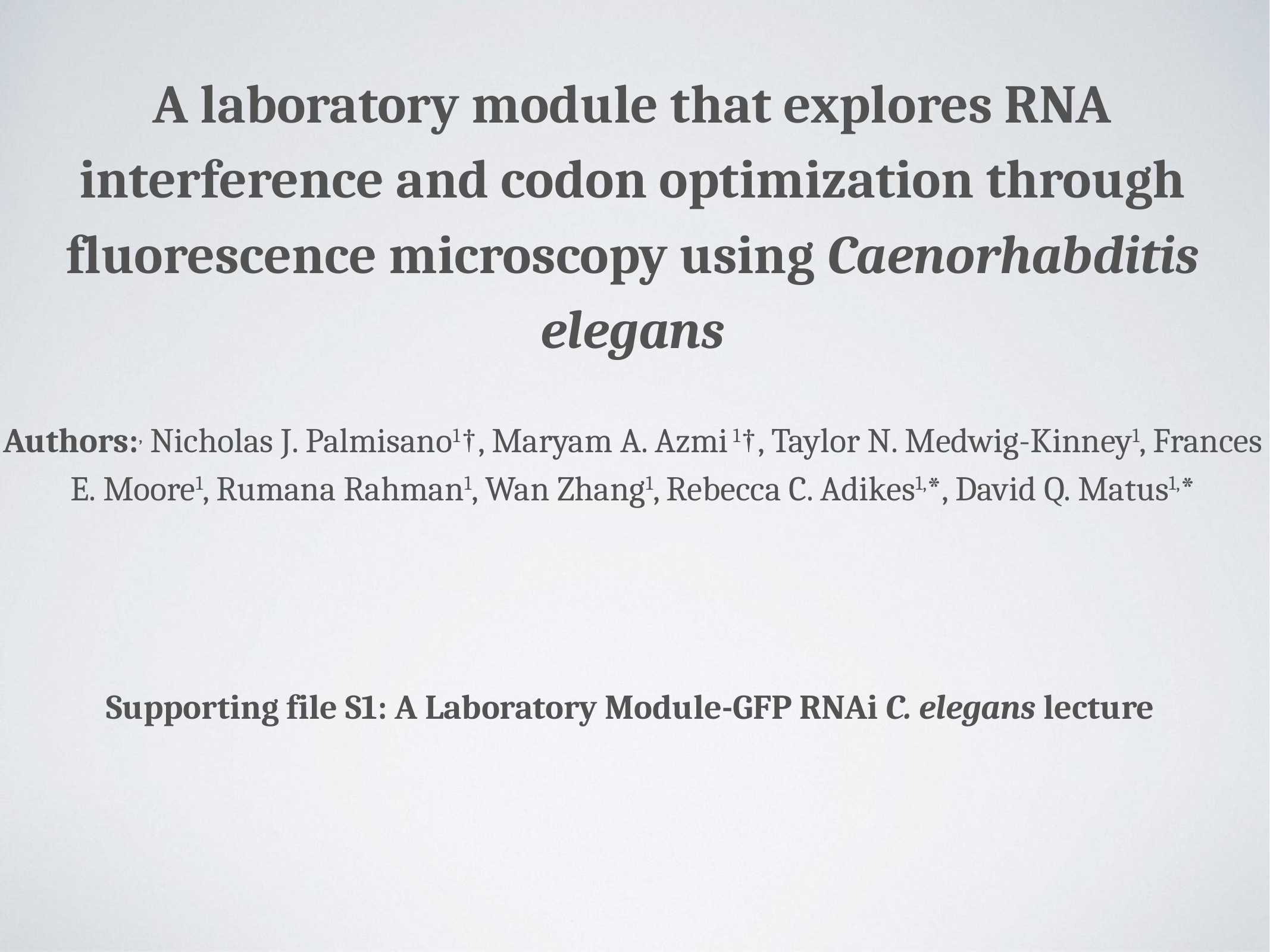

A laboratory module that explores RNA interference and codon optimization through fluorescence microscopy using Caenorhabditis elegans
Authors:^,^ Nicholas J. Palmisano^1^†, Maryam A. Azmi ^1^†, Taylor N. Medwig-Kinney^1^, Frances E. Moore^1^, Rumana Rahman^1^, Wan Zhang^1^, Rebecca C. Adikes^1,^*, David Q. Matus^1,^*
Supporting file S1: A Laboratory Module-GFP RNAi C. elegans lecture

### Slide 2
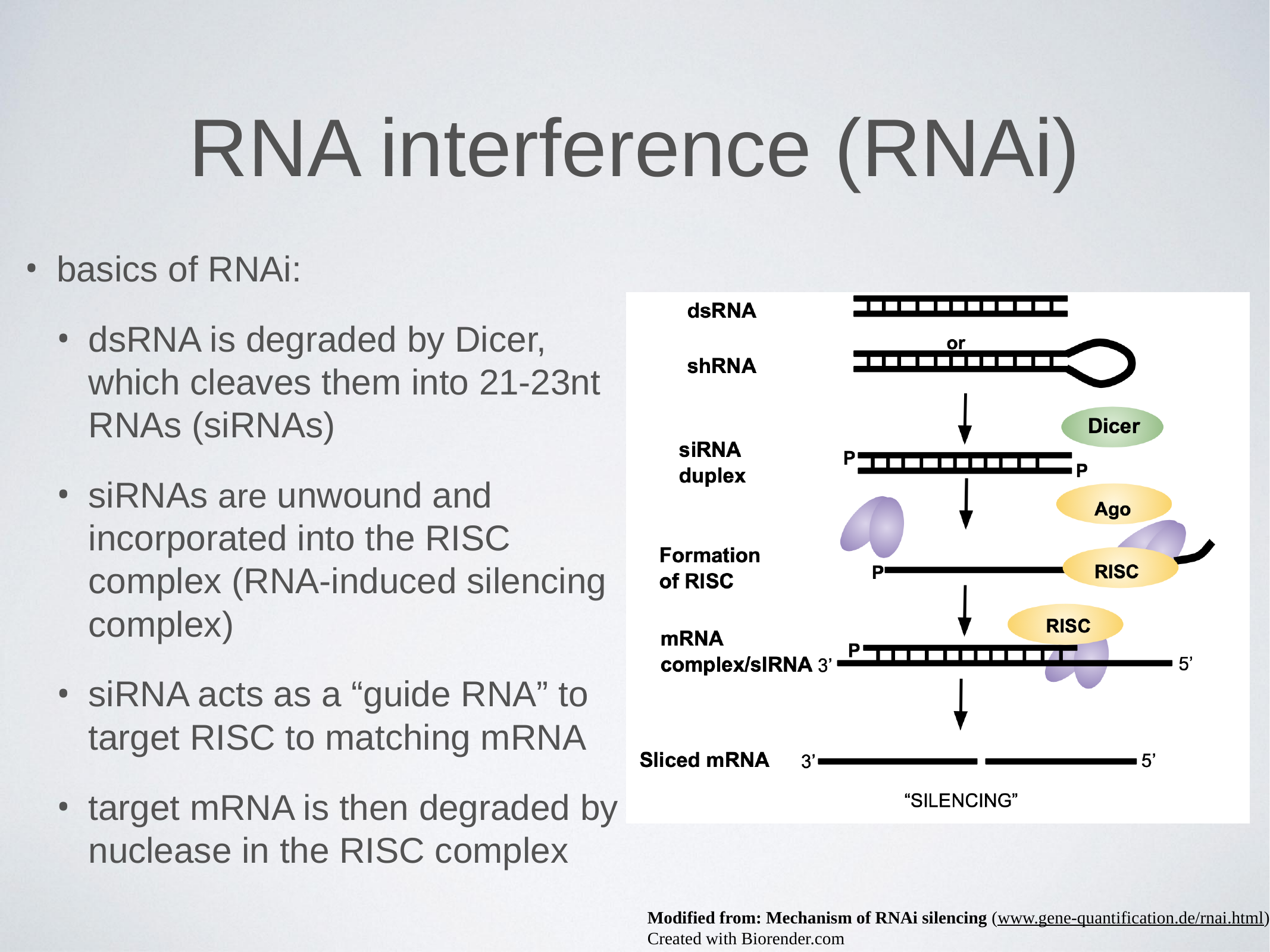

#### RNA interference (RNAi)
basics of RNAi:
dsRNA is degraded by Dicer, which cleaves them into 21-23nt RNAs (siRNAs)
siRNAs are unwound and incorporated into the RISC complex (RNA-induced silencing complex)
siRNA acts as a “guide RNA” to target RISC to matching mRNA
target mRNA is then degraded by nuclease in the RISC complex
Modified from: Mechanism of RNAi silencing (www.gene-quantification.de/rnai.html)
Created with Biorender.com

### Slide 3
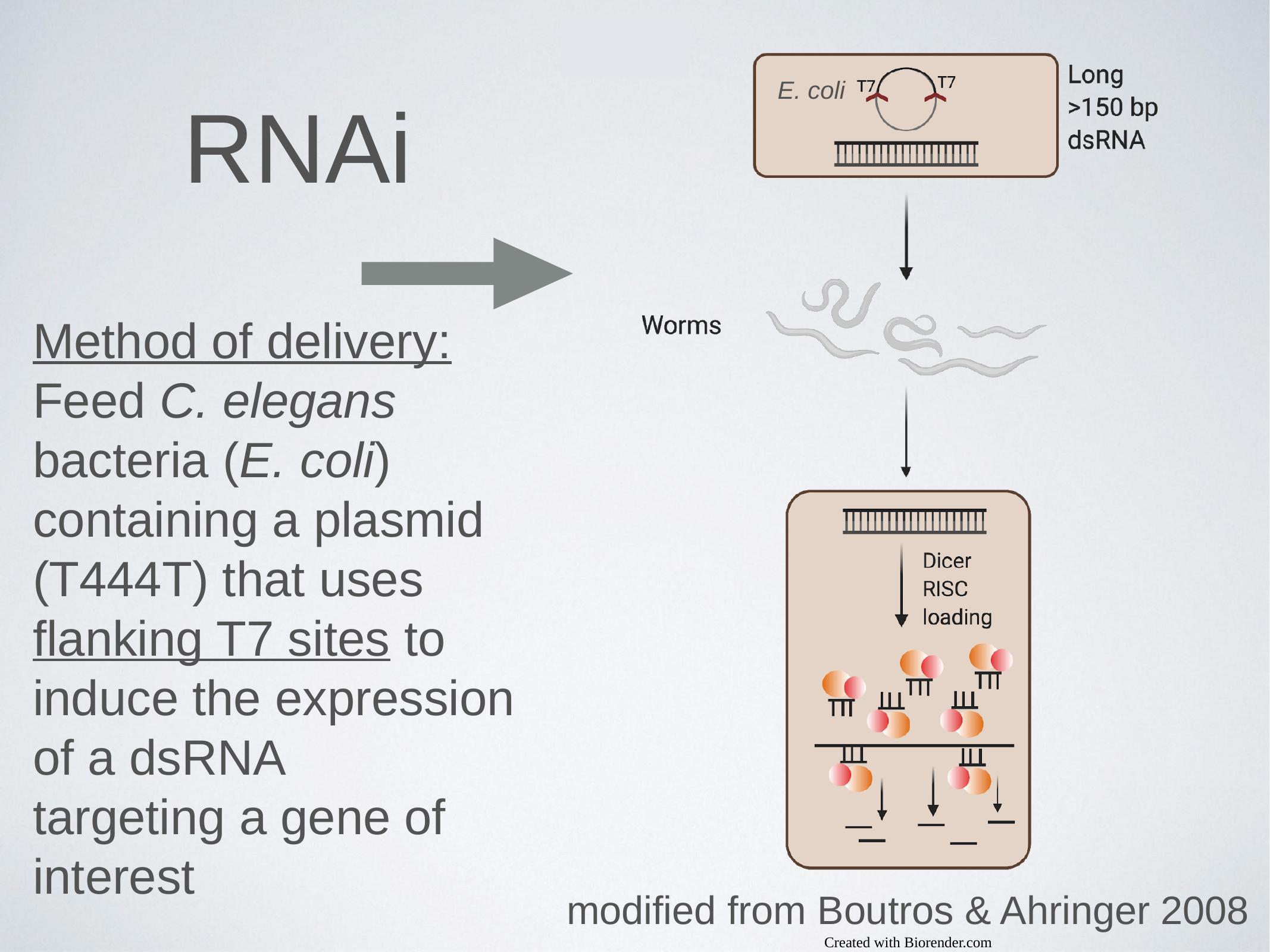

#### RNAi
E. coli
Method of delivery:
Feed C. elegans bacteria (E. coli) containing a plasmid (T444T) that uses flanking T7 sites to induce the expression of a dsRNA
targeting a gene of interest
modified from Boutros & Ahringer 2008
Created with Biorender.com

### Slide 4
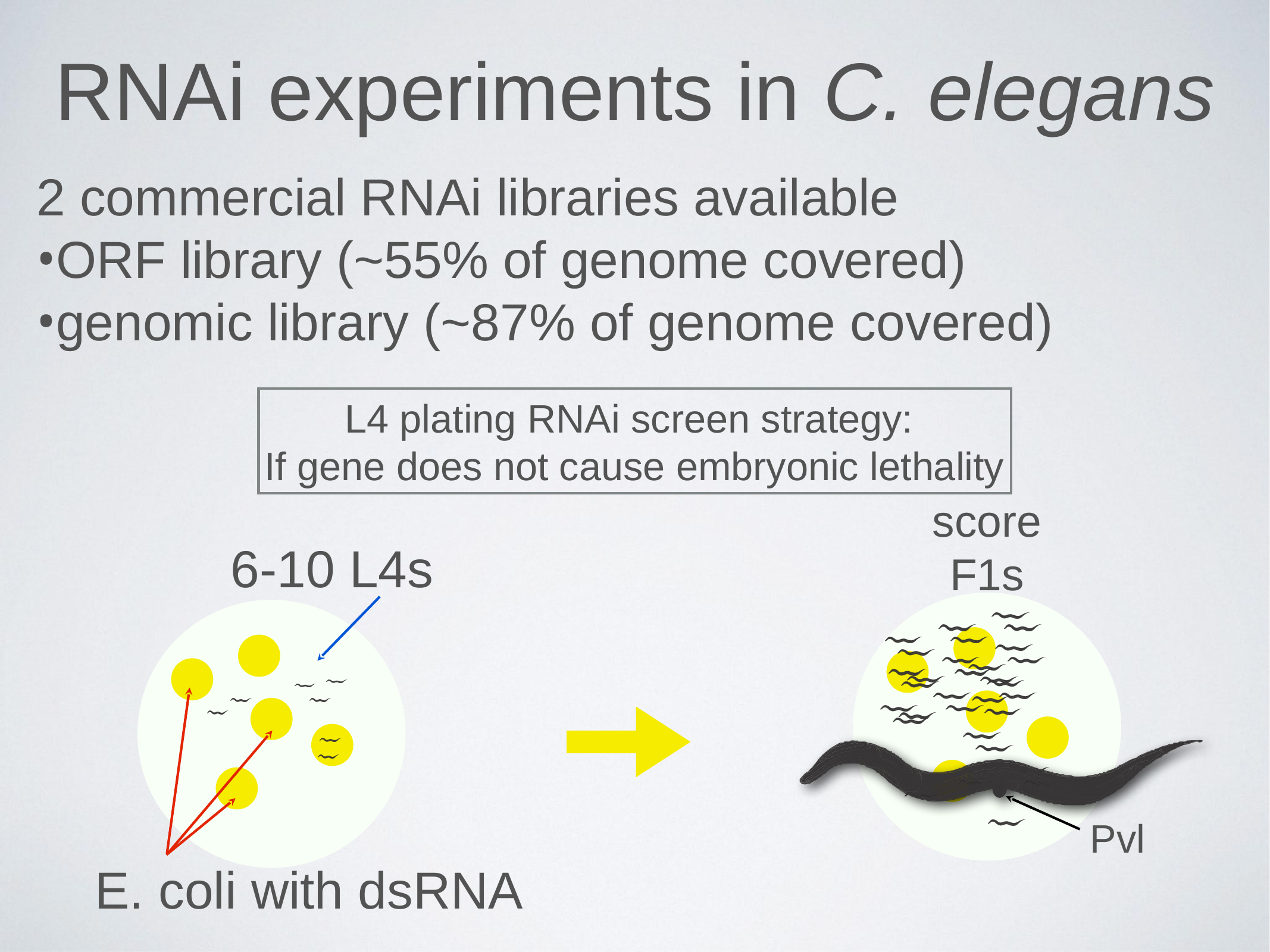

#### RNAi experiments in C. elegans
2 commercial RNAi libraries available
ORF library (~55% of genome covered)
genomic library (~87% of genome covered)
L4 plating RNAi screen strategy:
If gene does not cause embryonic lethality
score F1s
6-10 L4s
E. coli with dsRNA
Pvl

### Slide 5
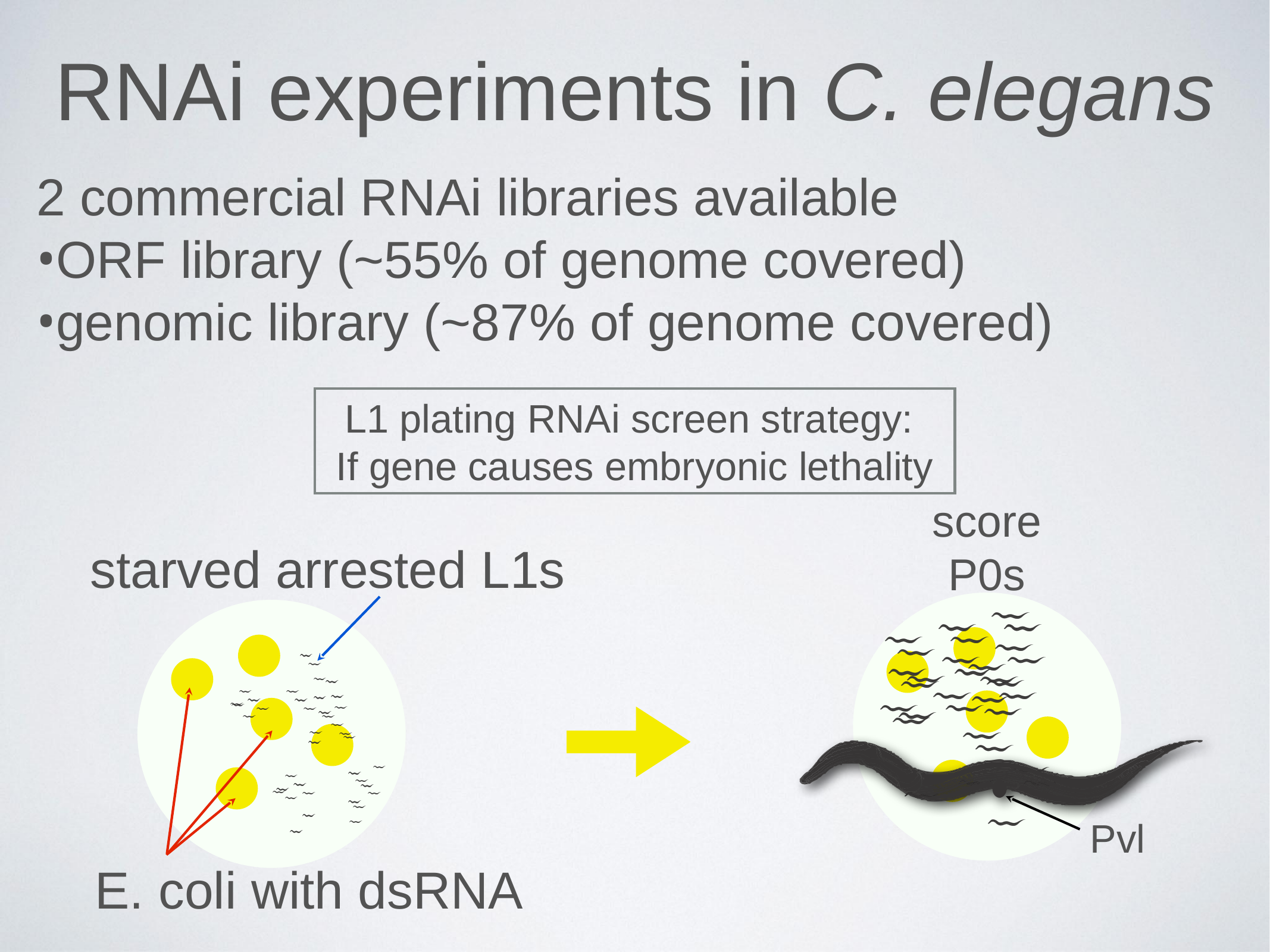

#### RNAi experiments in C. elegans
2 commercial RNAi libraries available
ORF library (~55% of genome covered)
genomic library (~87% of genome covered)
L1 plating RNAi screen strategy:
If gene causes embryonic lethality
score P0s
starved arrested L1s
E. coli with dsRNA
Pvl

### Slide 6
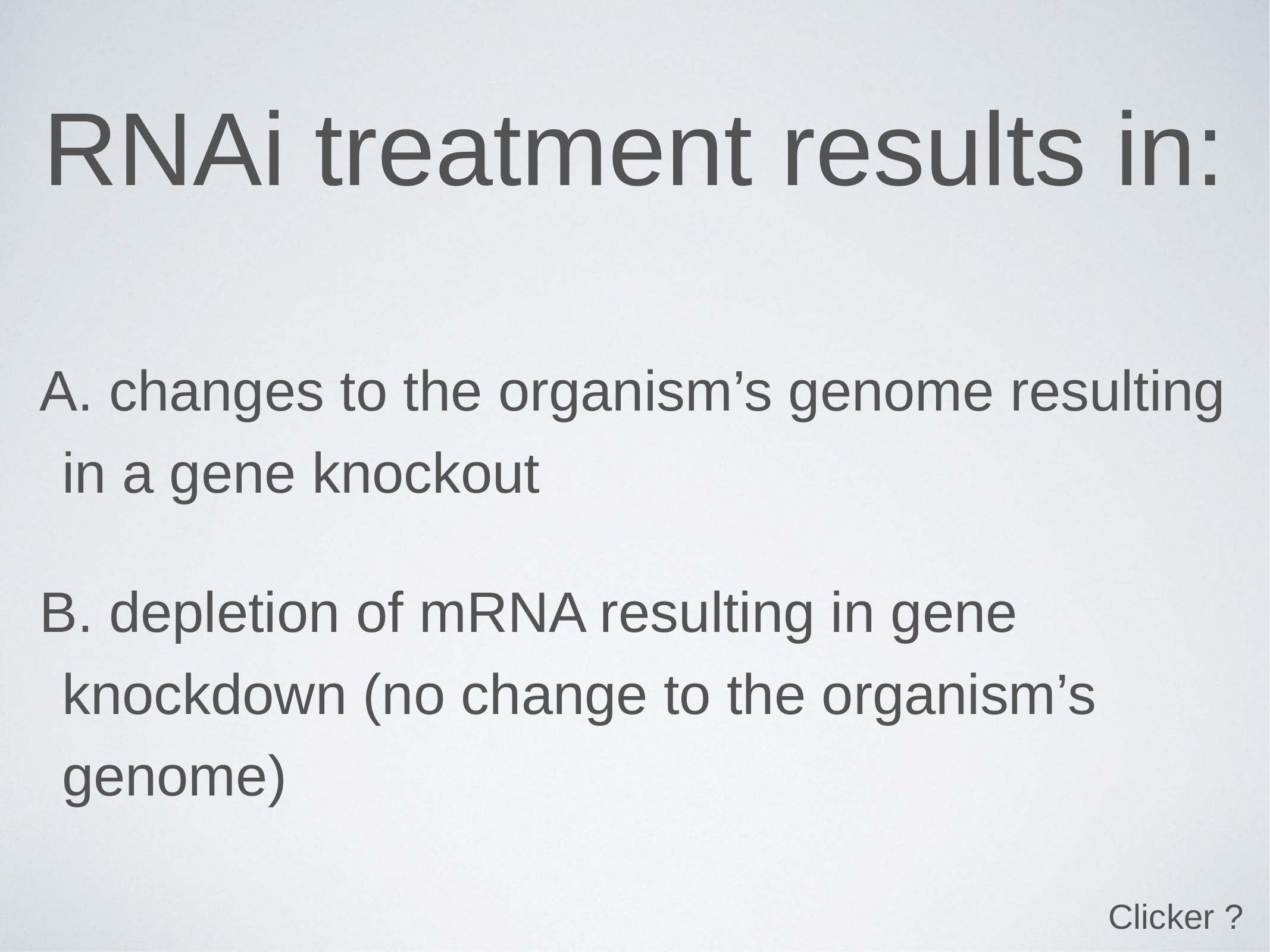

#### RNAi treatment results in:
 changes to the organism’s genome resulting in a gene knockout
 depletion of mRNA resulting in gene knockdown (no change to the organism’s genome)
Clicker ?

### Slide 7
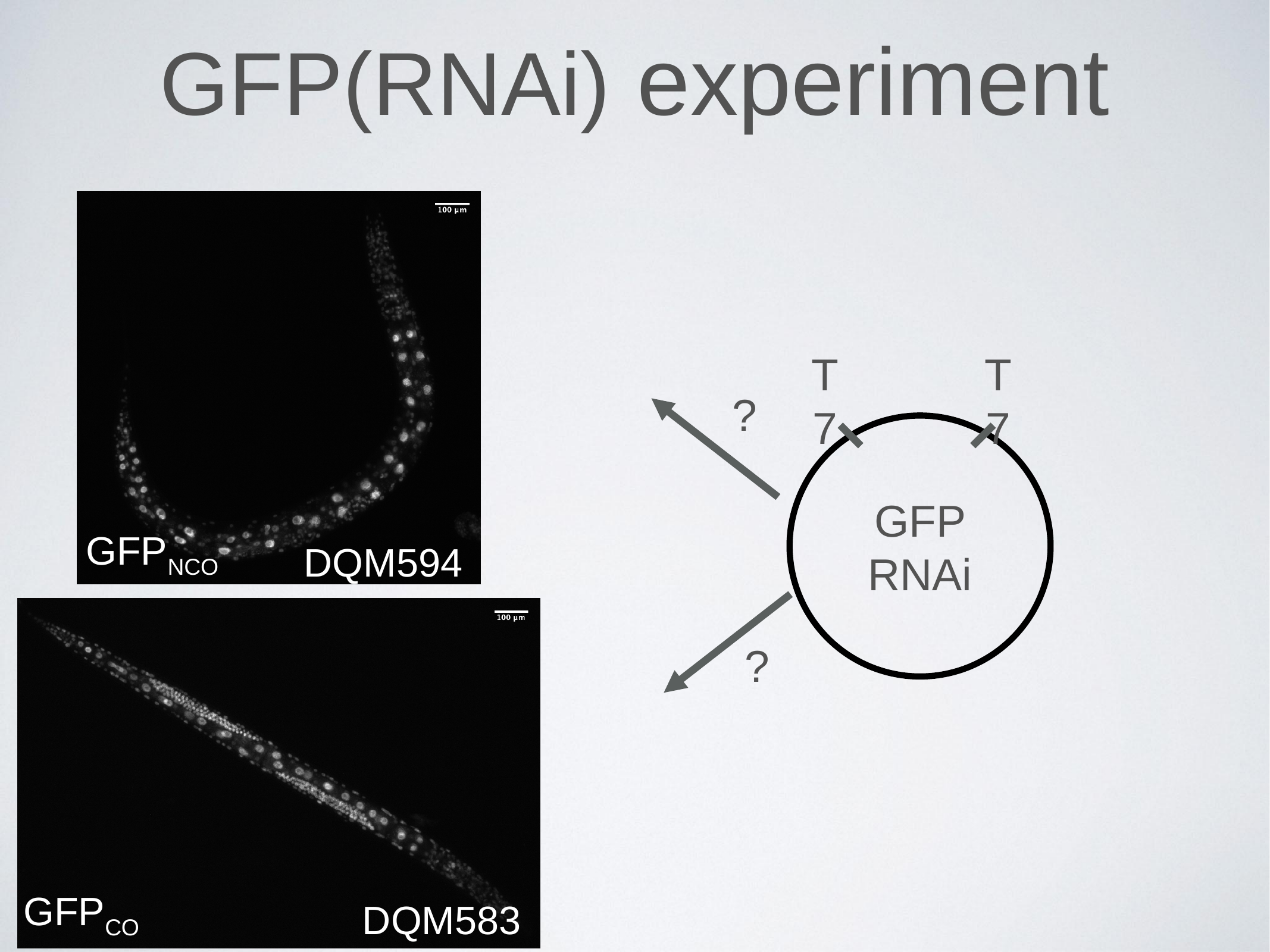

#### GFP(RNAi) experiment
T7
T7
?
GFP RNAi
GFP_NCO_
DQM594
?
GFP_CO_
DQM583

### Slide 8
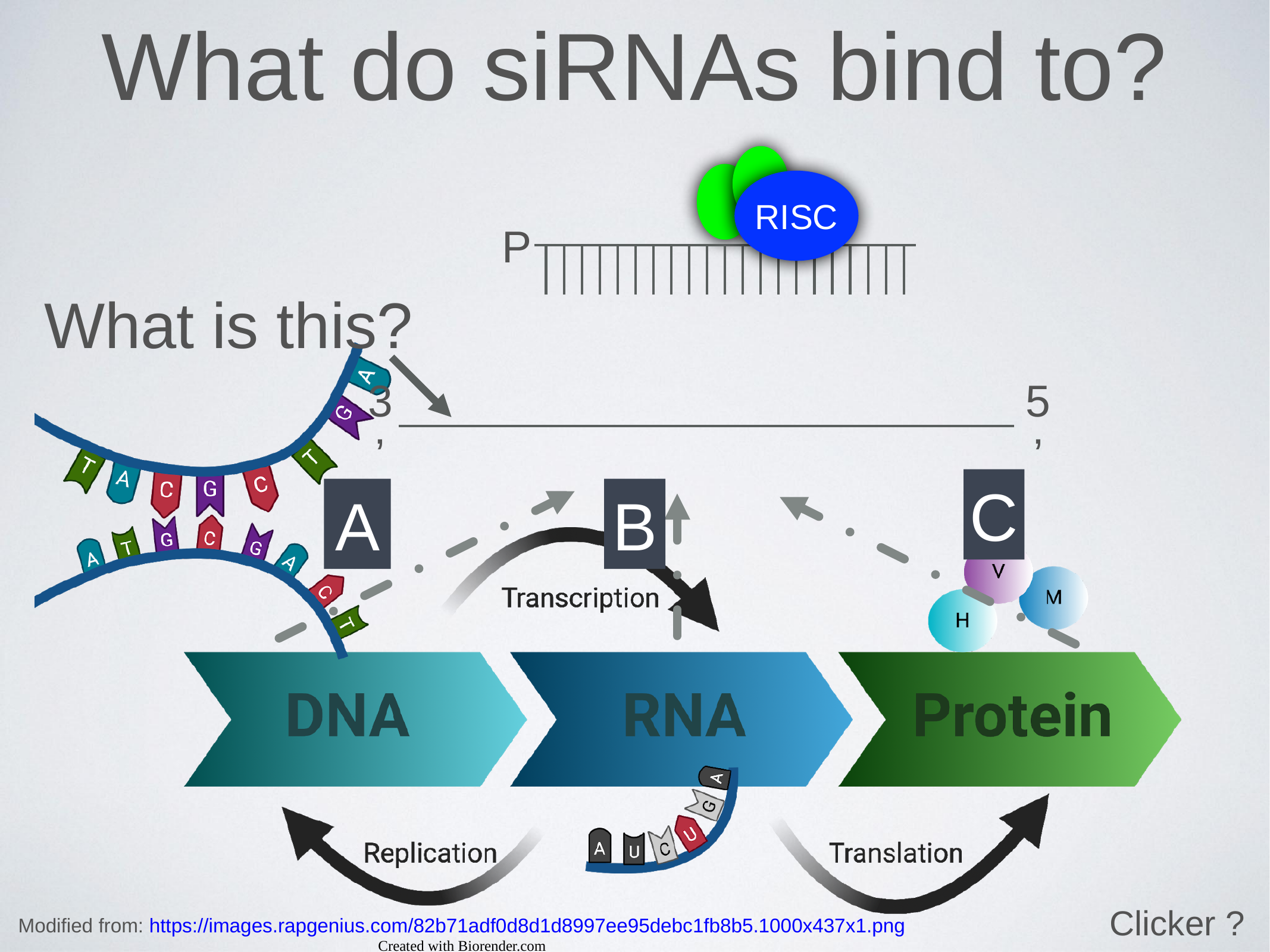

What do siRNAs bind to?
RISC
P
What is this?
3’
5’
C
A
B
Clicker ?
Modified from: https://images.rapgenius.com/82b71adf0d8d1d8997ee95debc1fb8b5.1000x437x1.png
Created with Biorender.com

### Slide 9
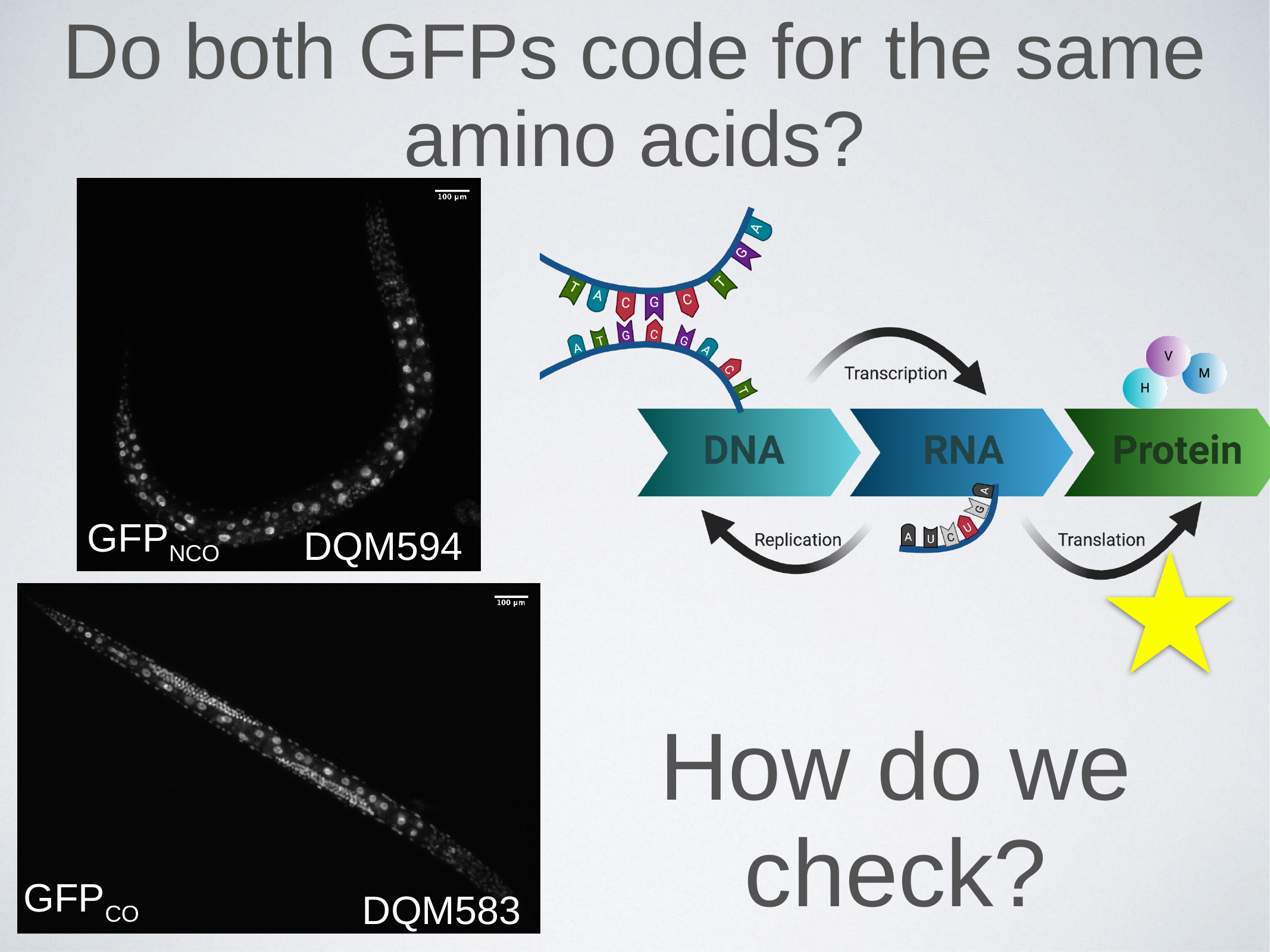

#### Do both GFPs code for the same amino acids?
GFP_NCO_
DQM594
How do we check?
GFP_CO_
DQM583

### Slide 10
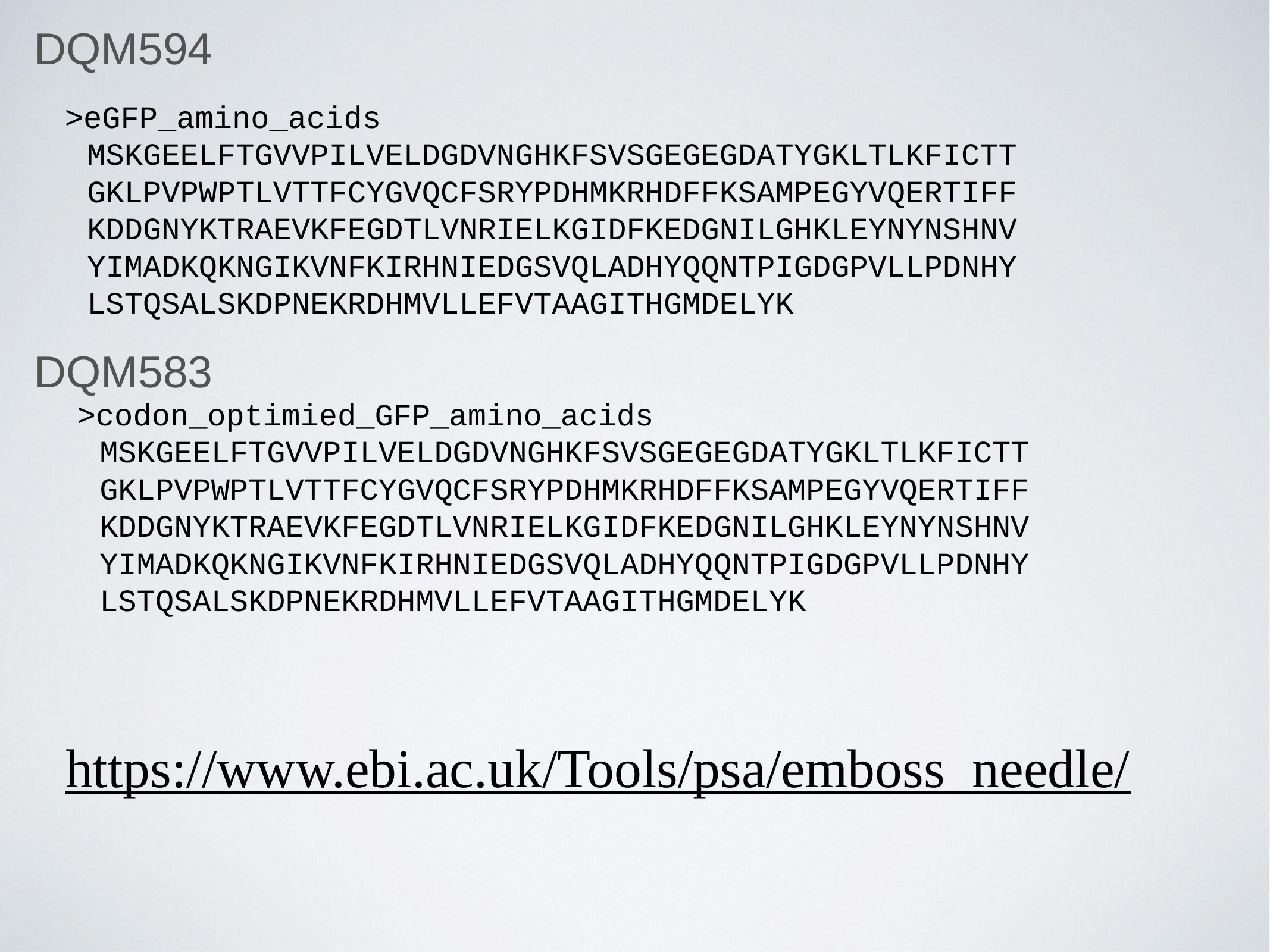

DQM594
>eGFP_amino_acids
	MSKGEELFTGVVPILVELDGDVNGHKFSVSGEGEGDATYGKLTLKFICTT
	GKLPVPWPTLVTTFCYGVQCFSRYPDHMKRHDFFKSAMPEGYVQERTIFF
	KDDGNYKTRAEVKFEGDTLVNRIELKGIDFKEDGNILGHKLEYNYNSHNV
	YIMADKQKNGIKVNFKIRHNIEDGSVQLADHYQQNTPIGDGPVLLPDNHY
	LSTQSALSKDPNEKRDHMVLLEFVTAAGITHGMDELYK
DQM583
>codon_optimied_GFP_amino_acids
	MSKGEELFTGVVPILVELDGDVNGHKFSVSGEGEGDATYGKLTLKFICTT
	GKLPVPWPTLVTTFCYGVQCFSRYPDHMKRHDFFKSAMPEGYVQERTIFF
	KDDGNYKTRAEVKFEGDTLVNRIELKGIDFKEDGNILGHKLEYNYNSHNV
	YIMADKQKNGIKVNFKIRHNIEDGSVQLADHYQQNTPIGDGPVLLPDNHY
	LSTQSALSKDPNEKRDHMVLLEFVTAAGITHGMDELYK
https://www.ebi.ac.uk/Tools/psa/emboss_needle/

### Slide 11
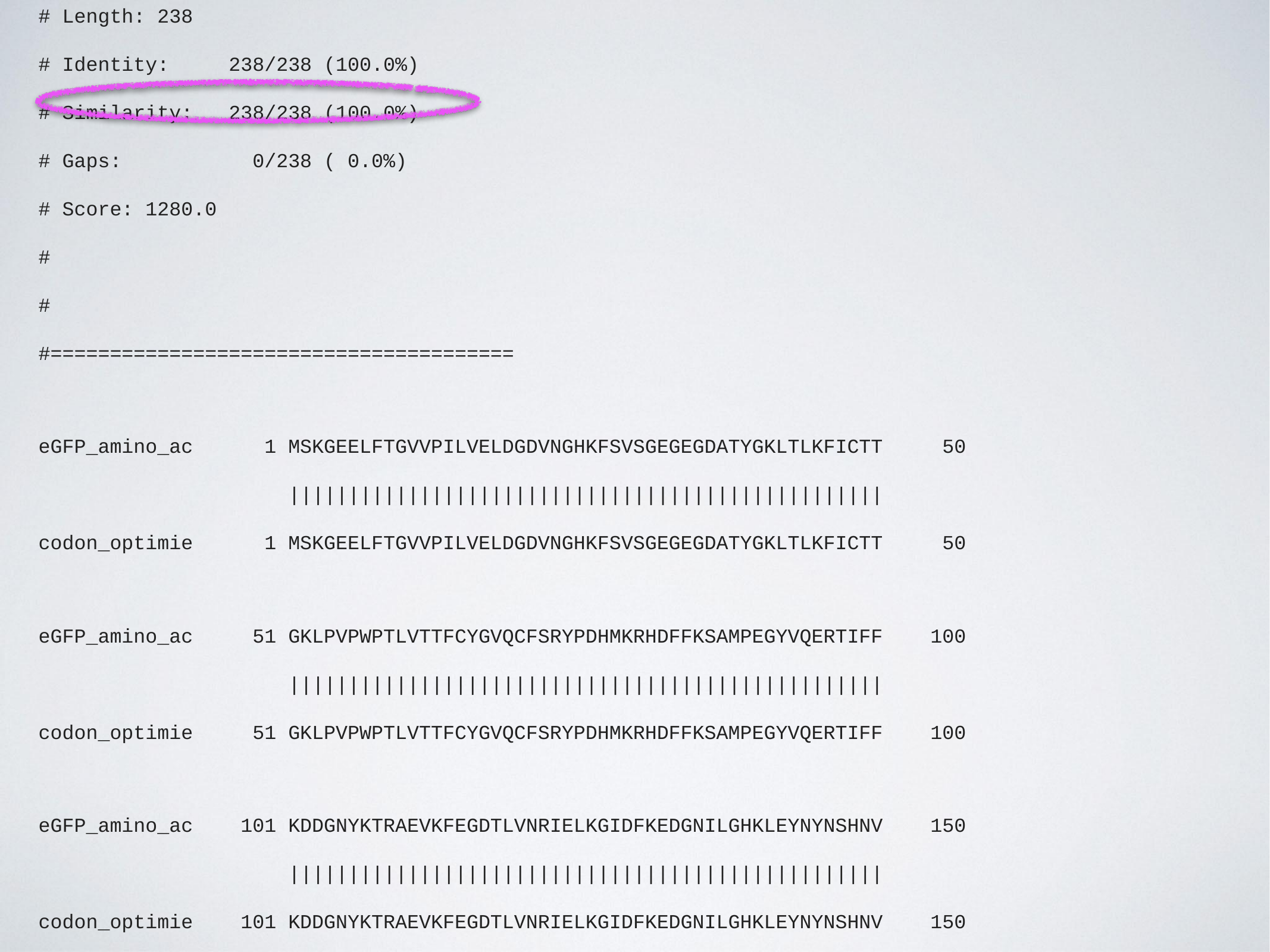

#### Aligned_sequences: 2
#### 1: eGFP_amino_acids
#### 2: codon_optimied_GFP_amino_acids
#### Matrix: EBLOSUM62
#### Gap_penalty: 10.0
#### Extend_penalty: 0.5
#
#### Length: 238
#### Identity: 238/238 (100.0%)
#### Similarity: 238/238 (100.0%)
#### Gaps: 0/238 ( 0.0%)
#### Score: 1280.0
#
#
#=======================================
eGFP_amino_ac 1 MSKGEELFTGVVPILVELDGDVNGHKFSVSGEGEGDATYGKLTLKFICTT 50
 ||||||||||||||||||||||||||||||||||||||||||||||||||
codon_optimie 1 MSKGEELFTGVVPILVELDGDVNGHKFSVSGEGEGDATYGKLTLKFICTT 50
eGFP_amino_ac 51 GKLPVPWPTLVTTFCYGVQCFSRYPDHMKRHDFFKSAMPEGYVQERTIFF 100
 ||||||||||||||||||||||||||||||||||||||||||||||||||
codon_optimie 51 GKLPVPWPTLVTTFCYGVQCFSRYPDHMKRHDFFKSAMPEGYVQERTIFF 100
eGFP_amino_ac 101 KDDGNYKTRAEVKFEGDTLVNRIELKGIDFKEDGNILGHKLEYNYNSHNV 150
 ||||||||||||||||||||||||||||||||||||||||||||||||||
codon_optimie 101 KDDGNYKTRAEVKFEGDTLVNRIELKGIDFKEDGNILGHKLEYNYNSHNV 150
eGFP_amino_ac 151 YIMADKQKNGIKVNFKIRHNIEDGSVQLADHYQQNTPIGDGPVLLPDNHY 200
 ||||||||||||||||||||||||||||||||||||||||||||||||||
codon_optimie 151 YIMADKQKNGIKVNFKIRHNIEDGSVQLADHYQQNTPIGDGPVLLPDNHY 200
eGFP_amino_ac 201 LSTQSALSKDPNEKRDHMVLLEFVTAAGITHGMDELYK 238
 ||||||||||||||||||||||||||||||||||||||
codon_optimie 201 LSTQSALSKDPNEKRDHMVLLEFVTAAGITHGMDELYK 238

### Slide 12
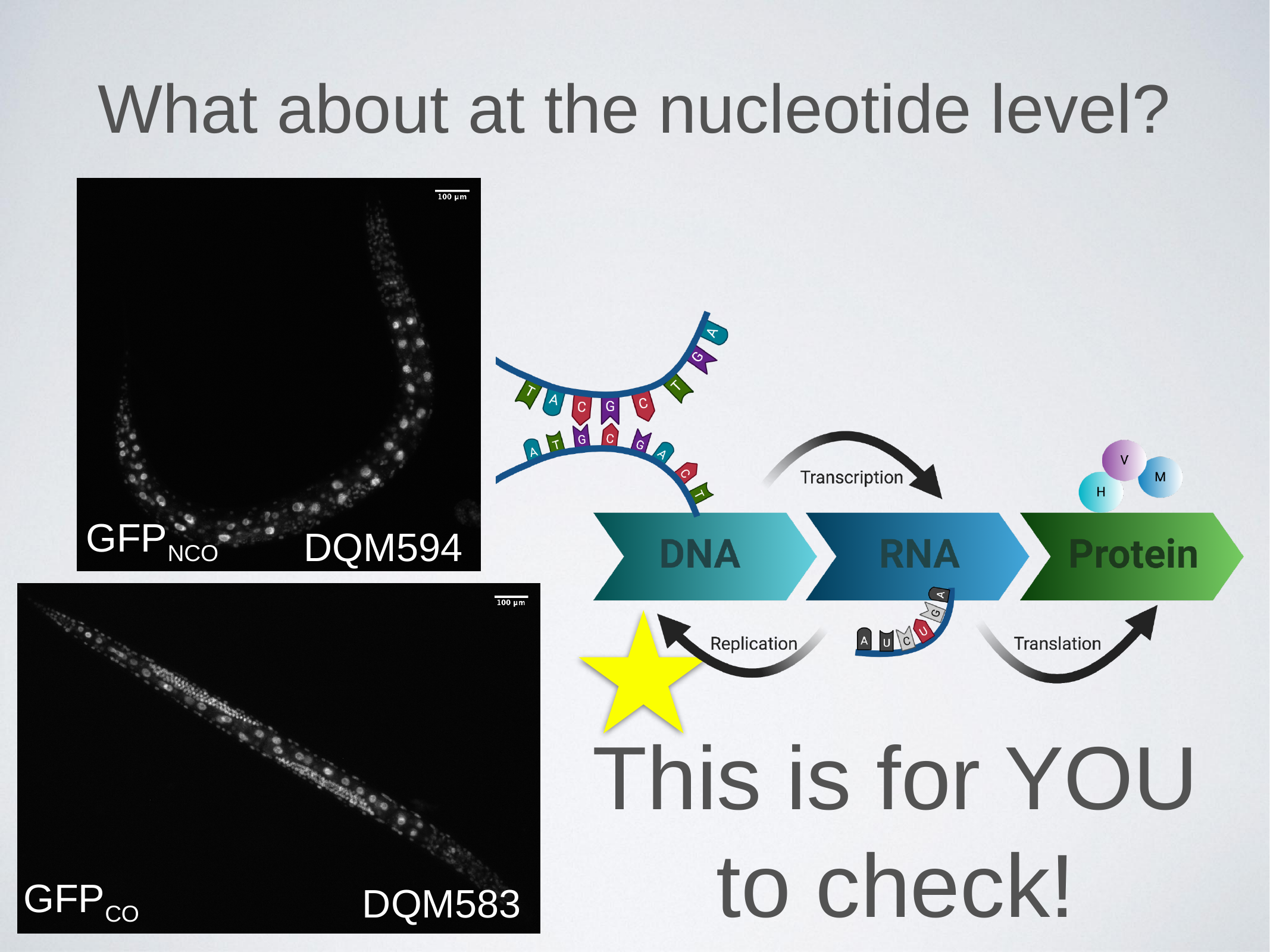

#### What about at the nucleotide level?
GFP_NCO_
DQM594
This is for YOU to check!
GFP_CO_
DQM583

### Slide 13
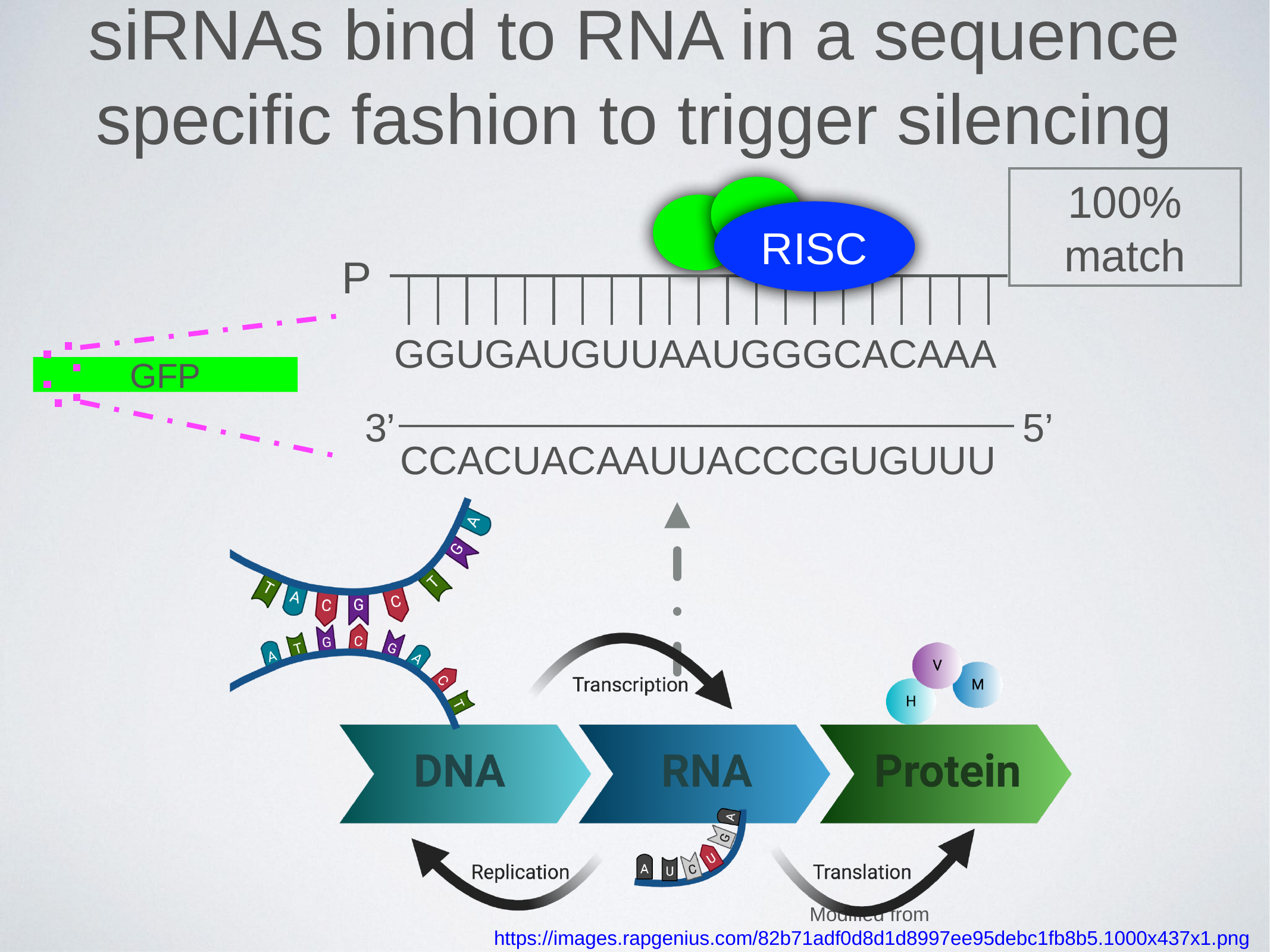

siRNAs bind to RNA in a sequence specific fashion to trigger silencing
100% match
RISC
P
GGUGAUGUUAAUGGGCACAAA
GFP
3’
5’
CCACUACAAUUACCCGUGUUU
Modified from https://images.rapgenius.com/82b71adf0d8d1d8997ee95debc1fb8b5.1000x437x1.png
Created with Biorender.com

### Slide 14
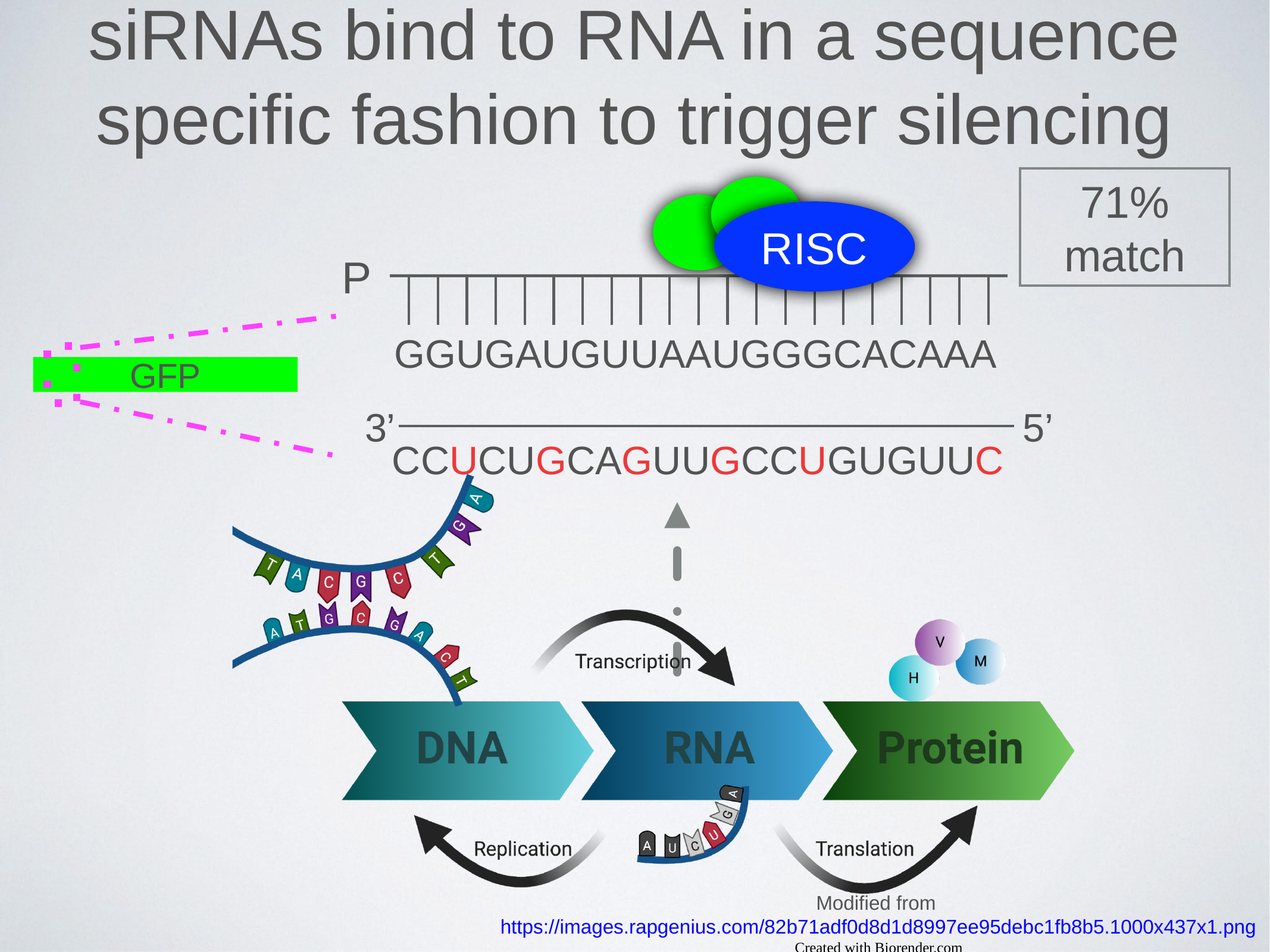

siRNAs bind to RNA in a sequence specific fashion to trigger silencing
71% match
RISC
P
GGUGAUGUUAAUGGGCACAAA
GFP
3’
5’
CCUCUGCAGUUGCCUGUGUUC
Modified from https://images.rapgenius.com/82b71adf0d8d1d8997ee95debc1fb8b5.1000x437x1.png
Created with Biorender.com

### Slide 15
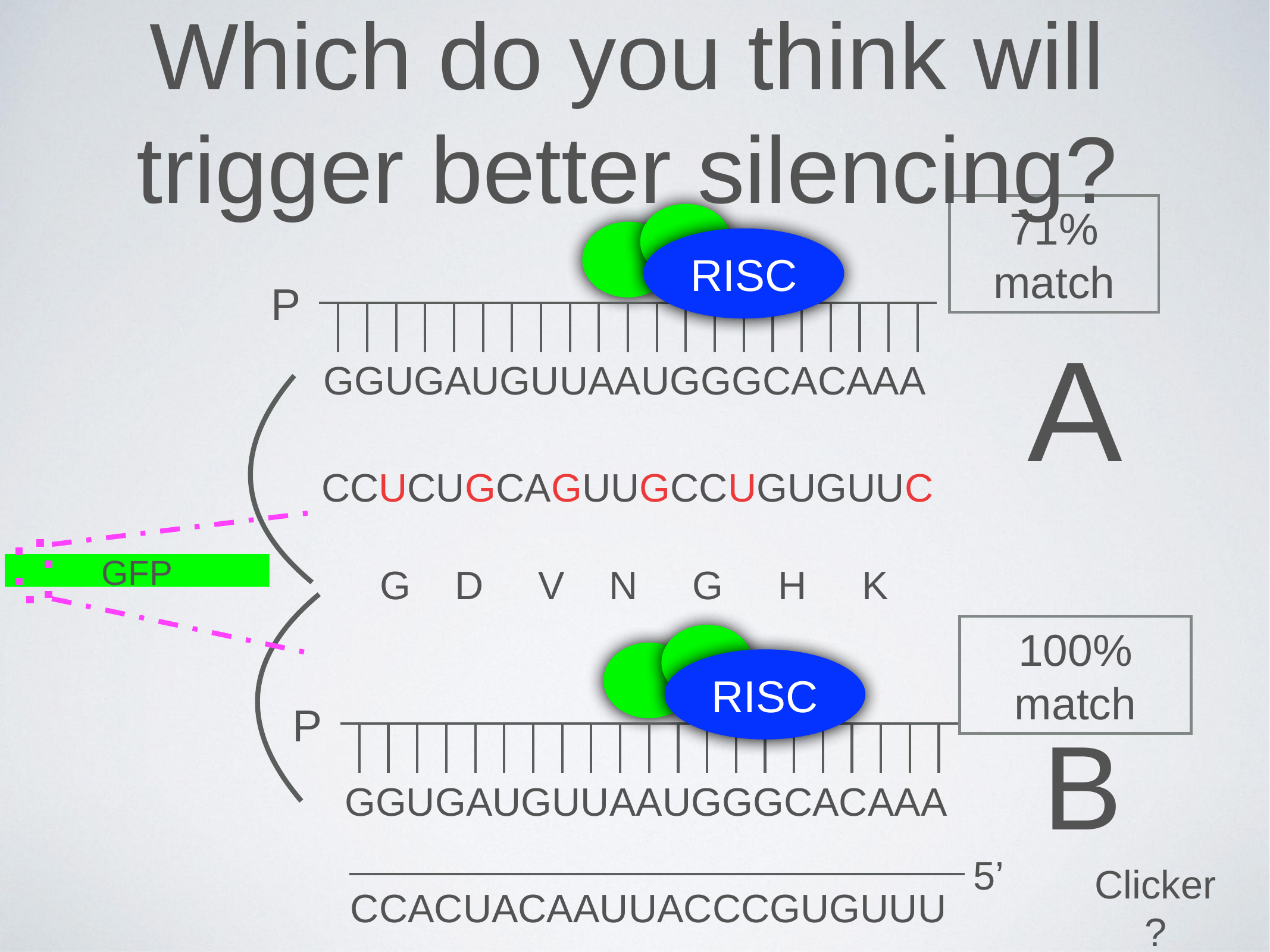

Which do you think will trigger better silencing?
71% match
RISC
P
GGUGAUGUUAAUGGGCACAAA
A
CCUCUGCAGUUGCCUGUGUUC
GFP
G D V N G H K
100% match
RISC
P
GGUGAUGUUAAUGGGCACAAA
B
5’
Clicker ?
CCACUACAAUUACCCGUGUUU

### Slide 16
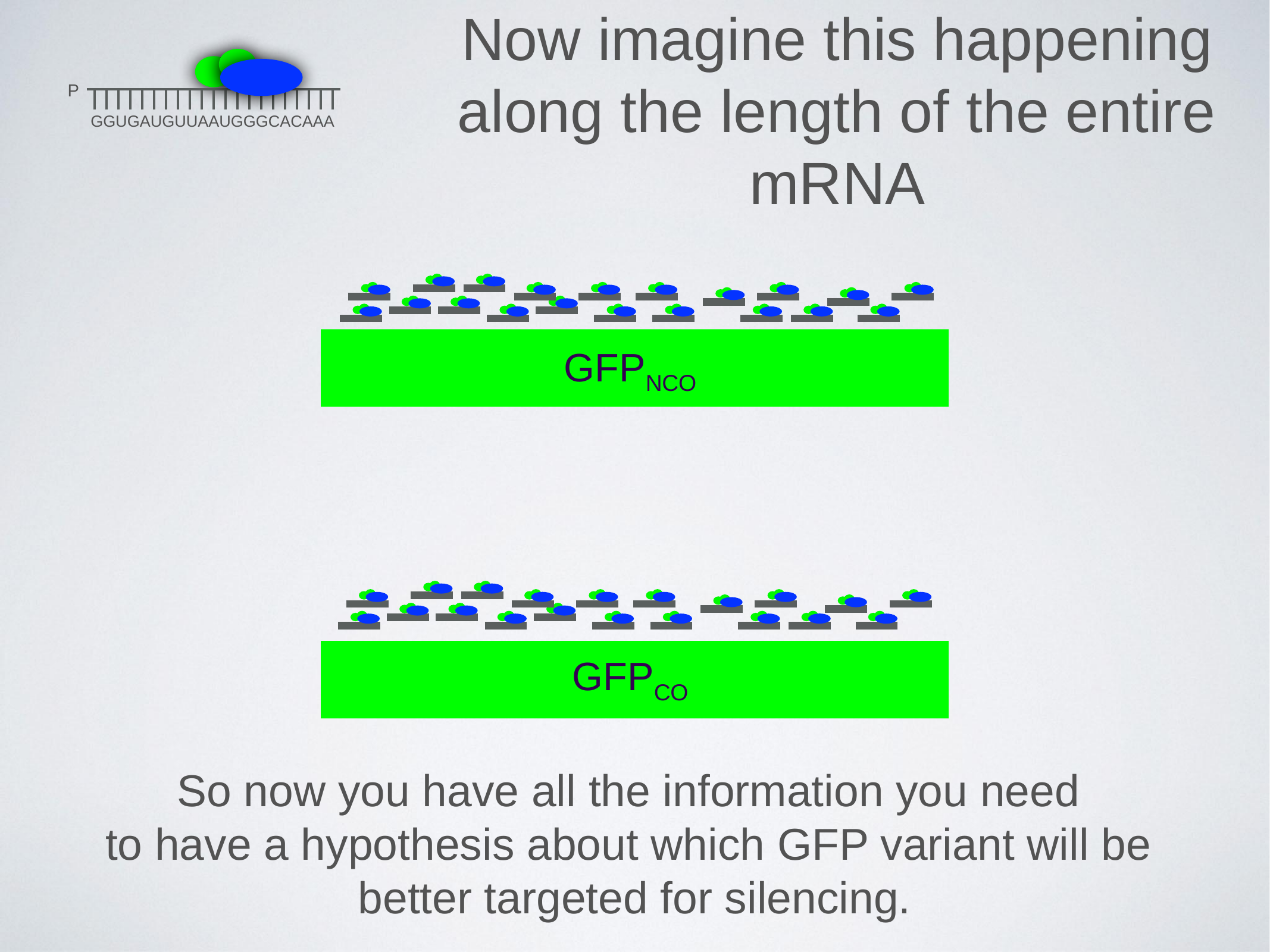

Now imagine this happening along the length of the entire mRNA
P
GGUGAUGUUAAUGGGCACAAA
GFP_NCO_
GFP_CO_
So now you have all the information you need
to have a hypothesis about which GFP variant will be
better targeted for silencing.
