## Supplemental File 2 for "A laboratory module that explores RNA interference and codon optimization through fluorescence microscopy using *Caenorhabditis elegans*"

**S2. A Laboratory Module-Grading Rubric and Example Lab Report**

| **Structure, weight, and criteria for judgment:** | | | |
| --- | --- | --- | --- |
| **Item** | **Rec. length** | **Weight** | **Criteria and comments** |
| **Results Figures:**   1. Amino acid and Nucleotide alignment figure comparing GFP sequences of DQM583 vs. DQM594 2. Images of GFP RNAi experiment & graphical representation of quantified data | 4 Figures + Results  Figure 1 (1/2 pg)  Figure 2 (1/2 pg) | 20 points total  Figure 1 (5 pts)  Figure 2 (5 pts) | -0.5 points for not following formatting  Each figure will have a legend. **Figure legends are a statement of what is contained in the figure.**  **Figure 1A-B.** Figure should show pairwise amino acid and nucleotide sequence alignments between the GFP sequence of DQM583 and DQM594.  **Figure 2A-D.** GFP RNAi experiment **micrographs** Representative image of DQM583 and DQM594 animals treated with control (T444T) and GFP RNAi (**4 images** from GFP channel (DIC/brightfield not necessary). Must include scale bar.)  **Figure 2E. Quantification of data as a graph:**  Dot plot (Plots of Data / R / Graph Pad/Excel) showing quantification of data, including **annotations showing significance and error bars showing standard deviation.** **Axes must be labelled with a font size that is easy to read!** |
| **Results Table**   1. Data table of quantification – highlight representative image | Table (1/3 pg) | Table (5 pt) | **Table 1:** Table should include data that was plotted (mean fluorescence intensity values for each imaging condition – 4 columns, labelled). Include the mean value for each data set and highlight which images were selected as representative images. |
| **Results Text**   1. Paragraph #2-3: GFP RNAi experimental results (reference Figure 1&2 and Table 1) | ~1/2 page | Paragraph 2 (5 pts) | Please describe the figures.  1. Why did you do the experiment? What was your hypothesis?  2. What were the results (describe the figures).  3. Summary sentence of what happened (what was your result?).  YOU MUST REFERENCE YOUR FIGURES IN THE RESULTS TEXT |
| **Acknowledgements** |  | 0 points | Acknowledge contributions from others |

**Detailed Outline of Lab Report with examples:**

**Figures and Results:**

**20 total points**

**Figures, Figure Legends, and Data Table:**

**15 points**

**Make sure to include scale bars and make sure you perform the required statistical tests so you can comment on the statistical significance in the figure legend where appropriate.**

**A**

```
# Aligned_sequences: 2
# 1: DQM583
# 2: DQM594
# Matrix: EBLOSUM62
# Gap_penalty: 10.0
# Extend_penalty: 0.5
# Length: 238
# Identity:     238/238 (100.0%)
# Similarity:   238/238 (100.0%)
# Gaps:           0/238 ( 0.0%)
# Score: 1280.0

DQM583      	    1 MSKGEELFTGVVPILVELDGDVNGHKFSVSGEGEGDATYGKLTLKFICTT     50
                     ||||||||||||||||||||||||||||||||||||||||||||||||||
DQM594      	    1 MSKGEELFTGVVPILVELDGDVNGHKFSVSGEGEGDATYGKLTLKFICTT     50

DQM583      	   51 GKLPVPWPTLVTTFCYGVQCFSRYPDHMKRHDFFKSAMPEGYVQERTIFF    100
                     ||||||||||||||||||||||||||||||||||||||||||||||||||
DQM594      	   51 GKLPVPWPTLVTTFCYGVQCFSRYPDHMKRHDFFKSAMPEGYVQERTIFF    100

DQM583      	  101 KDDGNYKTRAEVKFEGDTLVNRIELKGIDFKEDGNILGHKLEYNYNSHNV    150
                     ||||||||||||||||||||||||||||||||||||||||||||||||||
DQM594       	  101 KDDGNYKTRAEVKFEGDTLVNRIELKGIDFKEDGNILGHKLEYNYNSHNV    150

DQM583      	  151 YIMADKQKNGIKVNFKIRHNIEDGSVQLADHYQQNTPIGDGPVLLPDNHY    200
                     ||||||||||||||||||||||||||||||||||||||||||||||||||
DQM594      	  151 YIMADKQKNGIKVNFKIRHNIEDGSVQLADHYQQNTPIGDGPVLLPDNHY    200

DQM583      	  201 LSTQSALSKDPNEKRDHMVLLEFVTAAGITHGMDELYK    238
                     ||||||||||||||||||||||||||||||||||||||
DQM594      	  201 LSTQSALSKDPNEKRDHMVLLEFVTAAGITHGMDELYK    238
```

Figure 1**. Title.** (A) Amino acid alignment …(B) nucleotide alignment …

This can simply be a screen shot or print a PDF of the alignment you generate from ([<https://www.ebi.ac.uk/Tools/psa/emboss_needle/>](https://www.ebi.ac.uk/Tools/psa/emboss_needle/))

The above is an example using the amino acid alignment (make sure your figure also includes the nucleotide sequence alignment between DQM583 and DQM594)

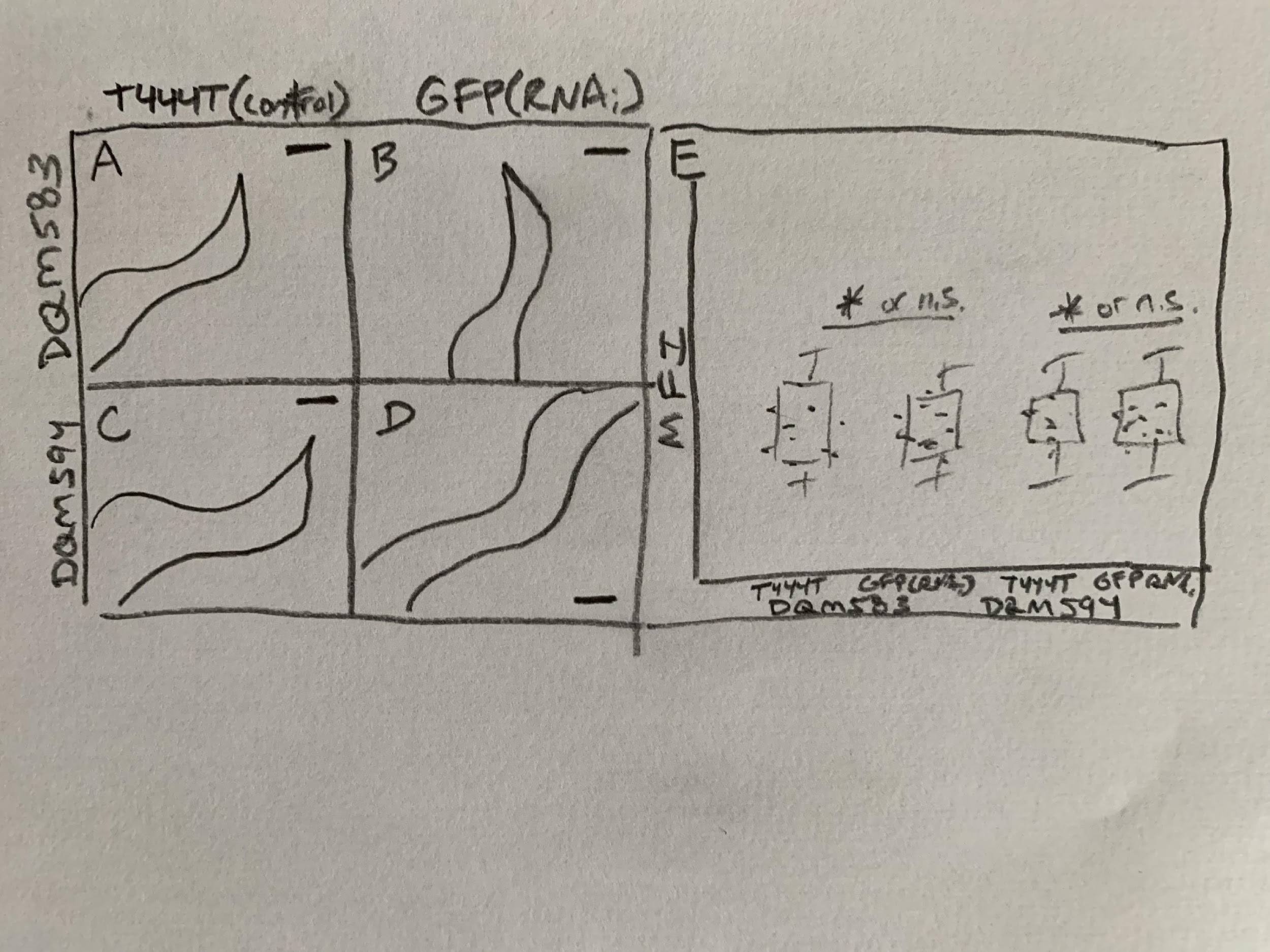

Figure 2. **Title**. (A)… (B)… (C)… (D)… (E) Plot… *, P=xxx, Student’s t-test or n.s., P=xxx, Student’s t-test.

Table 1. **Title**.

| **Image #** | **DQM583** | | **DQM594** | |
| --- | --- | --- | --- | --- |
|  | **T444T** | **GFP(RNAi)** | **T444T** | **GFP(RNAi)** |
| 1 | xxx | xxx | xxx | xxx |
| 2 | xxx | xxx | xxx | xxx |
| 3 | xxx | xxx | xxx | xxx |
| 4 | xxx | xxx | xxx | xxx |
| 5 | xxx | xxx | xxx | xxx |
| 6 | xxx | xxx | xxx | xxx |
| 7 | xxx | xxx | xxx | xxx |
| 8 | xxx | xxx | xxx | xxx |
| 9 | xxx | xxx | xxx | xxx |
| 10 | xxx | xxx | xxx | xxx |
| **Mean** | xxx | xxx | xxx | xxx |

Grey shaded boxes indicate images selected based on mean fluorescence intensity for Figure 2.

**Results (text): 5 points**

Presentation of results in text (refer to figures/tables/graphs appropriately and in order)

- **Paragraph 1-2** will describe the results of your GFP RNAi experiment. Based on your nucleotide alignment you should have a prediction of which strain (DQM583 or DQM594) should be more efficiently targeted by the GFP RNAi clone. Does your data confirm this result? **You must reference Figure 1, 2 and Table 1**. **Do not include a DISCUSSION of the results – this belongs in the DISCUSSION (which we are not including in this lab report)**

**Acknowledgements:**

0 points, but we will subtract points if data/images from others are used without giving credit.

**Example Lab Report:**
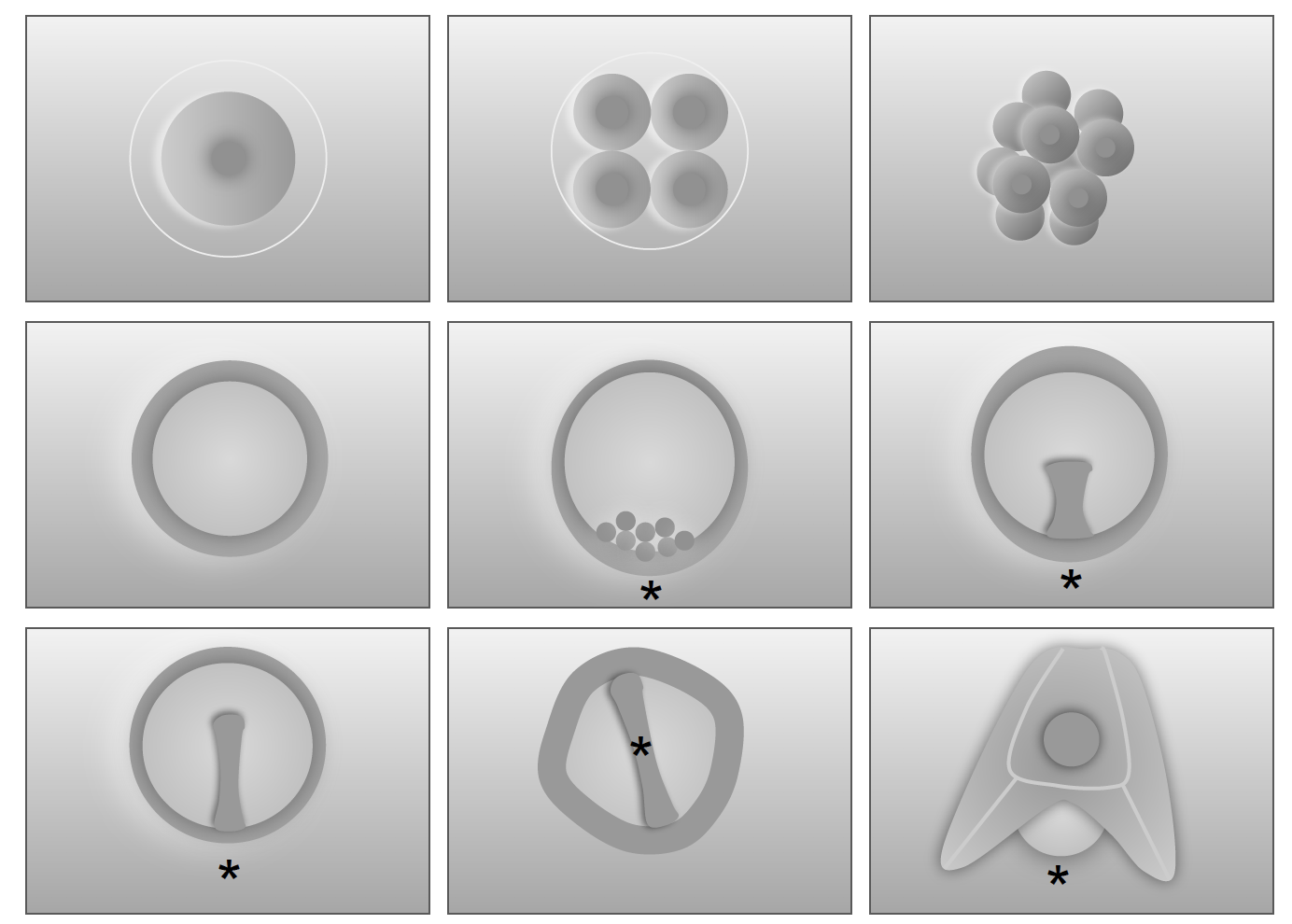

E

D

B

C

F

G

H

I

A

**Figure 1. Developmental series of the purple sea urchin, *Strongylocentrorus purpuratus*.** **A-C)** Cleavage stages showing radial cleavage pattern. **D)** Blastula stage. **E)** Onset of gastrulation, asterisk denotes vegetal pole and site of gastrulation, white arrowheads denote ingressing primary mesenchyme cells (PMCs). **F)** Mid-gastrulation. **G)** Late gastrulation. **H)** Formation of dipleurula larval stage. **I)** Dipleurula pluteus swimming larvae, arrow denotes biomineralized skeleton.

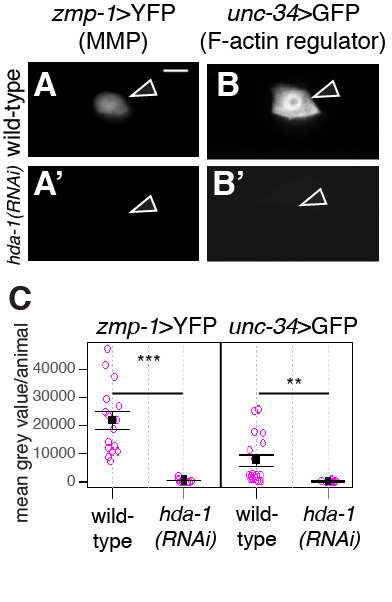

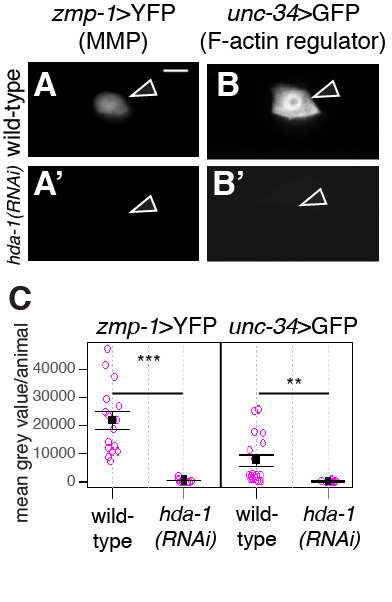

**Figure 2. *hda-1* regulates *zmp-1* and *unc-34* reporters.** Depletion of *hda-1* by RNAi results in loss of *zmp-1*>YFP (**A’**) and *unc-34*>GFP (**B’**) as compared to wild-type (**A-B**). Scale bar, 5 µm. **C)** Dot plot quantification of *zmp-1*>YFP (left) and *unc-34*>GFP (right) following *hda-1(RNAi)* treatment. ****P*<0.001, **P<0.01, Student’s t-test, error bar = standard deviation.

**Results**

**Developmental series of the purple sea urchin**

We followed the embryogenesis of the purple sea urchin, *Stongylocentrotus* purpuratus, from fertilization through development of the free swimming pluteus larvae (Fig. 1). Using DIC microscopy, we identified radial cleavage stages (Fig. 1A-C) and blastula formation (Fig. 1D). At approximately 17 hours post fertilization (hpf) we observed the process of gastrulation (Fig. 1E-G), initiated by the ingression of the primary mesenchyme cells (Fig. 1E). Following gastrulation, the pluteus larvae (Fig. 1H-I) was visualized, which enabled identification of morphological features specific to this life history stage, including the biomineralized skeleton (Fig. 1I).

**The histone deacetylase, *hda-1*, regulates MMPs and F-actin modifiers in the *C. elegans* anchor cell.**

Transgenic animals expressing YFP/GFP-tagged transcriptional reporters specific to the upstream regulatory regions of the matrix metalloproteinase (MMP), *zmp-1*, or the F-actin regulator, *unc-34*, were examined during the third larval stage of *C. elegans* post-embryonic development. (Fig. 2) YFP/GFP expression was visualized in the anchor cell (AC) in wild-type animals to establish baseline expression levels (Figure 2A-B). Depletion of the histone deacetylase, *hda-1*, by RNAi treatment, resulted in loss of the AC-specific expression of *zmp-1*>YFP (Fig. 2A’) and *unc-34*>GFP (Fig. 2B’). Quantification of this data identified *hda-1* as an upstream positive regulator of *zmp-1* (Fig. 1C, n ≥ 15 examined for each treatment, ***P<0.0001, Student’s t-test) and *unc-34* (Fig. 2C, n ≥ 15 examined for each treatment, **P<0.001), Student’s t-test) transcriptional activity in the AC during the L3 stage.
