## Supplemental File 3 for "A laboratory module that explores RNA interference and codon optimization through fluorescence microscopy using *Caenorhabditis elegans*"

### **S3. A Laboratory Module-GFP RNAi Module Worksheet Discussion Questions & Answers**

#### **Lab discussion questions for GFP RNAi worksheet(in-person/online):**

**Question** **1.** Based off the EMBOSS Needle alignment results from the GFP RNAi worksheet, which strain (GFP_NCO_ strain or GFP_CO_ strain) would the RNAi be most effective against? What would your hypothesis be?

**Answer:** The GFP_NCO_ strain – Despite the 100% identical amino acid alignments (shown in class), the GFP nucleotide sequences alignment between the GFP_NCO_ and GFP_CO_ strains were 71% similar. Comparing each strain’s GFP DNA sequence with the RNAi sequence provided, the NCO GFP sequence was more similar (or 97.9% similar (97.9% identical)) to the GFP RNAi sequence than the CO GFP sequence.

Possible alternative correct hypotheses:

- The GFP_NCO_ strain (DQM594) would be targeted more efficiently by the GFP_NCO_ RNAi for GFP knock-down.
- The GFP_NCO_ strain will have little/no GFP expression after RNAi treatment.
- The GFP_CO_ strain (DQM583) would have less/no GFP knock-down after RNAi treatment.
- The GFP_CO_ strain (DQM583) will have more GFP expression than the GFP_NCO_ strain after RNAi treatment.

**Question** **2.** From your observations using the fluorescence microscopes in lab, are there any differences in fluorescence intensity between the GFP_CO_ and the GFP_NCO_ strains fed on bacteria containing empty vector control?

**Answer:**

1) Yes – The GFP_CO_ strain is brighter under the same exposure/brightness and contrast settings compared to the GFP_NCO_ strain, and the germline expression of GFP in the GFP_CO_ strain is visible unlike in the GFP_NCO_ strain.

**Question** **3.** With respect to question 2, what do you suppose gives rise to these differences between the two strains?

**Answer:** The differences in fluorescence intensity seen between both strains are due to differences in the codon usage bias between codon optimized GFP and non-codon optimized GFP.

**Question** **4.** From your observations using the fluorescence microscopes in lab, are there any differences in fluorescence intensity between the GFP_CO_ and the GFP_NCO_ strains fed on GFP_NCO_ RNAi?

**Answer:** Yes – the GFP_NCO_ strain was most affected by GFP_NCO_ RNAi because its nucleotide sequence has better homology to GFP_NCO_ than GFP_CO_.

**Question** **5.** Does your hypothesis of whether the GFP_NCO_ RNAi clone will better deplete GFP_NCO_ or GFP_CO_ match the result you obtained from your experiment? Why or why not?

The answer to this question is variable and instructors or teaching assistants will have to guide the students here depending on their answer. Below are answer categories with examples:

- **Excellent:** Student clearly and accurately states the hypothesis and how it relates to the result.

Example: We predicted that GFP_NCO_ RNAi will significantly knockdown the fluorescence intensity of the GFP_NCO_ strain since their sequences match almost 97%. Our observations while conducting the experiment support our hypothesis since the fluorescence intensity of the GFP_NCO_ strain is diminished when fed with the GFP_NCO_ RNAi.

- **Good:** Student provides a partial answer and needs prompting to relate either the hypothesis to the result or vice versa.

Example: There is no visible difference between the GFP_CO_ strain when fed with the control or the GFP_NCO_ RNAi. Whereas there is a visible difference in the GFP_NCO_ strain after the GFP_NCO_ RNAi treatment.

- **Weak:** Student proposes a wrong answer or cannot build a connection between the results and the experiment.

Example: Our hypothesis is that the GFP_NCO_ RNAi clone significantly knocks down the fluorescence intensity of the GFP_CO_ strain.
