## Supplemental File 4 for "A laboratory module that explores RNA interference and codon optimization through fluorescence microscopy using *Caenorhabditis elegans*"

**S4. A Laboratory Module-Detailed Protocols**

***These protocols were modified from Martinez and Matus, 2020(1) and Stiernagle, 2006(2)**

**I. Preparing stock solutions:**

Materials:

- CaCl_2_
- Carbenicillin
- Cholesterol
- ddH_2_O
- 100% ethanol
- KH_2_PO_4_
- K_2_HPO_4_
- KOH
- MgSO_4_
- NaCl
- Levamisole
- Na_2_HPO_4_
- Glass storage bottles
- pH meter

Equipment:

- Autoclave
- Hot plate
- Magnetic stirring bar

Protocol for instructor(s):

1. 5 mg/mL cholesterol stock solution:
   a. Dissolve 2.5 g of cholesterol in 500 mL of 100% ethanol.
   b. Mix at 30°C for 15-30 minutes to dissolve and store at room temperature.
2. 1 M CaCl_2_ stock solution:
   a. Dissolve 73.5 g of CaCl_2_ in 500 mL of ddH_2_O.
   b. Autoclave for 30 minutes.
   c. Store at room temperature.
3. Carbenicillin stock solution:
   a. Dissolve 100 mg of carbenicillin in 1 mL of ddH_2_O.
   b. Aliquot 1 mL in 1.5 mL microcentrifuge tubes and store at -20°C.
4. IPTG stock solution:
   a. Dissolve 2 g of IPTG in 8 mL of ddH_2_O.
5. 1 M KPO_4_ stock solution:
   a. Dissolve 68.0 g of KH_2_PO_4_ in 500 mL of ddH_2_O to obtain a 1M KH_2_PO_4_ solution.
   b. Dissolve 52.3 g of K_2_HPO_4_ in 300 mL of ddH_2_O to obtain a 1M K_2_HPO_4_ solution.
   c. While measuring the pH of 1 M KH_2_PO_4_, carefully add and stir in 1 M K_2_HPO_4_ slowly until the pH rises from 4.0 to 6.0.
   d. Store at room temperature.
6. 5 M KOH stock solution:
   a. Dissolve 280.55 g of KOH in 1 L of ddH_2_O.
   b. Aliquot 200 mL into 500 mL storage bottles and store at room temperature.
7. Luria-Bertani broth (LB)
   a. Dissolve 10 g of NaCl, 10 g of tryptone, and 5 g of yeast extract in 1 L of ddH_2_O.
8. 1 M MgSO_4_ stock solution:
   a. Dissolve 60.2 g of MgSO_4_ in 500 mL of ddH_2_O and autoclave for 30 minutes.
   b. Store at room temperature.
9. M9 Minimal Medium buffer:
   a. Add 3 g of KH_2_PO_4_, 6 g of Na_2_HPO_4_, and 5 g of NaCl to 1 L of ddH_2_O.
   b. Aliquot 200 mL into 500 mL storage bottles and autoclave for 30 minutes.
   c. Add 200 μL of 1M MgSO_4_ to each bottle.
10. 1 M Levamisole stock solution:
    a. Dissolve 2.407 g of levamisole in 10 mL of M9 buffer.
    b. Filter sterilize solution and aliquot into 1.5 mL microcentrifuge tubes.
    c. Store at 4°C for up to 1-2 weeks.

**II. Preparing and seeding nematode growth media (NGM) plates:**

Materials:

- ddH_2_O
- NaCl
- Peptone
- Bacteriological agar
- 5 mg/mL cholesterol
- 1 M CaCl_2_
- 1 M MgSO_4_
- 1 M KPO_4_, pH 6.0
- Luria-Bertani broth
- OP50 *E. coli* (available from Caenorhabditis Genetics Center)

Equipment:

- Serological pipette & tips
- Magnetic stir bar
- Hot plate
- Water bath (optional)
- 4°C storage

Protocol for instructor(s):

1. Prepare NGM plates: This will make ~200 plates
   a. Add 6 g of NaCl, 7 g of peptone, 34 g of bacteriological agar, 2 mL of 5 mg/mL cholesterol, and a magnetic stirring bar to 1,944 mL of ddH_2_O in a 4 L Erlenmeyer flask.
   b. Autoclave for 55 minutes.
   c. Let the NGM cool to 55-60°C (a water bath set to 55-60°C can be used for this purpose).
   d. Add 2 mL of 1 M CaCl_2_, 2 mL of 1 M MgSO_4_, and 50 mL of 1 M KPO_4_, pH 6.0.
   e. Thoroughly mix the medium in a flask on a stir plate.
   f. Dispense 8 mL per 60 x 15 mm petri dish.
   g. Allow agar-filled petri dishes (plates) to solidify for 48 hours and store at room temperature.
2. Seed plates:
   a. Set up overnight culture of OP50 *E. coli* in LB.
   b. Pipette 200-300 μL of OP50 using a serological pipette.
   c. Allow plates to dry at room temperature for approximately 48 h. Store at 4°C and warm plates up at room temperature before use.

**III. *C. elegans* maintenance:**

Materials:

- Glass Pasteur pipette
- Glass cutter
- Pliers
- 90% platinum, 10% iridium wire (0.010 inches in diameter)
- Pencil grips (optional)
- NGM plates seeded with OP50 *E. coli*
- Permanent markers
- 15 mL or 50 mL conical tube
- 100% ethanol
- Rubber bands
- DQM594, non-codon-optimized GFP strain (available from Caenorhabditis Genetics Center)
- DQM583, codon-optimized GFP strain (available from Caenorhabditis Genetics Center)

Equipment:

- Alcohol burner or Bunsen burner
- Lighter or striker
- Metal laboratory spatula
- Stereo microscope
- Incubator set between 15-25°C (optional)

Protocol for instructor(s):

1. Assemble worm picks:
   a. Using a glass cutter, score and carefully break the glass where the Pasteur pipette reaches its narrowest point.
   b. Heat the cut edge over a flame until the glass begins to melt. Rotate Pasteur pipette as it melts such that the hole closes.
   c. While the glass is still hot, carefully press an approximately 1 inch long piece of 90% platinum/10% iridium wire into the glass using pliers Note: Use caution during this step to prevent injury.
   d. Let the glass cool. Using pliers, shape the wire into a hockey stick shape. Optional: add a pencil grip to the Pasteur pipette.
2. Assemble spatula setup:
   a. Place a metal laboratory spatula in an empty 15 mL or 50 mL conical tube.
   b. Fill the tube with 100% ethanol to cover the flat edge of the spatula. Note: Do not overfill the tubes with ethanol to prevent students from burn-related injuries.

Protocol for students:

1. Ensure that plates are labeled with the strain designation and date before any animals are deposited onto them. Note: Be sure to label the plate itself instead of the lid, as lids can get switched accidentally.
2. Flame sterilize your worm pick by heating the platinum wire over a Bunsen burner flame for 1-2 seconds. Wait another 1-2 seconds for the platinum to cool before attempting to pick up any animals. Note: Your worm pick will need to be flame-sterilized before handling each new strain to prevent cross-contamination.
3. Under a stereo microscope, pick up some bacteria with your worm pick (to act as an adhesive) and carefully scoop up the animals you would like to move to either a microscope slide (See section VI. slide making below) or a new plate.
4. Plates can be secured using a rubber band and should be stored lid side down at the desired temperature.

**IV. Preparing RNA interference (RNAi) plates**

IPTG/Carb plates can be made and stored at 4°C for up to three weeks prior to seeding with bacteria. Once seeded with bacteria can be stored for one week at 4°C

Materials:

- Luria-Bertani broth (LB)
- Carbenicillin
- IPTG
- T444T empty vector control bacteria (available from Addgene)
- Codon-optimized GFP RNAi bacteria (available from Addgene)
- Non-codon-optimized GFP RNAi bacteria (available from Addgene)

Equipment:

- P200 pipette
- P20 pipette
- 1-200 µL micropipette tips
- Tube shaker
- 37°C incubator

Protocol for instructor(s):

1. Transform 1 uL of RNAi bacterial plasmid into competent HT115 using the New England BioLabs transformation protocol (https://www.neb.com/protocols/2012/05/21/transformation-protocol).
2. Prepare IPTG & carbenicillin treated plates: This will make ~200 plates
   a. Prepare NGM media as described above.
   b. Add 1.875 mL of 0.8 M IPTG and 2 mL of carbenicillin (100 mg/mL) into 2 L of NGM media.
   c. Dispense 8 mL per 60 x 15 mm petri dish.
   d. Allow agar-filled petri dishes (plates) to solidify at room temperature for 48 hours and store at 4°C.
3. Seed NGM plates with RNAi bacteria:
   a. Set up an overnight culture of RNAi bacteria in 3-7 mL of LB broth treated with carbenicillin (1:500). Note: Carbenicillin can be added to each tube individually or can be added to a stock of LB (in this case it should be stored at 4°C for up to a month).
   b. Add 1:500 IPTG to the RNAi culture tubes (e.g., 14 μL of IPTG to 7 mL of LB).
   c. Incubate with shaking at 37°C for 1 hour.
   d. After 1 hour, seed RNAi plates with 200 μL of the bacterial liquid culture using a micropipette, and allow plates to dry overnight.
   e. Once dry, L1 larvae can be placed onto the plates and incubated at the desired temperature.

**V. Synchronization of *C. elegans* in development:**

Materials:

- P1000 pipette
- 15 mL conical tubes
- ddH_2_O
- 5M NaOH or KOH
- Bleach (3% sodium hypochlorite)
- M9 buffer

Equipment:

- Tube rocker
- Clinical centrifuge

Protocol for instructor(s):

1. Prepare 6 plates per group that have many gravid adults. This can be done through chunking.

Protocol for students:

1. Identify plates (ideally 6) that contain many gravid adults.
2. Add 1 mL of ddH_2_O to each plate. Swirl the liquid around the plates and/or use your pipette to resuspend and then remove animals from the bacteria. Tilt the plates and transfer the liquid to a 15 mL conical tube. Repeat with all of the remaining plates.
3. Once worms are collected into a conical tube, add ddH_2_O to a final volume of 8 mL.
4. To each conical tube, add 600 µL of 5M base (NaOH or KOH), followed by 1200 µL (600 µL x 2) of bleach (3% sodium hypochlorite).
5. Incubate the tube on a rocker for approximately 5-7 minutes. After 5 minutes, inspect the tube to ensure all adult bodies have dissolved. If adult bodies still remain, incubate in bleach solution until all adults have completely dissolved. Note: This step is very time sensitive – bleaching for too short a time will result in undissolved adults, whereas bleaching for too long will dissolve the eggs.
6. Spin down the 15 mL conical tube in a clinical centrifuge at approximately 2000-2500 rpm for 2 minutes. Be sure to balance the centrifuge with a conical tube containing an equal volume of water, if necessary.
7. After spinning is complete, discard the supernatant (alkaline bleach solution), leaving a pellet of eggs toward the bottom of the tube. Note: This pouring should be done in one motion, as repetitive pouring can disrupt the pellet of eggs.
8. Fill the tube with water (up to the 15 mL mark). Start by adding a small volume, flicking the tube to resuspend the pellet, before filling to the top. Spin down the tube once more at 2000-2500 rpm for 2 minutes and discard the supernatant.
9. Repeat the water wash step once more.
10. In the same fashion as the water wash steps (steps 9 and 10), perform two washes with M9 buffer.
11. After the final wash with M9 buffer, resuspend with approximately 2-3 mL of M9 buffer and incubate on the rocker.
12. The eggs will hatch and arrest in the L1 stage of development due to starvation conditions. You can plate these worms 1-2 days after bleaching.

**VI. Slide making**

Materials:

- 5% Noble agar
- P2 pipette
- M9 buffer
- Laboratory labeling tape
- Plastic or glass Pasteur pipettes
- Levamisole or Nemagel solution

Equipment:

- Heat block
- Stereo microscope

Protocol for instructor(s):

1. Prepare 5% Noble agar:
   a. Add 5 g of Nobel agar to 100 mL of ddH_2_O in a 250 mL storage bottle.
   b. Microwave until the agarose is completely dissolved (solution will become translucent).
2. Set temperature of heat block to 62°C and place the tubes of 5% Noble agar inside.

Protocol for students:

1. Generate a slide making station by taking two slides and placing two layers of laboratory tape on each (refer to supporting file S3).
2. Arrange these slides vertically side-by-side and place a third slide (without tape) in between them.
3. Pipette a small drop of molten agar onto the slide without tape.
4. Place a fourth slide perpendicular across the setup quickly (before the agar solidifies) and press down on either side of the agar pad.
5. Wait until the agar pad solidifies and then separate the slides just before use, as the agar pad will begin to dry out once exposed to air.
6. To the center of the agar pad, add a droplet (~5µL) of M9 buffer containing 5mM Levamisole, and add worms to this droplet. Finish by adding a coverslip on top of the slide. As an alternative to Levamisole, surround an M9 droplet (without any levamisole) with cooled (4°C) NemaGel solution and add worms to the M9 droplet. Place a droplet of NemaGel solution onto a coverslip and apply the coverslip to the slide.

**VII. Image quantification**

Materials:

- Computer with Fiji image processing package (<https://imagej.net/Fiji>)

Protocol for instructor(s):

1. Upload students’ images to a shared folder (e.g., Google Drive).

Protocol for students:

1. Open the FIJI application.
2. Open image file(s) in FIJI by navigating to **File > Open...**
3. Adjust the image brightness and contrast by navigating to **Image > Adjust > Brightness/Contrast…** Note: Be sure NOT to click “Apply” or you will change the pixel values of the image.
4. Specify the measurements you would like to collect by navigating to **Analyze > Set Measurements…**, selecting **Area** and **Mean gray value** and pressing **OK**. Note: this only needs to be done once.
5. Select the **Freehand selections** tool from the toolbar and draw around your region of interest (ROI).
6. Navigate to **Analyze > Measure** to measure the area and mean gray value of the ROI.
7. Click within the ROI selection and drag and drop it to an area of background.
8. Navigate to **Analyze > Measure** to measure the area and mean gray value of the background.
9. Record measurements of your ROI and background. The background-subtracted mean gray value (MGV) can be calculated as MGV_ROI_ - MGV_background_.

1.	Martinez MAQ, Matus DQ. 2020. Auxin-mediated Protein Degradation in Caenorhabditis elegans. Bio Protoc 10.

2.	Stiernagle T. Maintenance of C. elegans doi:10.1895/wormbook.1.101.1. WormBook.
