## Supplemental File 5 for "A laboratory module that explores RNA interference and codon optimization through fluorescence microscopy using *Caenorhabditis elegans*"

**S5. A Laboratory Module-Student GFP RNAi Worksheet**

This worksheet will help you work through the GFP RNAi experiment.

**Questions to think about to form your hypothesis:**

1. Do you think the GFP RNAi clone will target one or both GFP variants (GFP_NCO_ or GFP_CO_ variants) (hint, does RNAi specificity function at the **nucleotide** or **amino acid** level)?
2. How similar are the two GFP variants at the amino acid level? At the nucleotide level?

**Data you need from us:**

Nucleotide sequences:

>GFP_NCO_(DQM594)

ATGAGTAAAGGAGAAGAACTTTTCACTGGAGTTGTCCCAATTCTTGTTGAATTAGATGGTGATGTTAATGGGCACAAATTTTCTGTCAGTGGAGAGGGTGAAGGTGATGCAACATACGGAAAACTTACCCTTAAATTTATTTGCACTACTGGAAAACTACCTGTTCCATGGCCAACACTTGTCACTACTTTCTgTTATGGTGTTCAATGCTTcTCgAGATACCCAGATCATATGAAACgGCATGACTTTTTCAAGAGTGCCATGCCCGAAGGTTATGTACAGGAAAGAACTATATTTTTCAAAGATGACGGGAACTACAAGACACGTGCTGAAGTCAAGTTTGAAGGTGATACCCTTGTTAATAGAATCGAGTTAAAAGGTATTGATTTTAAAGAAGATGGAAACATTCTTGGACACAAATTGGAATACAACTATAACTCACACAATGTATACATCATGGCAGACAAACAAAAGAATGGAATCAAAGTTAACTTCAAAATTAGACACAACATTGAAGATGGAAGCGTTCAACTAGCAGACCATTATCAACAAAATACTCCAATTGGCGATGGCCCTGTCCTTTTACCAGACAACCATTACCTGTCCACACAATCTGCCCTTTCGAAAGATCCCAACGAAAAGAGAGACCACATGGTCCTTCTTGAGTTTGTAACAGCTGCTGGGATTACACATGGCATGGATGAACTATACAAA

> GFP_NCO__amino_acids

MSKGEELFTGVVPILVELDGDVNGHKFSVSGEGEGDATYGKLTLKFICTTGKLPVPWPTLVTTFCYGVQCFSRYPDHMKRHDFFKSAMPEGYVQERTIFFKDDGNYKTRAEVKFEGDTLVNRIELKGIDFKEDGNILGHKLEYNYNSHNVYIMADKQKNGIKVNFKIRHNIEDGSVQLADHYQQNTPIGDGPVLLPDNHYLSTQSALSKDPNEKRDHMVLLEFVTAAGITHGMDELYK

>GFP_CO_ (DQM583)

ATGAGTAAAGGAGAAGAATTGTTCACTGGAGTTGTCCCAATCCTCGTCGAGCTCGACGGAGACGTCAACGGACACAAGTTCTCCGTCTCCGGAGAGGGAGAGGGAGACGCCACCTACGGAAAGCTCACCCTCAAGTTCATCTGCACCACCGGAAAGCTCCCAGTCCCATGGCCAACCCTCGTCACCACCTTCTGCTACGGAGTCCAATGCTTCTCCCGTTACCCAGACCACATGAAGCGTCACGACTTCTTCAAGTCCGCCATGCCAGAGGGATACGTCCAAGAGCGTACCATCTTCTTCAAGGACGACGGAAACTACAAGACCCGTGCCGAGGTCAAGTTCGAGGGAGACACCCTCGTCAACCGTATCGAGCTCAAGGGAATCGACTTCAAGGAGGACGGAAACATCCTCGGACACAAGCTCGAGTACAACTACAACTCCCACAACGTCTACATCATGGCCGACAAGCAAAAGAACGGAATCAAGGTCAACTTCAAGATCCGTCACAACATCGAGGACGGATCCGTCCAACTCGCCGACCACTACCAACAAAACACCCCAATCGGAGACGGACCAGTCCTCCTCCCAGACAACCACTACCTCTCCACCCAATCCGCCCTCTCCAAGGACCCAAACGAGAAGCGTGACCACATGGTCCTCCTCGAGTTCGTCACCGCCGCCGGAATCACCCACGGAATGGACGAGCTCTACAAG

>GFP_CO__amino_acids

MSKGEELFTGVVPILVELDGDVNGHKFSVSGEGEGDATYGKLTLKFICTTGKLPVPWPTLVTTFCYGVQCFSRYPDHMKRHDFFKSAMPEGYVQERTIFFKDDGNYKTRAEVKFEGDTLVNRIELKGIDFKEDGNILGHKLEYNYNSHNVYIMADKQKNGIKVNFKIRHNIEDGSVQLADHYQQNTPIGDGPVLLPDNHYLSTQSALSKDPNEKRDHMVLLEFVTAAGITHGMDELYK

>GFP_NCO_ RNAi_sequence

ATGAAAGGGAGAAGAACTTTTCCTGGAGTTGTCCCAATTTTGTTGAATTAGATGGTGATGTTAATGGGCACAAATTTTCTGTCAGTGGAGAGGGTGAAGGTGATGCAACATACGGAAAACTTACCCTTAAATTTATTTGCACTACTGGAAAACTACCTGTTCCATGGCCAACACTTGTCACTACTTTCTCTTATGGTGTTCAATGCTTTTCAAGATACCCAGATCATATGAAACGGCATGACTTTTTCAAGAGTGCCATGCCCGAAGGTTATGTACAGGAAAGAACTATATTTTTCAAAGATGACGGGAACTACAAGACACGTGCTGAAGTCAAGTTTGAAGGTGATACCCTTGTTAATAGAATCGAGTTAAAAGGTATTGATTTTAAAGAAGATGGAAACATTCTTGGACACAAATTGGAATACAACTATAACTCACACAATGTATACATCATGGCAGACAAACAAAAGAATGGAATCAAAGTTAACTTCAAAATTAGACACAACATTGAAGATGGAAGCGTTCAACTAGCAGACCATTATCAACAAAATACTCCAATTGGCGATGGCCCTGTCCTTTTACCAGACAACCATTACCTGTCCACACAATCTGCCCTTTCGAAAAGATCCCAAACGAAAAGAGAGACCACATGGGTCCTTTCTTGAGTTTGTAACAGCTGGCTGGGGATTACACATGGCATGGATGAACTATACAA

**Instructions:**

*Steps 1-4 can be done in or out of class

1. Generate nucleotide and amino acid alignments - pair-wise or multi-alignments. You can make pair-wise alignments here (select DNA for nucleotide alignments and PROTEIN for amino acid alignments):
   - [<https://www.ebi.ac.uk/Tools/psa/emboss_needle/>](https://www.ebi.ac.uk/Tools/psa/emboss_needle/)
1. **Save these alignments as a PDF or as text files and format them to include in your lab report as Figure 3.**
1. What is the % match and number of mis-matched nucleotides between the two GFP constructs? ___________
2. What is the % match and number of mis-matched amino acids between the two GFP constructs? ___________
3. What is the % match and number of mis-matched nucleotides between the GFP RNAi construct and GFP NCO (DQM594) _____and between GFP RNAi and GFP CO (DQM583) _____?
