## Supplemental File 6 for "A laboratory module that explores RNA interference and codon optimization through fluorescence microscopy using *Caenorhabditis elegans*"

**S6. A Laboratory Module-Student instructions for GFP RNAi Module**

**The GFP RNAi experiment:**

**The purpose of this exercise is multi-fold**. By the end of this experiment, you will have:

- learned how perform feeding RNAi experiments in *C. elegans*
- learned about the **specificity of RNAi**
- gained familiarity with quantitative fluorescence microscopy
- generated publication quality figures and figure legends
- demonstrated ethical and responsible use of acquired shared data

**Experiment:**

**How to visualize depletion of GFP by RNAi treatment and quantify fluorescence intensity using Fiji/ImageJ.**

In this feeding RNAi experiment you will treat two transgenic strains of *C. elegans* (**DQM594 and DQM583**) that broadly express a GFP-tagged transgene fused to the histone, H2B, with dsRNA that targets GFP. Your instructors will provide you with a synchronous population of developmentally arrested L1 stage animals, which you will aliquot onto two plates: an experimental treatment plate (GFP RNAi plate) and a negative control plate (T444T/empty vector), which contains an “empty vector” version of the plasmid that generates dsRNA in the same bacterial strain as the experimental treatment (HT115 *E. coli*).

Your instructor(s) have plated L1 stage animals at the appropriate time so that on Day 1 you can score the results of the experiment using 48-hour old animals, using epifluorescence microscopes.

You and your partner(s) will work with **either DQM594 (GFP­_NCO_) or DQM583 (GFP_CO_) for one lab session and then work with the other strain during the next lab session.** The two strains contain **two different versions of GFP**: eGFP (original enhanced GFP or GFP_NCO_) **or** codon optimized GFP (GFP_CO_). For more information on the strains see page 7.

**Session 1:**

**Quantitative fluorescence imaging of GFP and documentation of the GFP RNAi experiment**

- Approximately 15 minutes before it is your turn to use one of the compound microscopes, make two slides with at least 5-10 worms each of T444T & GFP(RNAi) treated worms. Your goal is to image 10 worms per strain per condition.
- Collect transmitted light images and GFP images of the control (T444T) worms **first** (your instructor(s)/TA(s) will have determined the ideal exposure setting prior to the start of your imaging session)

**Question for students: Why is it important to use the same exposure setting between control and experimental treatments?**

- Collect transmitted light and GFP images of the GFP(RNAi) treated worms **second**

**Session 2:**

Continue the quantitative fluorescence imaging of GFP and documentation of the GFP RNAi experiment as you did in Lab 2 using whichever strain you did not yet image.

**Data Analysis:**

Once you have collected 10 images (10 animals) of each strain for each condition, you can analyze your data. To do this make sure you have installed Fiji/Image on the computer you are using. Then follow the directions provided in the Tutorial Videos 1-5.

Once you have the Mean Fluorescence Intensity for all of your images, you can plot your data and perform the appropriate statistical test (i.e. student’s *t-test)*.

**Instructions to graph your data:**

Graph your data, we recommend a dot plot using R (if you know how to use R), Graph Pad Prism (if available) or the online web tool “Plots of Data”. Below are instructions for Plots of Data.

**Using Plots of Data [1] to plot your results:**

(<https://huygens.science.uva.nl/PlotsOfData/>)

1.
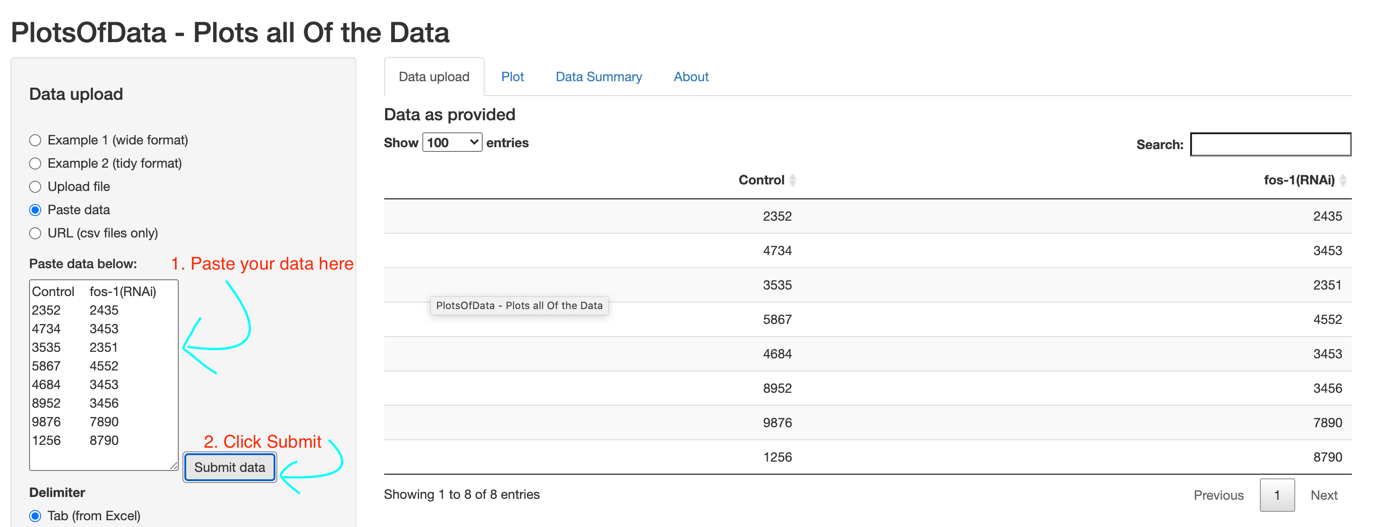
Navigate to the website and select “Paste data” and Paste your data from excel/Fiji and click “submit”
2.
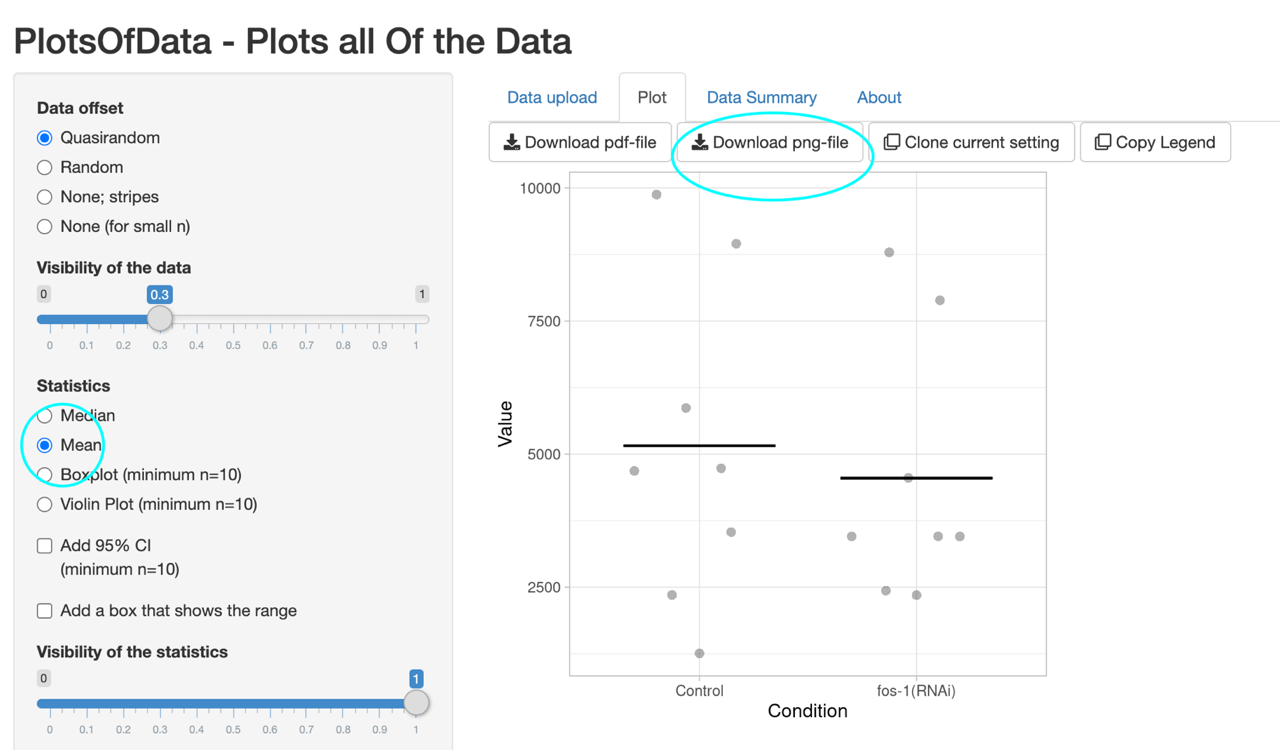
Click the “Plot” tab (next to “Data Upload” and select “Mean” – you can customize the plot, label axes, etc. and then Export the plot as a .png file for inserting into your lab report assignment.
3. Annotate significance (*) or lack of significance (n.s) from the result of your *t-test* on the plot (your plot will not look like this, this is random data):

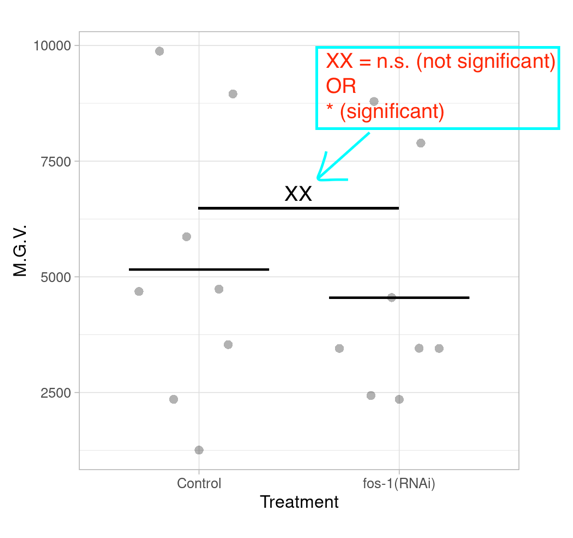

Listen to the information from your instructors about statistical testing for the GFP RNAi experiment. For lots of great background reading on doing stats in a *C. elegans* lab, go to the [wormbook.org](http://wormbook.org) chapter on stats from Dr. David Fey (University of Wyoming):

<http://www.wormbook.org/chapters/www_statisticalanalysis/statisticalanalysis.html>

Compile images and plot of data for your lab report. See Tutorial Video 5 (Supporting file S7. A Laboratory Module-Student Transcripts for Tutorial Videos 1-5) for how to generate images complete with scale bars for your figure.

**For your lab report**:

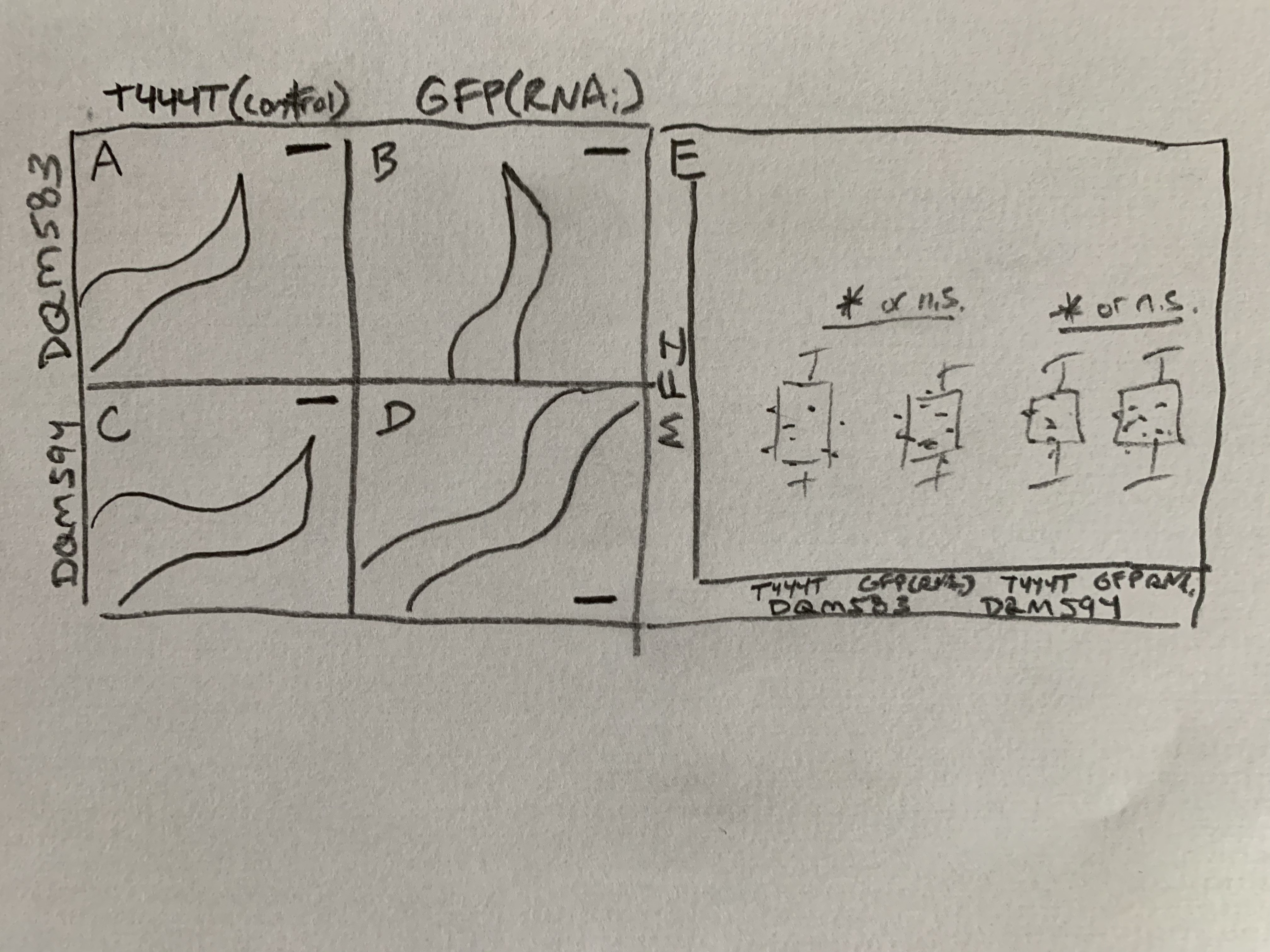

Figure 2. **Title**. (A)… (B)… (C)… (D)… (E) Plot… *, P=xxx, Student’s t-test or n.s., P=xxx, Student’s t-test. (see S2. A Laboratory Module-Grading rubric and example lab report for more information).

Table 1. **Title**.

| **Image #** | **DQM583** | | **DQM594** | |
| --- | --- | --- | --- | --- |
|  | **T444T** | **GFP(RNAi)** | **T444T** | **GFP(RNAi)** |
| 1 | xxx | xxx | xxx | xxx |
| 2 | xxx | xxx | xxx | xxx |
| 3 | xxx | xxx | xxx | xxx |
| 4 | xxx | xxx | xxx | xxx |
| 5 | xxx | xxx | xxx | xxx |
| 6 | xxx | xxx | xxx | xxx |
| 7 | xxx | xxx | xxx | xxx |
| 8 | xxx | xxx | xxx | xxx |
| 9 | xxx | xxx | xxx | xxx |
| 10 | xxx | xxx | xxx | xxx |
| **Mean** | xxx | Xxx | xxx | xxx |

Grey shaded boxes indicate images selected based on mean fluorescence intensity for Figure 2.

**Additional Information on Worm Strains**

**A. Strain list**

Strain names are listed in brackets. Promoter fusions are designated by a “ > ” symbol, whereas linkage of open reading frames is denoted by a “ :: ” symbol. Promoters and mutant alleles are listed in italics; encoded protein products are listed in Roman. Semicolons separate distinct transgenes or alleles within the same strain. Brief descriptions follow in parentheses; see the subsequent section for more detail.

[DQM594]	***bmd170 [eft-3>H2B::GFP_NCO_ Chr I]***

[DQM583]	***bmd141*** [***eft-3>H2B::GFP_CO_ (codon-optimized) Chr I]***

**B. Descriptions of transgenes**

***bmd170***	*H2B::GFP_NCO_* fusion protein under the control of a strong ubiquitous	promoter (*eft-3*) inserted by CRISPR/Cas9-genome engineering at a	neutral site on Chromosome I. This GFP transgene is silenced in the	germline.

***bmd141***	*codon optimized H2B::GFP* fusion protein under the control of a strong	ubiquitous promoter (eft-3) inserted by CRISPR/Cas9-genome	engineering at a neutral site on Chromosome I.
