## Supplemental File 7 for "A laboratory module that explores RNA interference and codon optimization through fluorescence microscopy using *Caenorhabditis elegans*"

**S7. A Laboratory Module-Transcripts for Tutorial Videos 1-5**

**Tutorial Videos are available** [**here**](https://www.dropbox.com/sh/bfh5dobn9v8gprf/AAACJO0nsfKb8oaFr-kxv25Ea?dl=0)**.**

In this instructional video series Maryam and I will be going over the steps on how to quantify data successfully using FIJI/ImageJ as well as produce a publication quality figure.

**Tutorial Video 1: Opening Images in Fiji/Image J – Adjusting Brightness and Contrast**

Frances:

Welcome to Part 1 of the 5-part instructional video series. In this video I will be taking you through how to upload your image on ImageJ as well as how to adjust the brightness and contrast of your image and set your parameters. You should see to the left a series of shortcuts that may be beneficial when using ImageJ that Maryam generated. At any point feel free to write these shortcuts down or screen capture them at your own discretion. Note that mac users should replace the control button on the list with their command button. If ImageJ takes up your entire screen, click on the “restore down” button to minimize it. To start you are going to want to take the image you want to analyze and drag it to the ImageJ toolbar. You should see a pop-up window that has your image. Ensure that the images you’re working on are not your original data because ImageJ can only undo once. To avoid permanently altering your data, right click on your image and click on duplicate making sure to click duplicate stack. You can then rename your duplicated image whatever you’d like for ease of presenting. We will leave the title how it is. You can click the bottom bar to switch between your DIC channel and your fluorescence channel. Moving on to analyzing your data you are going to want to click on the toolbar and select analyze at the top of your screen then set measurements. Here you may see a few boxes already pre-selected. You are going to want to unselect each of these and select ‘mean gray value’. Next you should click ‘Image’ Adjust Brightness and Contrast. From there you should see a smaller pop up window that has four slide bars. You can manually adjust brightness and contrast by sliding the slide bars provided or you can click the auto button. Be careful when adjusting the brightness and contrast not to click apply as this will permanently alter the brightness and contrast of your image. In the next video we will discuss how to outline your worm and measure the mean gray value.

**Tutorial Video 2: Measuring Mean Fluorescence Intensity in Fiji/Image J: Single Z data**

Frances:

Welcome to Part 2 of the 5-part instructional video series. In this video I will be discussing two techniques for how to outline your worm as well as how to account for background microscope noise and measure the mean gray value. The first tool you can use to outline your worm is the freehand tool. This technique is recommended for people who have a drawing tablet or other drawing tools. Here you are going to want to trace your worm as precisely as possible. Since Maryam is using a mouse you can see that the outline is not precise. After drawing this outline you're going to want to switch to the fluorescence channel. From there you are going to click analyze and measure or alternatively you can click the letter M on your keyboard. You should see your mean gray value pop up. Alternatively, if you don’t have a drawing tablet or other precision tools and you have a mouse or trackpad you can select the polygon selections tool from the toolbar. This tool allows for more precision as you can see the outline is much closer to the actual worm outline. Then again you can click analyze and measure. Both numbers should be similar however you can use whatever tool gets you the more precise results. Now we need to take into account any background microscope noise that may be present. In order to do this, we need to determine the darkest part of the image. To do that, go to image → adjust → threshold (alternatively, you can use the shortcut ctrl+shift+T). Ensure that you have the dark background selected. Adjust the sliders and the last part of the image to turn red will be the area where you take your background measurement from - ensure that the area is not out of the field of view. If ‘dark background’ is not selected, the last region of your image to turn red will be the area where you take your background value from. Click the rectangle tool and draw a rectangle in the region that is what you determined to be the darkest area. Next click on analyze tools and then ROI manager. A small pop up box will appear click add. This will allow you to keep the same box area between each image to allow for consistent data measurements. You reset to remove the threshold. Do not click on apply! Take your background measurement by clicking M. Continue acquiring the measurements for the remaining worms. It is important to keep track of the image that the values are coming from when we need to determine our representative images. To save your values, right click and select all of the values and then right click again to copy them and paste them onto a spreadsheet. We will discuss this step in further details in Instructional video part 4. In the next video Maryam will go over how to quantify single cell data.

**Tutorial Video 3: Measuring Mean Fluorescence Intensity in Fiji/Image J: Confocal Z-stack data**

Maryam:

Welcome to the 3rd part of the instructional video series. Here, I’ll be showing you how to measure the mean gray value of single cell data and take the background noise measurements for that cell. We’ll be using the free-hand tool for this part as it will be faster and more precise in this case. However, feel free to use the polygon tool discussed in the 2nd instructional video series if you prefer. Select the freehand selection tool from the toolbar. You will notice that your image consists of 2 channels, a differential interference contrast channel (DIC) and a fluorescence channel and it has a separate slider below the channel’s slider. This slider denotes each z-stack through the worm’s body. It’s best to zoom into the nucleus you want to quantify for better clarity and ease of outlining, you can use the + sign on your keyboard to zoom in. Outline the nucleus and go to analyze → measure or alternatively, you can click on the letter M on your keyboard as the shortcut. After you’re obtained your value, you can grey out the nucleus that you have measured, in order to keep track of what you’ve already done. You can also use the pencil tool on the toolbar to number the nucleus. Refer to our 2nd instructional video for more information on how to determine the region to subtract your background. I have already predetermined the darkest part of my background to be here. Drag the outline of your nucleus over to the darkest part of your image and take your background measurement by clicking M. Continue acquiring the measurements for the remaining worms. It is important to keep track of the image that the values are coming from when we need to determine our representative images. To save your values, right click and select all of the values and then right click again to copy them and paste them onto a spreadsheet. We will discuss this step in further details in the next Instructional video.

**Tutorial Video 4: Compiling Data**

Maryam:

Welcome to the 4th part of the instructional video series. Here we will go over ways to transfer our data from the results window into a spreadsheet and determine how to pick your representative images. Start by clicking on results from your results window and select options. Uncheck the boxes for copy column headers and copy row numbers. And then click on ok. Next, right click on the results window and choose select all. Alternatively, you can use the shortcut control+A for windows or command+A if you’re using mac to select all the values. Right click again and choose copy and once again you can use the keyboard shortcut control/command+C to copy the measurements. Paste the values in your spreadsheet by right clicking on a cell and choosing paste or using the shortcut control/command+V. All that is left to do is to obtain the mean fluorescence intensity of each of value which is done by subtracting the mean gray value by the background value. Go to an empty cell and click on the = sign to start your formula and from there choose the cell with your mean gray value - the cell with your background value. When you have finished doing this for all your values, calculate the average of your mean fluorescent intensities. The formula is = average, open brackets, select the range of cells that you want the average of and click enter. The image whose mean fluorescent intensity is closest to the mean will be your representative image.

The next, and final, video of this series will show you how to add scale bars to your representative images, assign them the same brightness and contract values for visual comparison and transform your worm to follow the *C. elegans* axis convention.

**Tutorial Video 5: Formatting images for figure generation**

Frances:

Welcome to the final instructional video of this series. In this video I will show you how to add scale bars to your representative images, assign them the same brightness and contrast values for visual comparison and transform your worm to follow the *C. elegans* axis convention. We will begin with assigning our images the same brightness and contrast values to ensure that all of our images are comparable. Therefore, we must set the range of our values based on our brightest strain and condition. This will be the strain and condition with the highest mean fluorescent intensity average. In this case it is the codon optimized strain coupled with the control RNAi. By using Auto, almost 10% of our data gets saturated, which means that we cannot differentiate between individual details. To avoid over saturation, slide the maximum slider until you are able to see each individual nucleus with the nucleolus. When Maryam zooms in on these nuclei, you can see that saturation covers the details, whereas reducing the maximum range of brightness and contrast enables us to view the details. Click on set and take note of the min and max values. You may choose to propagate these values to all the images if you have them open, however, this will also propagate to your DIC channel. Click on the next image, go to set and type in the values and click okay. My values were 1 and 57. Do this for each strain at each condition. Next, we will rotate our worm to be in the correct direction based on *C. elegans*’ convention. This means that the worm should be positioned such that it’s head or anterior is to the left, it’s tail or posterior is to the right, its dorsal side is up and ventral side is down. In this case, the crescent here, is the ventral side. You can see the pharynx here, that is the anterior side. Click on Image → transform → rotate. Check the preview box and adjust the angle until you get the correct axis. Click on ok. Sometimes you might have to flip your image horizontally or vertically. These options are also available under transform. In order to work on the scale bars you must know the microns per pixel value. Based on the microscope used and the magnification that the image was acquired at, your instructor will provide you with your specific microns per pixel value for your images. To enter this value, go to image → properties and enter the “microns” for unit length and the value in pixel width. Height will automatically and voxel depth will be 0 for your compound scope images, or step-size in microns for your confocal images. Check global and click ok. Consider using region of interest, which is the ROI manager, to ensure that your micrographs will be the same size. Once you have selected the area for your micrograph, right click to duplicate the image. To insert the scale bar, go to analyze → tools → scale bar and change the settings to your liking. Click on okay when you are done. Select the image that you would like to copy and click on edit → copy to system. Paste the image into the document software of your choice. Ensure that you label what each micrograph’s strain and condition are. Feel free to contact us on our emails listed to the left.

Good luck!

**Shortcuts:**

Analyze --> Set Measurement --> check "Mean Gray Value" (uncheck everything else) --> OK

+: Zoom in

-: Zoom out

Mouse scroll bar: Scroll through channels

Hand tool = Space bar: grip tool - use touchpad/mouse to drag through image

Analyze --> Measure = M key on keyboard

Image --> Adjust --> Brightness & Contrast= Ctrl+Shift+C

Image --> Adjust --> Threshold = Ctrl+Shift+T

Analyze --> Tools --> ROI Manager = T key on keyboard

Edit --> Fill (fills all images) = Ctrl+F or Backspace: Remove part of image (fills only 1 slice)

Results Window --> Results --> Options --> uncheck "Copy row numbers" and "Copy column headers" --> OK

Ctrl+C: Copy measurement results

**Spreadsheet:**

Ctrl+V: Paste data

Ctrl+C: Copy data
