## Supplemental File 13 for "A laboratory module that explores RNA interference and codon optimization through fluorescence microscopy using *Caenorhabditis elegans*"

**S13. A Laboratory Module-Jigsaw Active Learning Activity & Post-Module Assessment (Optional)**

**Overview:**

In this activity based on the jigsaw method teaching strategy, students will randomly be put into groups of 3 students and each assigned a number, 1 through 3. In Phase I, students will go to a learning station based on the number they are assigned to become an “expert” in one of the following areas: (1) transcription and translation, (2) codon bias and optimization, and (3) sequence alignment. We estimate that 30-60 minutes should be sufficient time for each group to become “experts” on their respective activities. Questions associated with the learning levels of Bloom’s taxonomy are provided at the end of each activity to assess student mastery of their respective topics (this completes Phase I). After the completion of Phase I, new groups are formed, with each group consisting of one expert from each original group. Each expert teaches their expertise to their group members. A post-activity assessment is provided for students to work on in groups or independently (this completes Phase II). During each phase, instructors should join each group to hear conversations and provide any feedback they see fit.

**Phase I**

1. **Transcription and translation**

*Learning goal:*

Students will learn about the central dogma of molecular biology (the flow of information from DNA to mRNA to protein.

*Learning objective:*

Students will be able to transcribe and translate any DNA sequence of interest into a protein product.

*Instructions:*

1. In a web browser, navigate to the University of Utah’s Genetic Science Learning Center website (<https://learn.genetics.utah.edu/content/basics/txtl/>).
2. Complete the virtual “Transcribe and Translate a Gene” interactive activity available. After selecting an example gene, students will visualize the process of transcription and translation, and test their knowledge by selecting correct base pair and codon matches (screenshots of virtual activities below).

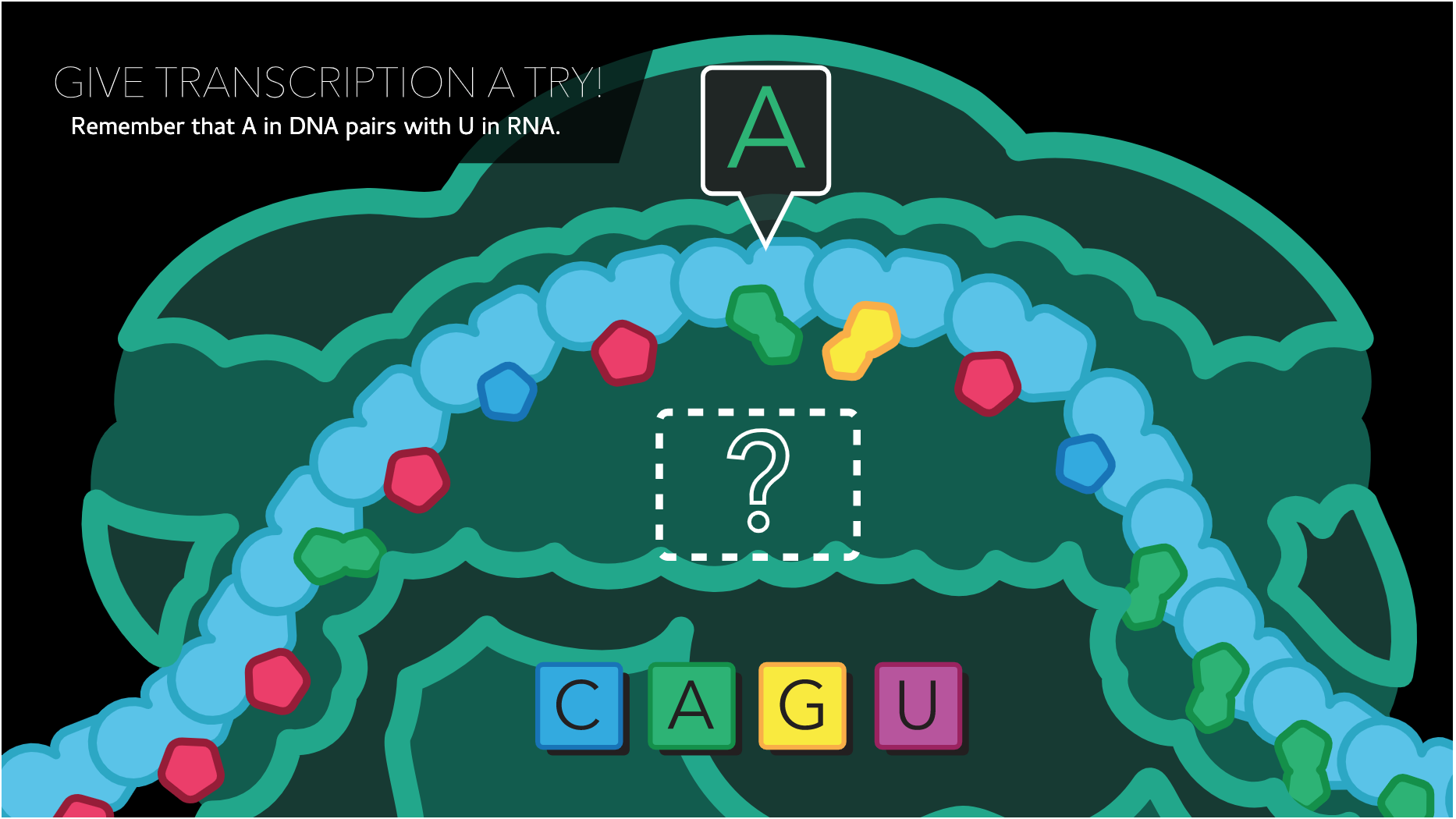

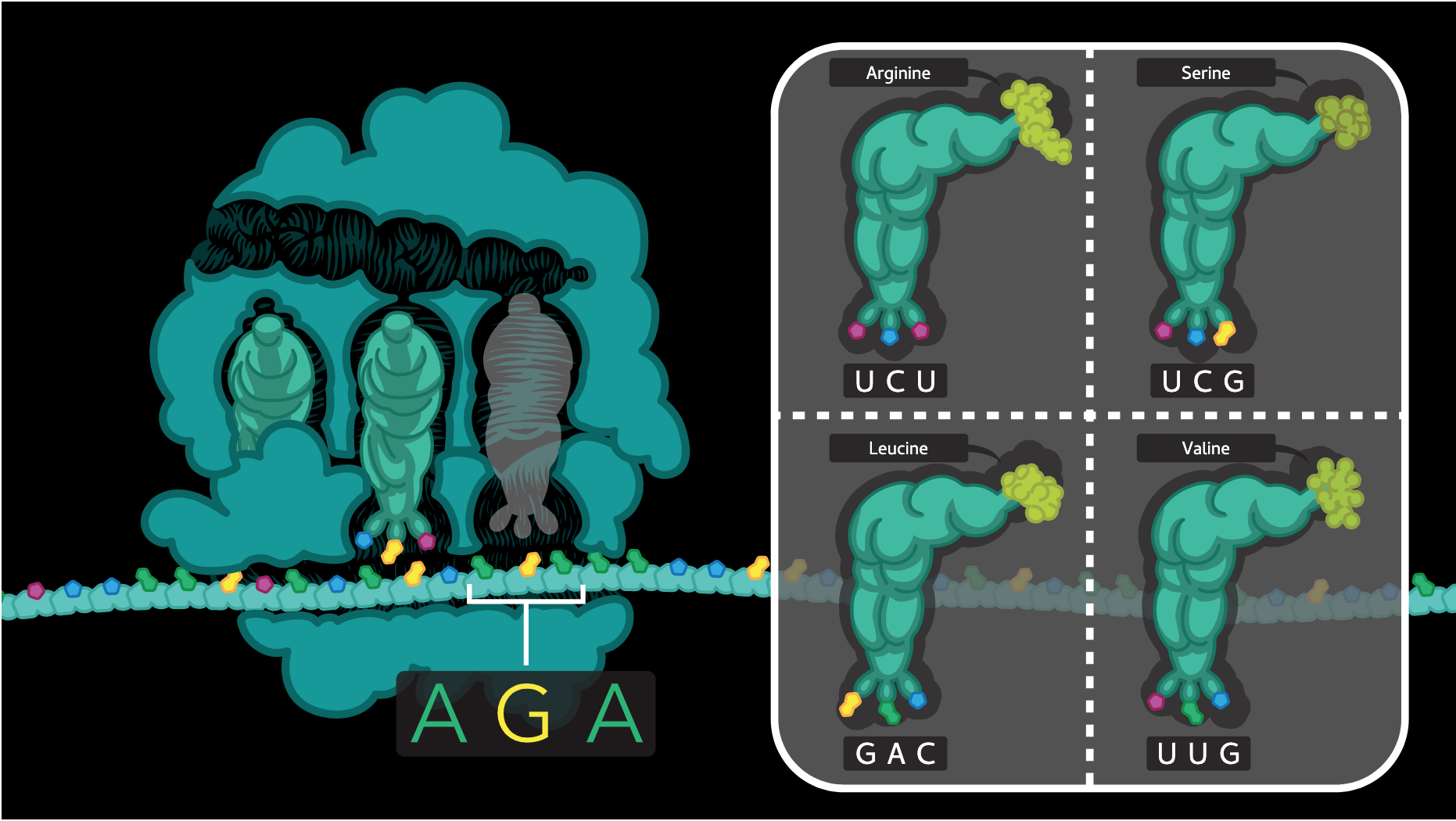

Questions for students *(Bloom’s taxonomy level: Remembering)*:

**Please note that answers are for instructors only*

1. Define what a codon is and describe its importance in the translation process?

*Answer:* A codon consists of 3 nucleotide bases in mRNA that codes for a specific amino acid. Codons must be translated one after another to build a functional protein.

2. What are transfer RNAs (tRNAs) and how are they important in the translation process?

*Answer:* tRNAs are key components of the translational process that link amino acids to appropriate codons in mRNA. Each tRNA molecule contains a set of 3 nucleotides called an anticodon along with a corresponding amino acid. These anticodons bind to codons on mRNA inside of the ribosome. Therefore the pairing of anticodons on tRNA molecules to codons on mRNA molecules delivers amino acids to a growing polypeptide chain.

3. In your own words, briefly describe the **general steps** needed to convert a sequence of DNA into a protein?

*Answer:* The formation of a protein product from DNA occurs through two major steps. The first step is transcription, which involves the conversion of DNA to mRNA. This process is accomplished by RNA polymerase II, which uses DNA as a template to catalyze the formation of pre-mRNA through complementary base-pairing. This pre-mRNA is then processed into mature mRNA. The second step is translation. Here, mRNA serves as a template for the formation of a polypeptide product. In this process, ribosomes match tRNA anticodon sequences and accompanying amino acids to complementary codon sequences of mRNA. Translation begins with a methionine and ends with one of three stop codons (UAA, UAG, or UGA).

2. **Codon bias and optimization**

*Learning goal:*

Students will learn that many amino acids are specified by multiple RNA codons (Table 1), and learn about codon bias and the codons most commonly used in the *C. elegans* genome (Table 2; data from Sharp and Bradnam,1997^1^).

*Learning objective:*

Students will be able to use the *C. elegans* Codon Adapter tool to optimize any DNA sequence of interest for optimal expression in *C. elegans.*

*Instructions:*

Students will use the online *C. elegans* Codon Adapter tool^2^ to optimize a sequence for expression in *C. elegans* by entering the coding sequence into the box.

1. In a web browser, navigate to:

<https://blog.addgene.org/to-codon-optimize-or-not-that-is-the-question>. Read this blog to gain a thorough understanding of codon usage bias.

2. After reading the blog above, navigate to: <https://worm.mpi-cbg.de/codons/cgi-bin/optimize.py>.
3. Copy the sequence provided below, which is the coding sequence for mCerulean, a fluorescent protein derived from the jellyfish *Aequorea victoria*:

ATGGTGAGCAAGGGCGAGGAGCTGTTCACCGGGGTGGTGCCCATCCTGGTCGAGCTGGACGGCGACGTAAACGGCCACAAGTTCAGCGTGTCCGGCGAGGGCGAGGGCGATGCCACCTACGGCAAGCTGACCCTGAAGTTCATCTGCACCACCGGCAAGCTGCCCGTGCCCTGGCCCACCCTCGTGACCACCCTGACCTGGGGCGTGCAGTGCTTCGCCCGCTACCCCGACCACATGAAGCAGCACGACTTCTTCAAGTCCGCCATGCCCGAAGGCTACGTCCAGGAGCGCACCATCTTCTTCAAGGACGACGGCAACTACAAGACCCGCGCCGAGGTGAAGTTCGAGGGCGACACCCTGGTGAACCGCATCGAGCTGAAGGGCATCGACTTCAAGGAGGACGGCAACATCCTGGGGCACAAGCTGGAGTACAACGCCATCAGCGACAACGTCTATATCACCGCCGACAAGCAGAAGAACGGCATCAAGGCCAACTTCAAGATCCGCCACAACATCGAGGACGGCAGCGTGCAGCTCGCCGACCACTACCAGCAGAACACCCCCATCGGCGACGGCCCCGTGCTGCTGCCCGACAACCACTACCTGAGCACCCAGTCCAAGCTGAGCAAAGACCCCAACGAGAAGCGCGATCACATGGTCCTGCTGGAGTTCGTGACCGCCGCCGGGATCACTCTCGGCATGGACGAGCTGTACAAG

4. Paste the sequence into the white space labeled “sequence”. Under name, enter mCerulean.
5. Under “Options”, use the default settings where “introns” (the number of synthetic introns added to the codon optimized sequence) is set to 3 and the “Codon Adaptation Index” is set to 1.0. The inclusion of synthetic introns has been shown to enhance protein expression in *C. elegans*. The codon adaption index is an effective measure of synonymous codon usage bias(1).
6. Once complete, click the button labeled “Optimize”. The “optimized sequence” under “Results” is a codon optimized sequence of mCerulean, which contains artificial introns (lowercase letters). New codons in the optimized sequence are highlighted in green, and the bar graphs at the bottom of the page compare the weight or relative adaptiveness of the codons.

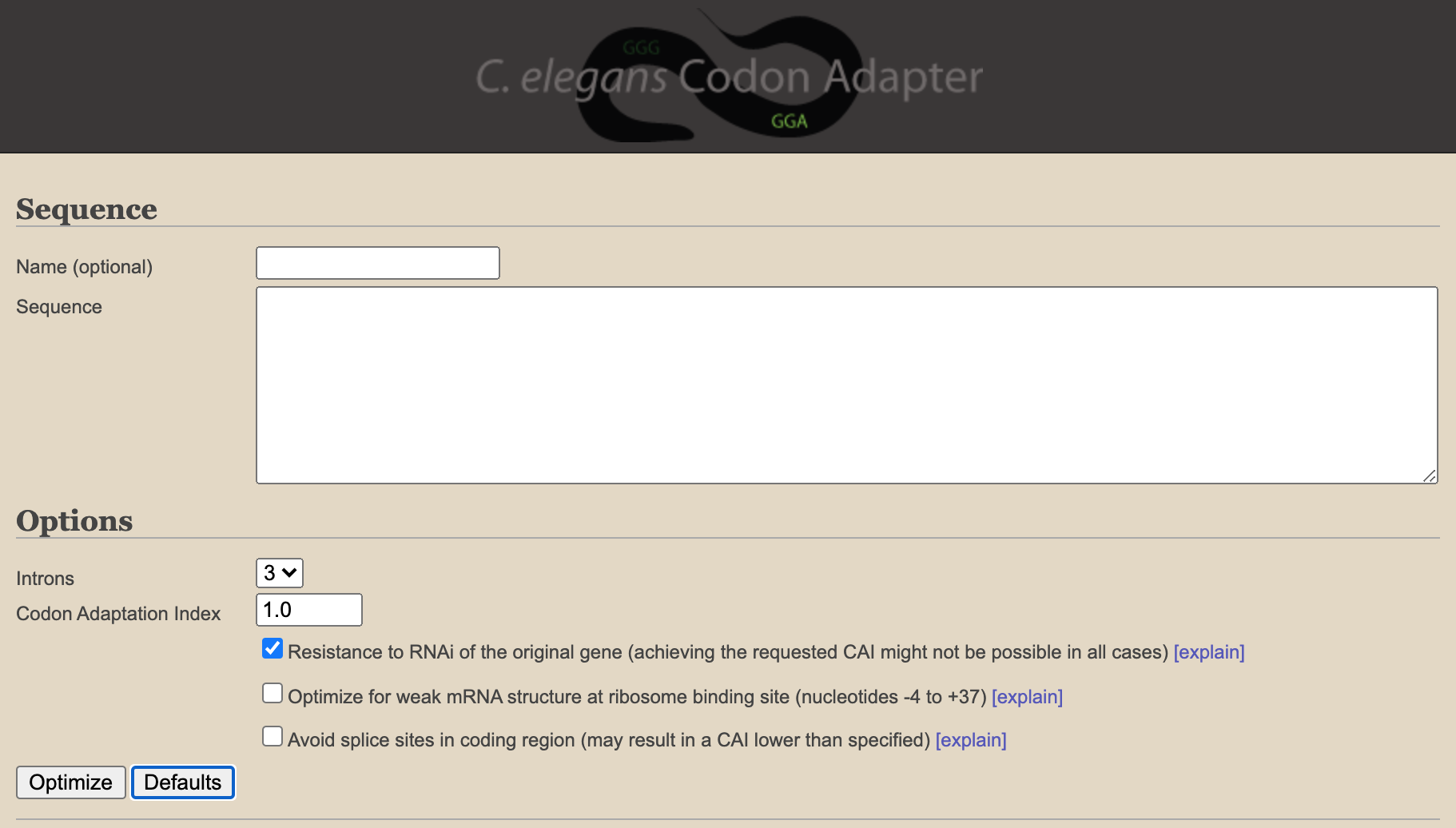

*Questions for students (Bloom’s taxonomy level: Understanding* and *applying)*:

1. Based on your readings, explain one possible consequence that may arise when introducing a non-codon optimized gene from one organism into another (i.e. from *D. melanogaster* to *C. elegans*)?

*Answer:* One possible consequence of introducing a non-codon optimized gene from one organism to another is that the gene may not be robustly expressed. One reason for this could be that the sequence of codons that make up the gene in the original organism are not common in the new organism. As a consequence, there may be a limitation in the availability of tRNAs for those unique codons, affecting the rate of translation.

2. When would it be important to ensure that the codon optimized version of a gene is resistant to RNAi that targets the original non-codon optimized version of the gene?

*Answer:* One possible situation where this would be important is when designing and testing the functionality of a translational fluorescence-tagged reporter for a gene of interest. Here, a scientist may want to determine if their fusion reporter rescues phenotypes that arise when the endogenous gene of interest is depleted, thereby showing their fusion reporter is “functional”. In this scenario, scientists would want to ensure that the codon optimized version of the gene of interest that is tagged to a fluorescent reporter is not affected by RNAi that targets the endogenous version of the gene of interest.

3. Why are coding DNA sequences, also known as CDS, which are sequences of DNA lacking introns, used during this analysis?

*Answer:* A coding sequence of DNA (CDS) is a region of DNA (exons) that corresponds to the sequence of a mature mRNA molecule, which in turn corresponds to the amino acid sequence of a protein. Given that codon optimization is important for efficient translation of mature mRNA into protein, and given that coding sequences of DNA correspond to mature mRNA, coding sequences of DNA are used in the analysis.

3. **Sequence alignment**

*Learning goal:*
Students will learn how to compare DNA sequences and understand the importance of reading frame.

*Learning objective:*

Students will be able to use the DNA sequence alignment tool from the European Bioinformatics Institute to compare the percent similarity and mismatches between two related nucleotide sequences.

*Instructions:*

1. In a web browser, navigate to:

<https://bitesizebio.com/9445/the-beginners-guide-to-dna-sequence-alignment/>. Read this webpage to gain a thorough understanding of the basics of nucleotide sequence alignments.

2. After reading the above webpage, in a new web browser page, navigate to the European Bioinformatics Institute webpage: <https://www.ebi.ac.uk/Tools/psa/emboss_needle/>.
3. Under “Step 1-Enter your nucleotide sequences”, use the drop-down menu to select for DNA. Using the DNA sequences provided below from two red fluorescent proteins, copy and paste both nucleotide sequences that are arranged in FASTA format.

>TagRFP-T

ATGGTGTCTAAGGGCGAAGAGCTGATTAAGGAGAACATGCACATGAAGCTGTACATGGAGGGCACCGTGAACAACCACCACTTCAAGTGCACATCCGAGGGCGAAGGCAAGCCCTACGAGGGCACCCAGACCATGAGAATCAAGGTGGTCGAGGGCGGCCCTCTCCCCTTCGCCTTCGACATCCTGGCTACCAGCTTCATGTACGGCAGCAGAACCTTCATCAACCACACCCAGGGCATCCCCGACTTCTTTAAGCAGTCCTTCCCTGAGGGCTTCACATGGGAGAGAGTCACCACATACGAAGACGGGGGCGTGCTGACCGCTACCCAGGACACCAGCCTCCAGGACGGCTGCCTCATCTACAACGTCAAGATCAGAGGGGTGAACTTCCCATCCAACGGCCCTGTGATGCAGAAGAAAACACTCGGCTGGGAGGCCAACACCGAGATGCTGTACCCCGCTGACGGCGGCCTGGAAGGCAGAACCGACATGGCCCTGAAGCTCGTGGGCGGGGGCCACCTGATCTGCAACTTCAAGACCACATACAGATCCAAGAAACCCGCTAAGAACCTCAAGATGCCCGGCGTCTACTATGTGGACCACAGACTGGAAAGAATCAAGGAGGCCGACAAAGAGACCTACGTCGAGCAGCACGAGGTGGCTGTGGCCAGATACTGCGACCTCCCTAGCAAACTGGGGCACAAACTTAATGGCATGGACGAGCTGTACAAG

>mRFP1

ATGGCCTCCTCCGAGGACGTCATCAAGGAGTTCATGCGCTTCAAGGTGCGCATGGAGGGCTCCGTGAACGGCCACGAGTTCGAGATCGAGGGCGAGGGCGAGGGCCGCCCCTACGAGGGCACCCAGACCGCCAAGCTGAAGGTGACCAAGGGCGGCCCCCTGCCCTTCGCCTGGGACATCCTGTCCCCTCAGTTCCAGTACGGCTCCAAGGCCTACGTGAAGCACCCCGCCGACATCCCCGACTACTTGAAGCTGTCCTTCCCCGAGGGCTTCAAGTGGGAGCGCGTGATGAACTTCGAGGACGGCGGCGTGGTGACCGTGACCCAGGACTCCTCCCTGCAGGACGGCGAGTTCATCTACAAGGTGAAGCTGCGCGGCACCAACTTCCCCTCCGACGGCCCCGTAATGCAGAAGAAGACCATGGGCTGGGAGGCCTCCACCGAGCGGATGTACCCCGAGGACGGCGCCCTGAAGGGCGAGATCAAGATGAGGCTGAAGCTGAAGGACGGCGGCCACTACGACGCCGAGGTCAAGACCACCTACATGGCCAAGAAGCCCGTGCAGCTGCCCGGCGCCTACAAGACCGACATCAAGCTGGACATCACCTCCCACAACGAGGACTACACCATCGTGGAACAGTACGAGCGCGCCGAGGGCCGCCACTCCACCGGCGCCTAA

4. Under “Step 2-Set your pairwise alignment options”, select the “OUTPUT FORMAT” as “Pair”. Keep other pairwise alignment options using the default settings and click “submit. The job will take a few moments to run, after which the results will automatically load. Values including sequence similarity and identity are provided. Horizontal lines (|) represent matches, where the nucleotides are identical. Periods (.) represent mismatches, where the nucleotides are different. Dashes (-) represent gaps, where a nucleotide is missing.

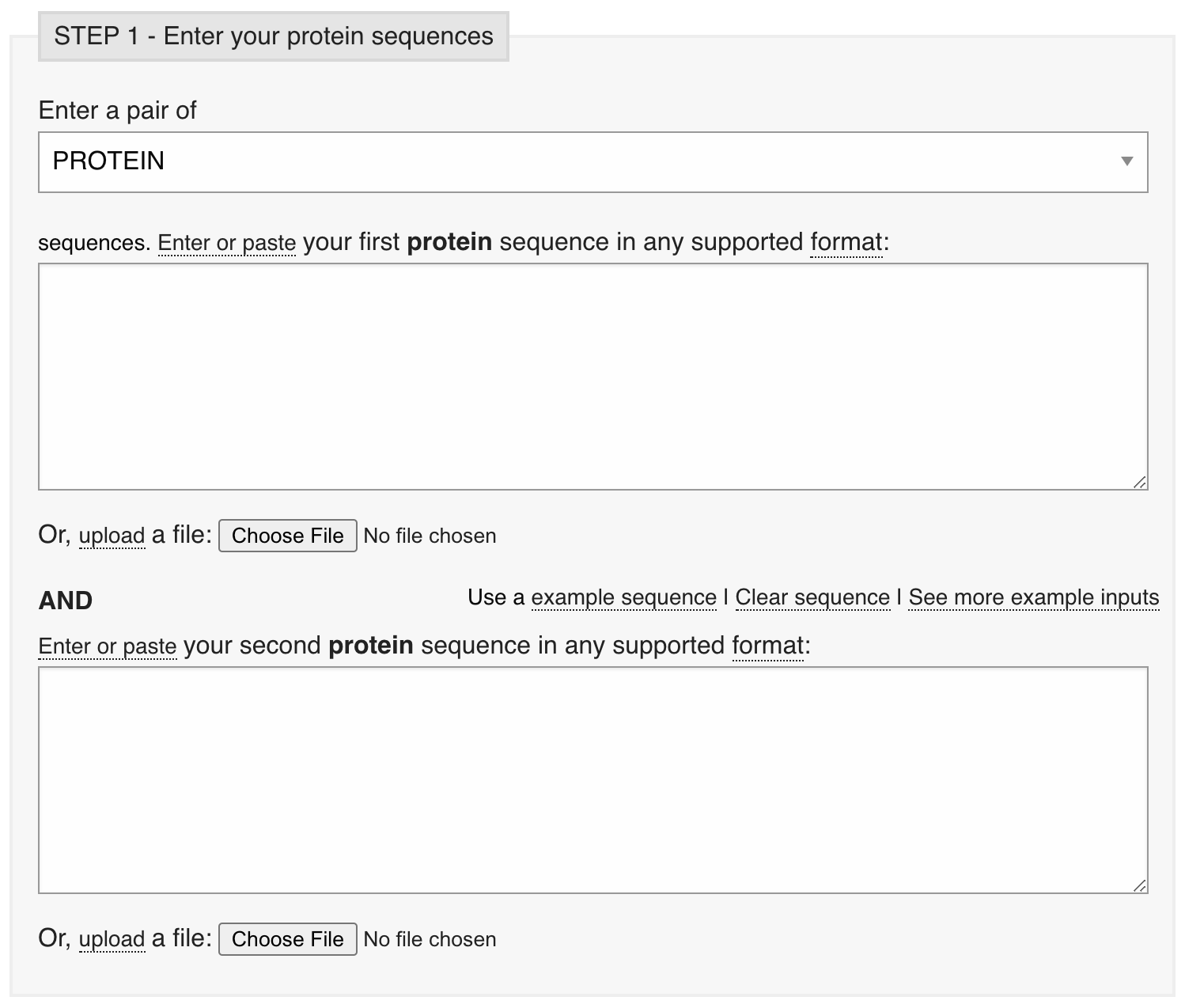

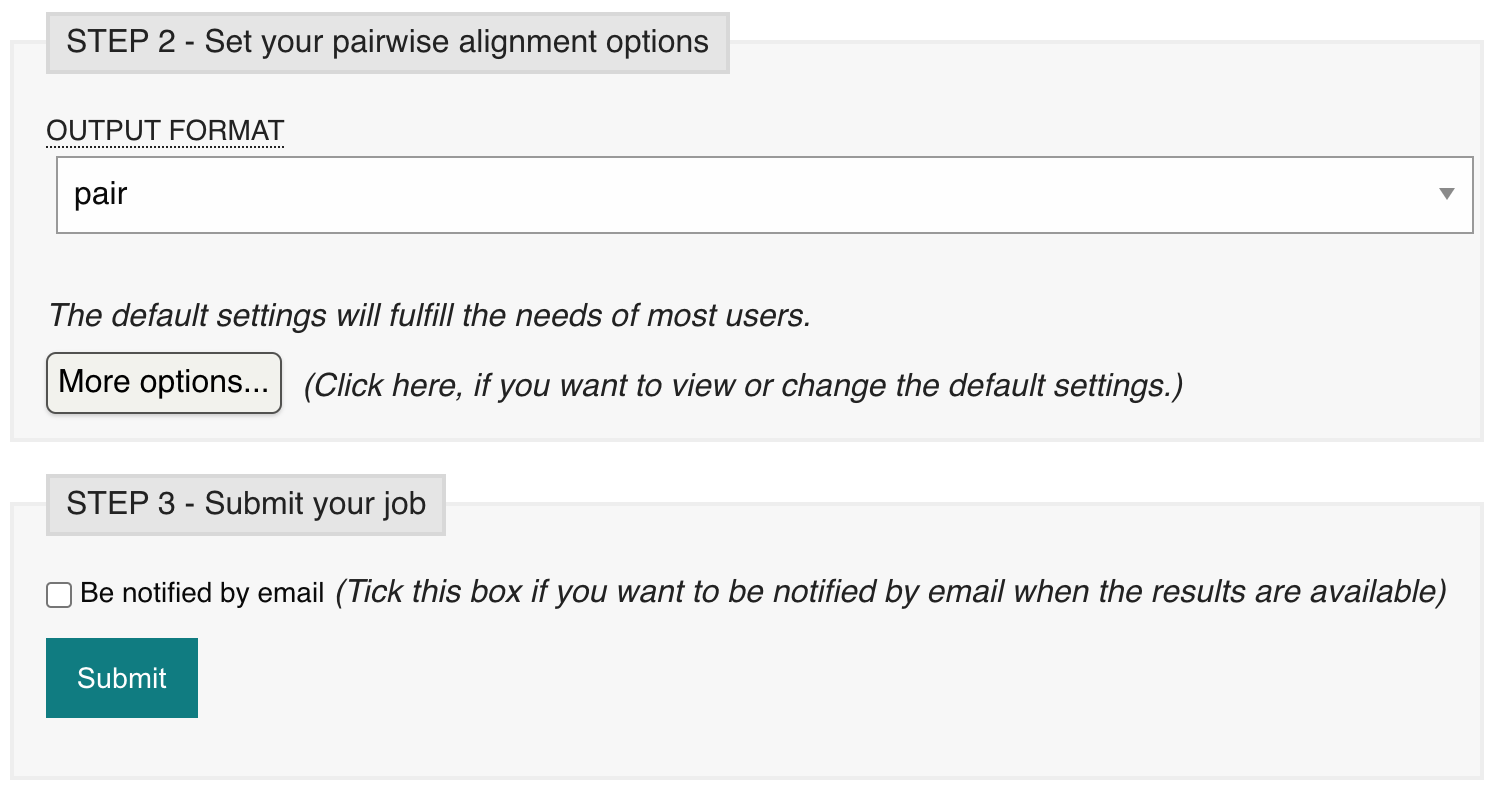

*Questions for students (Bloom’s taxonomy level: Applying)*:

1. Would you expect two closely related organisms to have more or less sequence similarity than two distantly related organisms? Why?

   *Answer:* The more closely related two organisms are, the less time has passed since their divergence from a common ancestor. As a result of less time passing, fewer mutations would have accrued that would lead to substantial changes in their DNA sequences.
2. Why might a 1 base pair gap be more detrimental than a 3 base pair gap?

*Answer:* Since codons represent a group of 3 nucleotides, a 1 base pair gap or “deletion” will disrupt the codon reading frame, resulting in a frameshift. This frameshift disrupts the original codon sequence thereby altering the original amino acid sequence code. On the other hand, a 3 base pair gap would maintain the original reading frame, with the exception of the 3 base pairs originally deleted, since codons are a group of 3 nucleotides. The ultimate consequence of a 3 base pair gap would be a loss of one amino acid.

**Phase II**

After 30-60 minutes, new groups of 3 will be formed, consisting of one “expert” from each prior activity/category. In these new groups, each student is responsible for teaching his or her expertise to each other student in their new group. The students are then assigned the accompanying post-activity assessment (see below) to complete individually to determine if adequate instruction and explanation was given by peers. The post-activity assessment should take no longer than 30 minutes to complete. After 30 minutes, instructors should review the answers of the post-activity assessment with students.

**Name: __________________________________________________	Post-activity assessment**

Base your answers to questions 1-7 below on the following DNA sequence of the *C. elegans* gene *sqt-1*:

ATGTCTGTAAAACTTGCGTGTTATGTGACGGCTTCGGTCACTGTCGCCACTCTTATGGTATGCTTCATGACCATGTCAACCATCTACTCGGAGGTTGATGGATTCAGAGAGAAGCTTGATACCGAGATGAATGTGTTCAGAgtgagttaaattatattttgaattttaaataatttttaattttctagCAATCTACCAATGGATTGTGGAAGGACATAGTTGTCATCGGAAGATCTAGCAAGCGTGTCCGTCGTCAATATGAAGAGACCAACGCTACCCCAACTCCACATGCTGATGGATCCCCATCTGCTCCACCAGGTCAACCACCAGCAGTTCCACCAGTCTTCAACCAGCCAAAGACTCCAAATGGAGCCAATGGAAATGGACCAACCTGCAACTGCAATGCTGATAACAAGTGCCCAGCTGGACCATCCGGACCAAAGGGAGTTCCAGGAGTTCCAGGACTCGACGGAGTTCCAGGACTTGACGGTGTTCCAGGAGTTGGAGCTGATGATATCGCTCCACAACGCGAGTCTGTCGGATGCTTCACTTGCCCACAAGGACCAGTTGGACCACCAGGAGCTCTTGGAAGACCAGGACCACGTGGACTTCCAGGACCAAGAGGACAAAATGGAAACCCAGGAAGAGATGGACAACCAGGACATCCAGGAGAGCAAGGATCATCCGGTCAAATCGGAAAGATCGGAGAGCCAGGACCACCAGGAGAGAAGGGACGCGACGCCGAGCATCCAATCGGAAGACCAGGACCAAAGGGACCAAGAGGAGATCAAGGACCAACAGGACCAGCTGGACAGAACGGTCTTCACGGACCACCAGGAGAGCCAGGAACCGTTGGACCAGAAGGACCATCTGGAAAGCAAGGACGTCAAGGACCAGACGGAACCCAGGGAGAGACTGGACCAGACGGAAGACCAGGAAAGGATGCCGAGTACTGCCAGTGCCCAGACAAGTCTCCACCATCAGAGGCTGTCAACGCCAACCGTGGATACAGAAATATCTAAattgttggtgttttctaataaaatatttgagattc

1. Select the following option that correctly matches the term to the flow of information:
   A. Transcription: RNA → DNA; Translation: DNA → protein
   B. Transcription: DNA → RNA; Translation: RNA → protein
   C. Translation: RNA → DNA; Translation: DNA → protein
   D. Translation: DNA → RNA; Transcription: RNA → protein
2. Underline the start codon
3. Circle the stop codon.
4. What are the first 3 nucleotides of the second exon?

___ ___ ___

5. What are the first 3 nucleotides of the first intron?

___ ___ ___

6. Write down the sequence of the third codon. Then transcribe and translate it.

___ ___ ___	→	___ ___ ___	→	___________________

7. Based on the **Codon Usage in *C. elegans*** table, is the sequence you identified above the most optimal codon for expression in *C. elegans*? (Circle) If NO, write down the codon that is the most optimal.
   A. YES
   B. NO ... ___ ___ ___

**Name: __________________________________________________	Post-activity assessment**

Base your answers to questions 8-15 on the sequence alignment below:

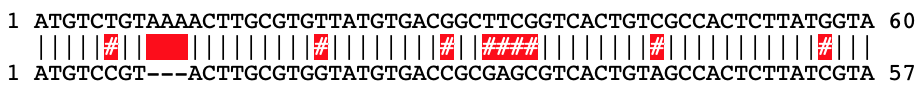

8. How many mismatches (in nucleotides) are there between the 2 sequences being compared?
   ______	nucleotides
9. How many gaps (in nucleotides) are there between the 2 sequences being compared?
   ______	nucleotides
10. What percent similarity does the sample sequence (bottom) share with the reference sequence (top)?

______	nucleotides	÷	60 nucleotides	=	______	%

11. Do the mutations in the bottom sequence disrupt the open reading frame? Why or why not?

YES NO
__________________________________________________________________________________

12. Which codon pairs in the sequences compared above no longer produce the same amino acid? (Circle)
    A. TGT; TGG
    B. ACG; ACC
    C. GCT; GCG
    D. TCG; AGC
13. Based on your answer to question 12, which amino acid does the codon in the reference (top) sequence code for?
    ___________________
14. Based on your answer to question 12, which amino acid does the codon in the sample (bottom) sequence code for?
    ___________________
15. What other codon would code for the amino acid you listed in question 13?
    ___________________

**Answer key:**

1. B
2. ATG (first 3 letters)
3. TAA (last 3 capitalized letters)
4. CAA
5. gtg
6. GTA → GUA → Valine/Val/V
7. No; GUC (GTC also acceptable)
8. 9
9. 3
10. 48; 80%
11. No; the deletion is a multiple of 3
12. A
13. Cysteine/Cys/C
14. Tryptophan/Trp/W
15. TGC/UGC

**Table 1: RNA codon table**

| **1st**  **base** | **2nd base** | | | | | | | | **3rd**  **base** |
| --- | --- | --- | --- | --- | --- | --- | --- | --- | --- |
|  | **U** | | **C** | | **A** | | **G** | |  |
| **U** | UUU | Phenylalanine  (Phe/F) | UCU | Serine  (Ser/S) | UAU | Tyrosine  (Tyr/Y) | UGU | Cysteine  (Cys/C) | **U** |
|  | UUC |  | UCC |  | UAC |  | UGC |  | **C** |
|  | UUA | Leucine  (Leu/L) | UCA |  | UAA | STOP | UGA | STOP | **A** |
|  | UUG |  | UCG |  | UAG | STOP | UGG | Tryptophan  (Trp/W) | **G** |
| **C** | CUU |  | CCU | Proline  (Pro/P) | CAU | Histidine  (His/H) | CGU | Arginine  (Arg/R) | **U** |
|  | CUC |  | CCC |  | CAC |  | CGC |  | **C** |
|  | CUA |  | CCA |  | CAA | Glutamine  (Gln/Q) | CGA |  | **A** |
|  | CUG |  | CCG |  | CAG |  | CGG |  | **G** |
| **A** | AUU | Isoleucine  (Ile/I) | ACU | Threonine  (Thr/T) | AAU | Asparagine  (Asn/N) | AGU | Serine  (Ser/S) | **U** |
|  | AUC |  | ACC |  | AAC |  | AGC |  | **C** |
|  | AUA |  | ACA |  | AAA | Lysine  (Lys/K) | AGA | Arginine  (Arg/R) | **A** |
|  | AUG | Methionine  (Met/M) | ACG |  | AAG |  | AGG |  | **G** |
| **G** | GUU | Valine  (Val/V) | GCU | Alanine  (Ala/A) | GAU | Aspartic acid  (Asp/D) | GGU | Glycine  (Gly/G) | **U** |
|  | GUC |  | GCC |  | GAC |  | GGC |  | **C** |
|  | GUA |  | GCA |  | GAA | Glutamic acid  (Glu/E) | GGA |  | **A** |
|  | GUG |  | GCG |  | GAG |  | GGG |  | **G** |

**Table 2: Codon Usage in *C. elegans***

| **AA** | **Codon** | **N** | **RSCU** |
| --- | --- | --- | --- |
| Ala | GCA | 537 | 0.35 |
|  | GCC* | 2631 | 1.72 |
|  | GCG | 171 | 0.11 |
|  | GCU* | 2777 | 1.82 |
| Arg | AGA | 906 | 1.45 |
|  | AGG | 43 | 0.07 |
|  | CGA | 168 | 0.27 |
|  | CGC* | 935 | 1.49 |
|  | CGG | 39 | 0.06 |
|  | CGU* | 1665 | 2.66 |
| Asn | AAC* | 2237 | 1.51 |
|  | AAU | 726 | 0.49 |
| Asp | GAC* | 1991 | 0.99 |
|  | GAU | 2046 | 1.01 |
| Cys | UGC* | 895 | 1.47 |
|  | UGU | 321 | 0.53 |
| Gln | CAA | 2363 | 1.35 |
|  | CAG* | 1147 | 0.65 |
| Glu | GAA | 2110 | 0.79 |
|  | GAG* | 3255 | 1.21 |
| Gly | GGA* | 5807 | 3.29 |
|  | GGC | 351 | 0.2 |
|  | GGG | 108 | 0.06 |
|  | GGU | 797 | 0.45 |
| His | CAC* | 935 | 1.32 |
|  | CAU | 479 | 0.68 |
| Ile | AUA | 40 | 0.03 |
|  | AUC* | 2483 | 2.01 |
|  | AUU | 1179 | 0.96 |
| Leu | CUA | 66 | 0.07 |
|  | CUC* | 2066 | 2.28 |
|  | CUG | 268 | 0.3 |
|  | CUU* | 2052 | 2.26 |
|  | UUA | 49 | 0.05 |
|  | UUG | 946 | 1.04 |

| **AA** | **Codon** | **N** | **RSCU** |
| --- | --- | --- | --- |
| Lys | AAA | 781 | 0.3 |
|  | AAG* | 4373 | 1.7 |
| Met | AUG | 1473 | – |
| Phe | UUC* | 2139 | 1.74 |
|  | UUU | 315 | 0.26 |
| Pro | CCA* | 3994 | 3.6 |
|  | CCC | 91 | 0.08 |
|  | CCG | 192 | 0.17 |
|  | CCU | 166 | 0.15 |
| Ser | AGC | 409 | 0.61 |
|  | AGU | 148 | 0.22 |
|  | UCA | 517 | 0.77 |
|  | UCC* | 1400 | 2.08 |
|  | UCG | 466 | 0.69 |
|  | UCU* | 1104 | 1.64 |
| STOP | UAA | 137 | 2.31 |
|  | UAG | 27 | 0.46 |
|  | UGA | 14 | 0.24 |
| Thr | ACA | 410 | 0.44 |
|  | ACC* | 1990 | 2.14 |
|  | ACG | 158 | 0.17 |
|  | ACU | 1167 | 1.25 |
| Trp | UGG | 591 | – |
| Tyr | UAC* | 1523 | 1.56 |
|  | UAU | 432 | 0.44 |
| Val | GUA | 215 | 0.19 |
|  | GUC* | 2057 | 1.81 |
|  | GUG | 564 | 0.49 |
|  | GUU | 1722 | 1.51 |

AA: amino acid

N: raw codon usage values (in genes exhibiting high bias)

RSCU: relative synonymous codon usage

*Optimal codons

Table adapted from Sharp & Bradnam (1997)

1.	Sharp PM, Li WH. 1987. The codon Adaptation Index--a measure of directional synonymous codon usage bias, and its potential applications. Nucleic Acids Res 15:1281-95.
