## Supplemental File 14 for "A laboratory module that explores RNA interference and codon optimization through fluorescence microscopy using *Caenorhabditis elegans*"

**S14. A Laboratory Module-Common Student Misconceptions and Questions**

### **Some Students’ misconceptions and difficulty grasping concepts:**

- Believing that RNAi knockdown of GFP in the GFP_NCO_ strain makes the genetic background of the strain susceptible to all RNAi – regardless of the sequence
- Trouble grasping the purpose of the experiment
- Thinking that GFP has a role in regulating the mean fluorescence intensity rather than the amount of protein produced
- Concluding that the GFP_NCO_ strain targets GFP whereas the GFP_CO_ strain can not target GFP as efficiently
- Assuming that the dsRNA is delivering the GFP gene to both of the strains and the experiment is testing to see which strain more readily accepts the gene

### **Common student mistakes during the module:**

- Incorrect or missing scale bars
- Students did not perform statistical tests, did not properly label their plots to display the results of the statistical test, or did not include the statistical test type used
- Failed to specify what error bars represent (i.e. standard deviation)
- Failed to specify sample size (n)
- Significance not matching the standard p-value convention
- Students did not correctly label x and y axes
- Incorrect thresholding of micrographs (image(s) qualitatively significantly brighter/dimmer than it/they should be)
- Illegible figure fonts
- Not all Images were oriented in the same direction
- Students failed to clearly state hypothesis
- Students did not correctly resize images during figure making ( i.e. stretching out the figure rather than properly scaling the figure)
- Lacked consistent levels of brightness/contrast between images within the same strain and conditions
- Provide figure legends that were too descriptive (i.e. figure legends summarize results rather than describe the details of what is being shown in the figure)
- Improper *C. elegans* nomenclature (protein names should be capitalized and not in italics, while gene names are in lower case and italicized)
- Students fail to normalize their data

**Common student questions centered around using ImageJ:**

**Question:** How do you adjust the contrast and brightness on a figure?

**Answer:** Open FIJI/ImageJ>Image>Adjust>Contrast/Brightness

**Question:** How do you measure full body “mean fluorescent intensity values”?

**Answer:** Select “freehand selections” from the toolbar and trace the body of the worm as best as you can using the DIC channel to help. Once the worm body is traced, click the fluorescent channel by clicking the drag bar at the bottom of your image. Click the button M on your keyboard (the shortcut for the measure function). Make sure that under Analyze>Set Measurement Mean Gray Value is checked.

**Question:** How do you account for background static in the image when measuring the intensity for the entire animals’ body?

**Answer:** Make a circle on the darkest part of your image and click M. Subtract the background measurement from the full body mean fluorescent intensity measurement number.

**Common student questions centered around RNAi and codon optimization:**

**Question:** How do the *C. elegans* strains receive the dsRNA from RNAi?

**Answer:** dsRNA is introduced to the worms via feeding.

**Question:** How does the dsRNA get into the E. Coli?

**Answer:** We prepare dsRNA from our labs pre-existing RNAi libraries in which the plasmid to produce the dsRNA has already been transformed into the bacteria. We grow *E. Coli* expressing the plasmid overnight at 37°C and use IPTG to induce the expression of RNA polymerase to produce dsRNA. We then plate the bacteria onto fresh NGM plates containing both antibiotic and IPTG to ensure the continual expression of the plasmid and the induction of the RNA polymerase to make the dsRNA and let it dry.

**Question:** Are all cells effected by RNAi?

**Answer:** Most cells are affected with the exception of neurons.

**Question:** Are there any restrictions on genes that can be codon optimized?

**Answer:** No, any gene can be codon optimized for optimal expression in *C. elegans*.

**Question:** At what level (transcription or translation) is codon optimization working?

**Answer:** Codon optimization significantly impacts translation of mRNA into protein given that codon usage and availability of tRNAs is tightly linked.
